# New insights into malaria susceptibility from the genomes of 17,000 individuals from Africa, Asia, and Oceania

**DOI:** 10.1101/535898

**Authors:** Malaria Genomic Epidemiology Network, Gavin Band, Quang Si Le, Geraldine M. Clarke, Katja Kivinen, Christina Hubbart, Anna E. Jeffreys, Kate Rowlands, Ellen M. Leffler, Muminatou Jallow, David J. Conway, Fatoumatta Sisay-Joof, Giorgio Sirugo, Umberto d’Alessandro, Ousmane B. Toure, Mahamadou A. Thera, Salimata Konate, Sibiri Sissoko, Valentina D. Mangano, Edith C. Bougouma, Sodiomon B. Sirima, Lucas N. Amenga-Etego, Anita K. Ghansah, Abraham V. O. Hodgson, Michael D. Wilson, Anthony Enimil, Daniel Ansong, Jennifer Evans, Subulade A. Ademola, Tobias O. Apinjoh, Carolyne M. Ndila, Alphaxard Manjurano, Chris Drakeley, Hugh Reyburn, Nguyen Hoan Phu, Nguyen Thi Ngoc Quyen, Cao Quang Thai, Tran Tinh Hien, Yik Ying Teo, Laurens Manning, Moses Laman, Pascal Michon, Harin Karunajeewa, Peter Siba, Steve Allen, Angela Allen, Melanie Bahlo, Timothy M. E. Davis, Victoria Cornelius, Jennifer Shelton, Chris C.A. Spencer, George B.J. Busby, Angeliki Kerasidou, Eleanor Drury, Jim Stalker, Alexander Dilthey, Alexander J. Mentzer, Gil McVean, Kalifa A. Bojang, Ogobara Doumbo, David Modiano, Kwadwo A. Koram, Tsiri Agbenyega, Olukemi K. Amodu, Eric Achidi, Thomas N. Williams, Kevin Marsh, Eleanor M. Riley, Malcolm Molyneux, Terrie Taylor, Sarah J. Dunstan, Jeremy Farrar, Ivo Mueller, Kirk A. Rockett, Dominic P. Kwiatkowski

## Abstract

We conducted a genome-wide association study of host resistance to severe *Plasmodium falciparum* malaria in over 17,000 individuals from 11 malaria-endemic countries, undertaking a wide ranging analysis which identifies five replicable associations with genome-wide levels of evidence. Our findings include a newly implicated variant on chromosome 6 associated with risk of cerebral malaria, and the discovery of an erythroid-specific transcription start site underlying the association in *ATP2B4*. Previously reported HLA associations cannot be replicated in this dataset. We estimate substantial heritability of severe malaria (*h^2^* ~ 23%), of which around 10% is explained by the currently identified associations. Our dataset will provide a major building block for future research on the genetic determinants of disease in these diverse human populations.

## Introduction

Genome-wide association studies (GWAS) have been very successful in identifying common genetic variants underlying chronic non-communicable diseases, but have proved more difficult for acute infectious diseases that represent a substantial portion of the global disease burden and are most prevalent in tropical regions. This is partly due to the practical difficulties of establishing large sample collections and reliable phenotypic datasets in resource-constrained settings, but also theoretical and methodological challenges associated with the study of pathogenic diseases in populations with high levels of genetic diversity and population structure^3–5^. The Malaria Genomic Epidemiology Network (MalariaGEN) was established in 2005 to overcome these obstacles with standardized protocols, common phenotypic definitions, agreed policies for equitable data sharing, and local capacity building for genetic data analysis, thereby enabling large collaborative studies across different countries where malaria is endemic^6^. Here we extend previous work by using data collected from 11 malaria-endemic populations to perform the most comprehensive genome-wide association study of human resistance to severe malaria to date.

## Results

### A genome-wide dataset representing the diversity of populations from the malaria-endemic belt

We generated genome-wide SNP typing on samples from our previously described case-control study, which includes 16,115 individuals admitted to hospital with severe symptoms of *Plasmodium falciparum* malaria, and 20,229 population controls^7^. In brief, individuals were sampled from 12 study sites, including ten in sub-Saharan Africa and the others in Vietnam and Papua New Guinea (**Table S1**). Severe malaria (SM) was ascertained according to the WHO criteria^8^, and represents a heterogeneous phenotype including diagnoses of cerebral malaria (CM), severe malarial anaemia (SMA), and other malaria-related symptoms (here referred to as other SM). For our GWAS discovery phase, a subset of samples, preferentially chosen for phenotype severity and DNA quality, were genotyped on the Illumina Omni 2.5M platform. We jointly processed these samples to produce a single set of estimated haplotypes for 17,960 individuals at the set of over 1.5M SNPs genome-wide that passed our quality control process (**Methods**). This dataset includes 6,888 individuals from Mali, Burkina Faso, Ghana, Nigeria, Cameroon, Tanzania, Vietnam and Papua New Guinea that have not previously been included in meta-analysis, as well as previously reported data from The Gambia, Malawi and Kenya ^9,10^, and thus reflects the haplotype diversity of a substantial portion of the malaria-endemic world.

### A reference panel enriched for African DNA improves imputation accuracy across the genome

The ethnically diverse nature of our study provides challenges for genomic inference, including for our ability to impute genotypes at potentially relevant untyped loci^11^. To address this, we sequenced the genomes of 773 individuals from ten ethnic groups in east and west Africa (specifically from the Gambia, Burkina Faso, Cameroon and Tanzania), including 207 family trios (**Figure 1**). We combined genotypes at single nucleotide polymorphisms (SNPs) in these data with Phase 3 of the 1000 Genomes Project to form an imputation reference panel which covers most common genetic variation^12^ and in which two-fifths of the donor families are of African ancestry (1,203 of 3,046 individuals). In principle, the additional representation of African DNA in this panel should lead to improvements in imputation accuracy for African study populations, and we found that this was indeed the case, with use of our panel leading to a large increase in accuracy relative to panels used in our previous GWAS^9–11^ and a more modest improvement relative to using the 1000 Genomes Phase 3 panel alone (**Figure S1**). Imputation of Vietnamese individuals, and those from Papua New Guinea (PNG) which is substantially diverged from any reference panel population (**Figure 1d**), were less affected by the inclusion of these additional haplotypes.

**Figure 1:**
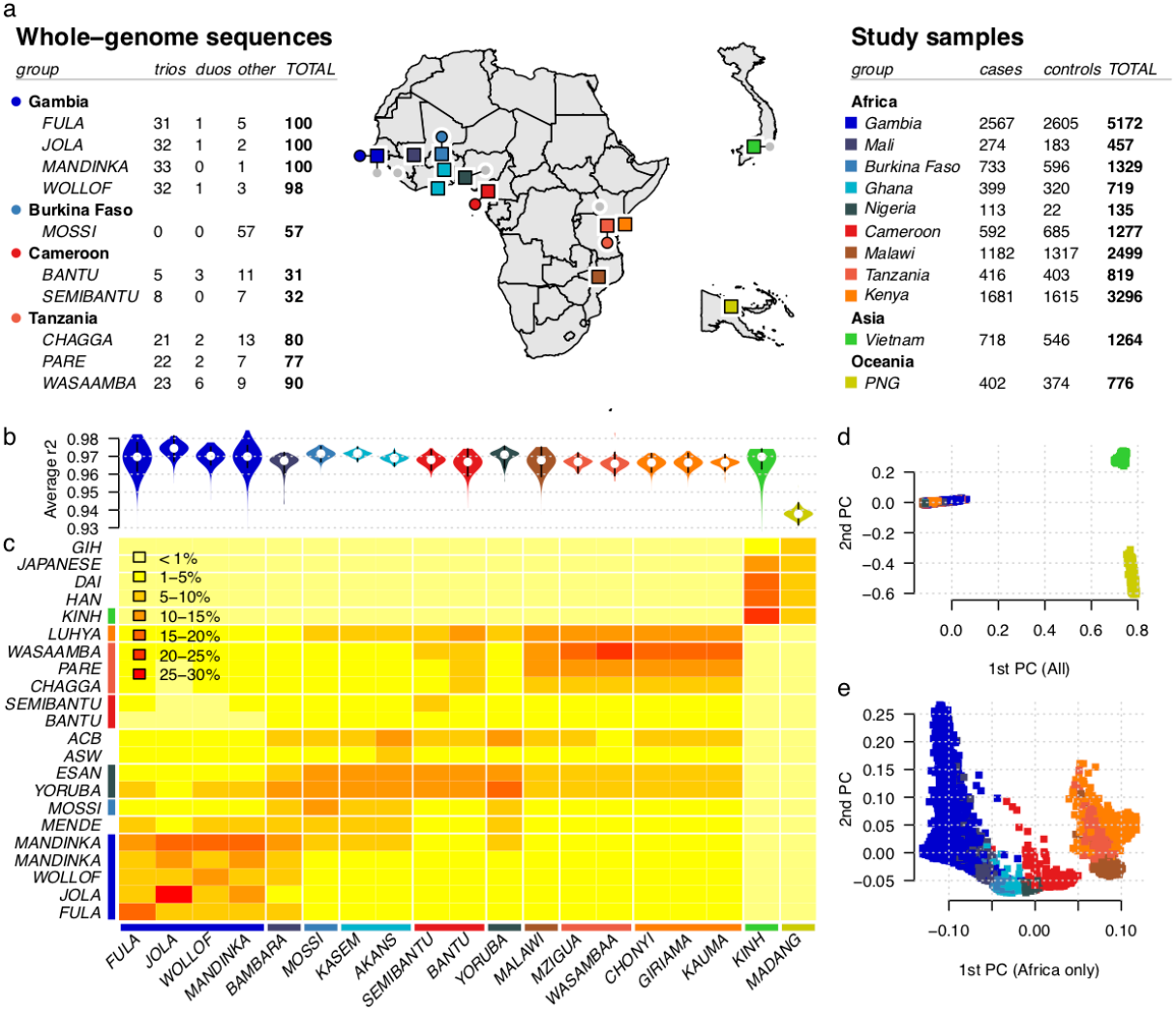
Overview of datasets and imputation performance. **a)** Counts of whole-genome sequenced samples (reference panel samples, left table), samples typed on the Omni 2.5M platform (study samples, right table), and geographic locations of sampling (map). Counts reflect numbers of samples following our quality control process. Sequenced samples were collected in family trios except in Burkina Faso, as shown. Colours shown in tables and map denote country of origin of reference panel (circles) and study samples (squares), with small grey circles indicating 1000 Genomes Project populations. **b)** Imputation performance, measured as the mean squared correlation between directly typed and re-imputed variants for each sample. **c)** Distribution of most similar haplotypes. For each GWAS sample, the average number of 1Mb chunks such that the most similar haplotype lies in the given reference panel population (y axis) is shown. Values are averaged over samples within each GWAS population (x axis). **d, e)** Principal components (PCs) computed across all 17,120 study samples, or the subset of 15,152 samples of African ancestry, after removing samples with close relationships.

We specifically examined imputation of malaria-protective alleles in the *HBB* gene, which have previously been found difficult to impute^9,11^. The SNP encoding the sickle cell mutation (rs334, chr11:5248232) was imputed with r>0.9 in all African populations, as compared with genotypes obtained through direct typing^7^. We did note that some potentially relevant loci still appear not to be accessible through these data, including the common deletion of *HBA1-HBA2* that causes alpha-thalassaemia (**Text S1**) and the region around the gene encoding the invasion receptor basigin^13^ (**Text S2**). Thus, although overall accuracy is high across the genome, additional work will be needed to access a smaller set of complex but potentially relevant regions.

### Association testing reveals a novel association on chromosome 6

We used the imputed genotypes to test for association with severe malaria, and with severe malaria subtypes, at over 15 million SNPs and indels genome-wide. Specifically, association tests were conducted within each population using logistic and multinomial logistic regression (implemented in SNPTEST; see **Methods**), including principal components to control for population structure, and we computed a fixed-effect meta-analysis summary of association across populations (**Figure S3**). In view of the complexities observed at known association signals ^9,11,14^, we also extended our previously described meta-analysis method ^7,10^ to the larger set of populations, phenotypes and variants considered here. This analysis produces an overall measure of evidence (the model-averaged Bayes factor, *BF_avg_*; **Figure 2a**) that is sensitive to more complex patterns of genetic effect than are detected by frequentist fixed-effect analysis (**Figure 2b**). The *BF_avg_* is computed under specific prior weights that are detailed in Methods. In particular, *BF_avg_* captures effects that are non-additive, as well as effects that are restricted to particular subphenotypes or population subgroups.

**Figure 2:**
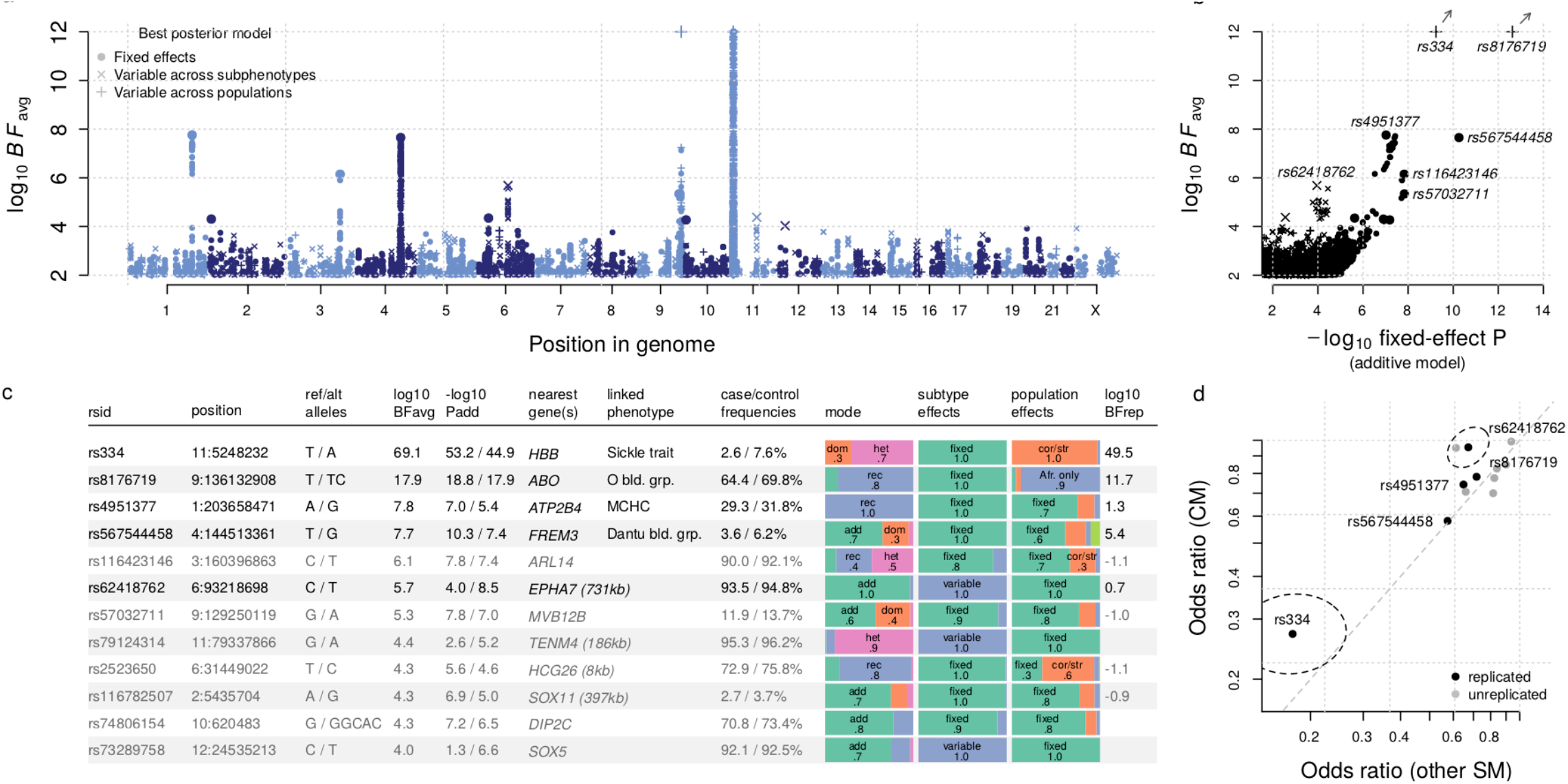
Evidence for association with severe malaria. **a)**: manhattan plot showing typed and imputed SNPs and indels genome-wide. Evidence is assessed under a range of models and summarized in the model averaged Bayes factor (log10 *BF_avg_*, y axis; values are clamped to a maximum value of 12) using prior weights specified in Methods. Shapes reflect whether the effect size for the model with the highest posterior weight is fixed across populations and subphenotypes (case-control effect,•), or suggests variation in effect between populations (x) or between sub-phenotypes (+). **b)** Comparison of model-averaged Bayes factor (log10 *BF_avg_*, y axis, as in **a)** and the evidence under an additive model of association with overall SM (-log10 *P_add_*, x axis). For visualization purposes we have removed variants in the region of rs334 (HbS, chromosome 11) and rs567544458 (glycophorin region, chromosome 4) except the lead variant. Shapes reflect the highest posterior model as in a. The values for rs334 and rs8176719 lie outside the plot as indicated by arrows; to visualize these we have projected toward the origin onto the plot boundary. **c)** Twelve regions of the genome with *BF_avg_* > 10,000, in decreasing order of evidence. Columns reflect the ID, genomic position, alleles, *BF_avg_*, P-value computed under an additive model of association with SM status or with SM subphenotypes, nearest gene and distance to nearest gene for intergenic variants, known linked phenotypes, and combined frequencies in study cases and controls. Bar plots summarise our inference about the mode of effect, and distribution of effects between SM subtypes and between populations. The last column reflects the evidence for association observed in replication samples (log10 *BF_rep_*), assessed using the effect size distribution learnt from discovery samples. Where not directly typed, a tag SNP was used to assess replication evidence as detailed in **Table S2**. Rows are in bold if they showed positive replication evidence (*BF_rep_* > 1). **d)** Comparison of estimated effect sizes on CM (y axis) and on unspecified SM cases (x axis) for the 12 variants in panel **c**. The 95% confidence region for rs334 and rs62418762 (dashed ellipses) are shown.

Several regions of the genome showed evidence for association in this analysis, including seven regions with *BF_avg_* > 1×10^5^, twelve with *BF_avg_* > 1×10^4^ (**Figure 2c**), and 95 with *BF_avg_ >* 1,000. Among these, our data convincingly replicate previously confirmed associations at *HBB, ABO, ATP2B4*, and in the glycophorin region on chromosome 4, while an additional previously reported variant in *ARL14* also shows stronger evidence in this larger sample ^10^. Both a direct interpretation of the Bayes factor, as well as false discovery rate methods suggest a relatively small number of our top signals represent real associations (e.g. roughly 5-9 associations among the top 95 given plausible prior odds^15^ of 10^−6^ to 10^−5^; 5 regions meeting FDR<5% based on the multinomial test *P_add_*). Compatible with this, we found evidence of replication for one newly identified locus (rs62418762; *BF_avg_*=4.8×10^5^; *BF_repiication_* = 5; replication *P_additive_*=0.02 for an additive risk effect of the derived allele on CM and SMA; combined *BF*=2.4×10^6^; combined *P_add_* = 6.7×10^−10^; **Figure 3**), in addition to the four previously confirmed associations, from among 26 regions that we attempted to replicate using direct typing in the additional set of 15,634 individuals not included in our GWAS (**Table S2**).

**Figure 3:**
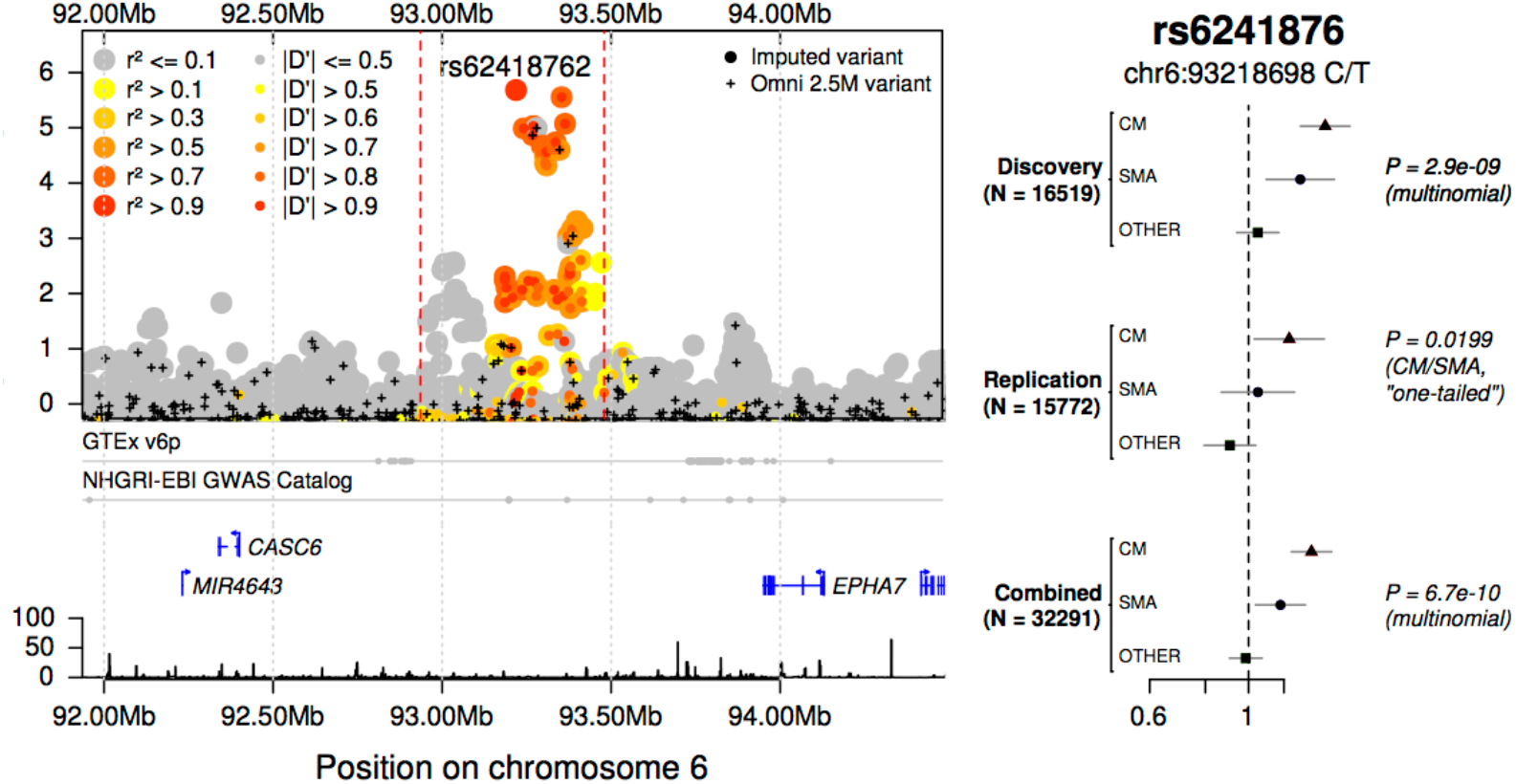
Evidence for association at rs62418762. **a)** Regional hitplot showing evidence for association (log10 *BF_avg_*, y axis) across a 2.5Mb region surrounding rs62418762 (x axis). Points are coloured by LD with rs62418762, estimated using African reference panel haplotypes. Directly-typed SNPs included in the phased dataset are denoted by black plusses. Below, the locations of significant tissue-specific eQTLs, previously identified association signals, regional genes and pseudogenes, and the Hapmap combined recombination rate map are annotated. **b)** Detail of discovery and replication evidence for association at rs62418762 under an additive model. Points and lines represent the estimated odds ratio of the non-reference (T) allele on severe malaria subtypes. Estimates are obtained using multinomial logistic regression in each population and combined across populations using fixed-effect meta-analysis. Top: effect sizes estimated from imputed genotypes in discovery samples. A P-value against the null that all three effect sizes are zero is given. Middle: effect sizes estimated from direct typing of rs62418762 in replication samples. P-value reflects the alternative hypothesis that CM and SMA effects are nonzero and in the direction observed in discovery. Bottom: meta-analysis of discovery and replication results.

Our estimates suggest rs62418762 has a relatively strong effect on cerebral malaria, but less so on other malaria subtypes (**Figure 3b**). rs62418762 lies in a region of chromosome 6 between *MAP3K7* and the nearest gene *EPHA7*, which encodes Ephrin type-A receptor 7, a regulator of neurodevelopment and neural cell adhesion ^16^. Other type A ephrin receptors have been implicated in liver-stage malaria ^17^. However, rs62418762 lies over 700kb distant from *EPHA7* and it is not immediately clear whether any functional link between them exists. A number of non protein-coding transcripts lie nearby (e.g. the pseudogene *ATF1P1*), as do reported signals of association with neurodegenerative disorders^18,19^, and these may provide clues as to the underlying mechanism.

### Genome-wide data suggests a residual polygenic contribution to malaria risk

We used published methods to estimate the heritability of severe malaria captured by genome-wide genotypes (e.g. heritability explained by SNPs on the genotyping array (h^2^)= 0.23, 95% CI = 0.16 − 0.30; computed across African populations using PCGC^20^ assuming a 1% population prevalence of severe malaria; **Figure S4** and **Table S3**). The *HBB, ABO*, glycophorin and *ATP2B4* loci appear to contribute around 11% of this total (i.e. around 2.5% of liability scale phenotype variation), and we noted a trend for the remaining heritability to concentrate near protein-coding genes and at lower-frequency variants (**Figure S4d**). A number of caveats apply to these estimates, which assume a particular relationship between effect size, allele frequency and the degree of LD surrounding causal variants, all of which may be distorted by effects of natural selection ^21,22^, and may further be affected by unmeasured environmental confounders. Nevertheless, our results are comparable to previously reported estimates from family-based studies in east Africa^23^, and suggest that further susceptibility loci will be discoverable with additional data. To assist researchers to integrate our data with other sources of information, including future genetic association studies and functional experiments, the raw and imputed data as well as a full set of results from our analysis are being made available (Data availability) and can be found via the MalariaGEN website (http://www.malariagen.net).

### Joint analysis of associations reveals substantial heterogeneity

We examined the nature of association at replicated loci in more detail (**Figure 2 and Figure 4**), revealing considerable heterogeneity in mode of effect (**Figure 2c**) and across populations and subphenotypes. The sickle haemoglobin allele (HbS, encoded by rs334) is present in all African populations studied, but varies considerably in estimated effects, with greater than five-fold difference between the strongest estimate (OR=0.10 in the Gambia) and the weakest (OR=0.5 in Cameroon). This does not seem to be explained by the presence of the competing HbC mutation in west African populations (**Figure 4**), which is at low frequency in Cameroon. The O blood group-encoding mutation rs8176719 is common in all populations but, in our analysis, appears to have the greatest protective effect in African populations, with an opposite direction of effect observed in Papua New Guinea, an unexpected feature which is also observed in replication samples (**Figure S5**) and could point to hitherto unexplained subtleties in the mechanism of protection of this allele ^24,25^. The structural variant DUP4, which encodes the Dantu NE blood group phenotype ^14^ and is in LD with rs567544458 in our genome-wide imputation (**Fig 2c**), is essentially absent from study populations outside east Africa (f < 0.5% except in Malawi, Tanzania and Kenya). Finally, our data demonstrate differences in effect size between malaria subphenotypes, with all of the replicating variants displaying smaller effects in nonspecific cases of severe malaria compared to cerebral or severely anaemic cases, likely reflecting the mixed phenotypic composition of this set of samples (**Figure 2d** and **Figure 4a**). Inclusion of these variants in a joint model suggests they act largely independently in determining malaria risk (**Figure 4c**).

**Figure 4:**
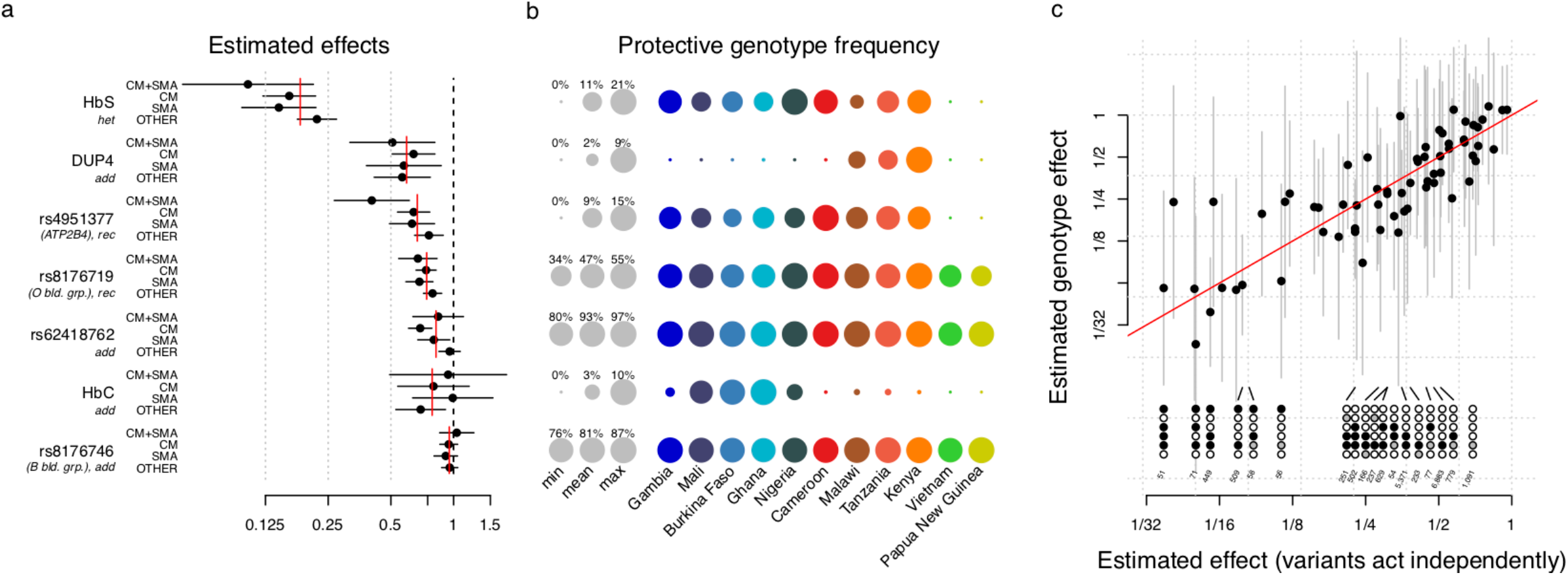
A joint model for natural genetic resistance to malaria. **a)** Effect sizes for severe malaria subtypes are estimated in a joint model which includes the five replicating associated variants and two additional variants (rs33930165, which encodes haemoglobin C, and rs8176746 which reflects the A/B blood group) in associated regions. The model was fit across all eleven populations assuming the effect on each phenotype is fixed across populations, and including a population indicator and five principal components in each population as covariates. Each variant is encoded according to the mode of inheritance of the protective allele inferred from discovery analysis. Red lines indicate the overall effect across severe malaria subtypes, computed as an inverse variance-weighted mean of the per-phenotype estimates. **b)** The frequency of the protective allele (for effects inferred as additive) or protective genotype (for non-additive effects) of each variant in each population. Grey circles depict the minimum, mean, and maximum observed frequencies across populations. Coloured circles reflect the per-population frequencies. Frequency estimates are computed using control samples only. **c)** Comparison of effect size estimates for combinations of genotypes (stacked circles) carried by at least 50 study individuals, across the top 6 variants in panel a. Black filled and open circles denote the protective and risk dosage at the corresponding variant; grey circles denote carrying a heterozygote dosage at effects inferred to be additive. Effect size estimates are computed using the model in panel a) assuming independent effects (x axis), or jointly allowing each genotype its own effect (y axis).

### Functional annotation across cell types reveals an erythroid-specific transcription start site in *ATP2B4*

We reasoned that functional annotations might provide clues to further putatively causal variants among our list of most associated regions. To assess this, we annotated all imputed variants with information indicative of functional importance, including location and predicted function within genes^26^, chromatin state ^27–29^ and transcriptional activity ^28,30,31^ across cell types, status as an eQTL ^31–33^, and association evidence from other traits^34–37^. This analysis revealed a number of potentially functional variants with evidence of association (**Table S4**).

We uncovered compelling evidence that the association in *ATP2B4*, which encodes the plasma membrane calcium pump PMCA4, is driven by an alternative transcription start site (TSS) that is specific to erythroid cells (**Figure 5**). Specifically, we found that associated SNPs overlap a binding site for the GATA1 transcription factor, an important regulator of expression in erythroid cells ^27,29,38^, in the first intron of *ATP2B4*. The derived allele at one of these SNPs (rs10751451) disrupts a GATA motif^31^, and the same SNPs have previously been shown to associate with *ATP2B4* expression levels in whole blood ^33^, in experimentally differentiated erythrocyte precursors ^31^, and with PMCA4 levels in circulating RBCs ^39^. We noted that the GATA1 site lies just upstream of an exon that is not listed in the GENCODE ^40^, RefSeq ^41^, or ENSEMBL ^42^ transcript models, but that can be found in FANTOM5 ^43^ (Figure 5b). Inspection of RNA-seq data revealed that this forms the first exon of the main *ATP2B4* transcript expressed by erythroid cells^29,31,44^, as well as by the K562 cell line, but appears not to be expressed in other cell types ^27,28,32^ (**Figure 5a**). Moreover, the derived allele, which is associated with malaria protection, is correlated with decreased expression of this alternative first exon (P=0.01, using data from 24 fetal and adult erythroblasts^31^; **Figure 5g**) and all subsequent exons (P<0.03) of *ATP2B4*, but not with the annotated first exon (P=0.25). These results indicate that the malaria-associated SNPs affect *ATP2B4* expression in a erythroid-specific manner, by affecting the promoter of a TSS that is only active in these cells.

**Figure 5:**
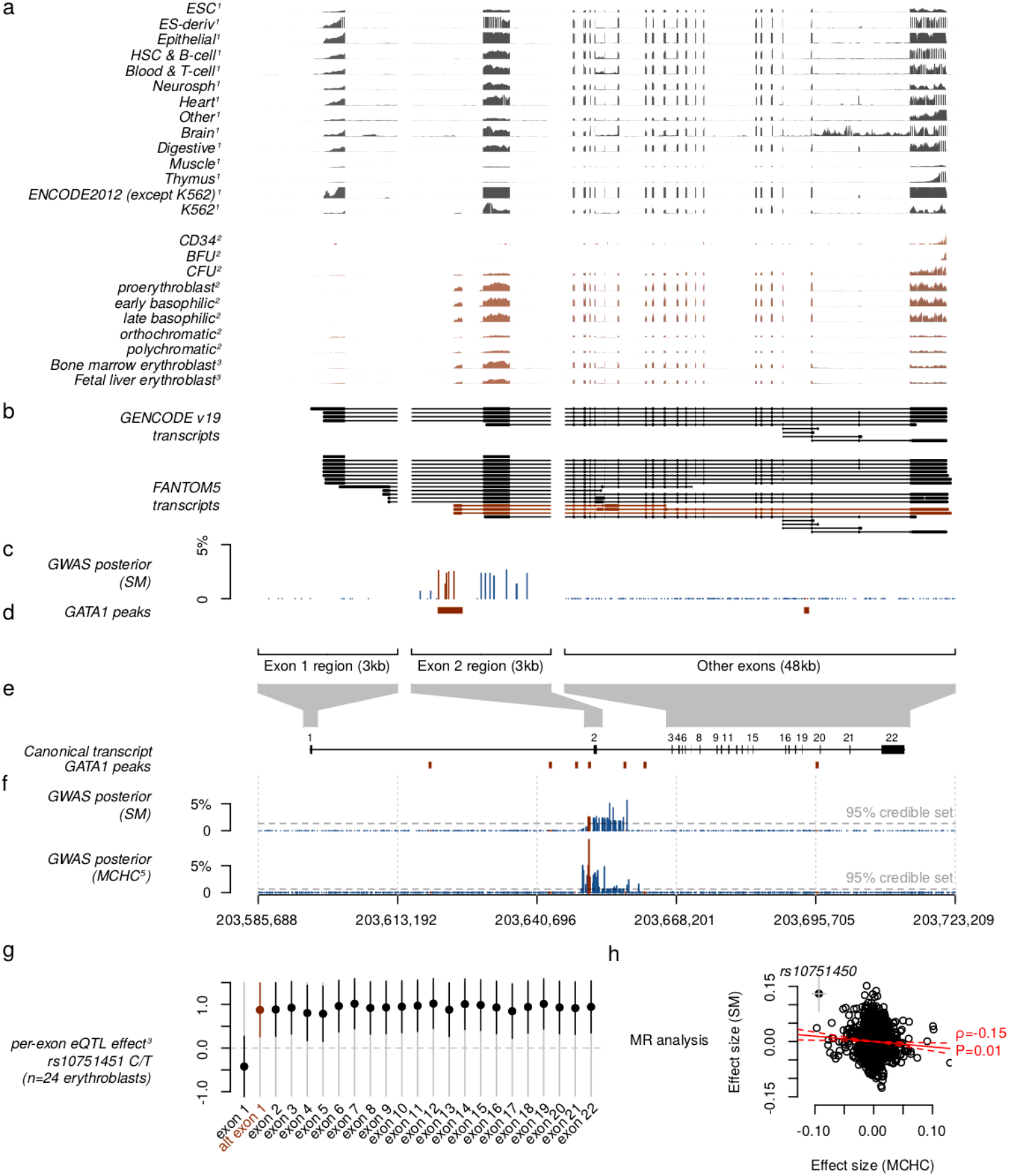
the *ATP2B4* association is driven by an erythrocyte-specific transcription start site. **a)** normalized RNA-seq coverage for (1) 56 cell types from Roadmap Epigenomics^28^ and ENCODE, (2) human HSPCs and experimentally differentiated erythroid cells from three biological replicates^30^, (3) ex-vivo differentiated adult and fetal human erythroblasts from 24 individuals^31^, and (4) circulating erythrocytes from ^44^. Coverage is shown across expanded regions of *ATP2B4* exon 1, exon 2 including the putative alternative first exon, and the remaining exons. For Roadmap and ENCODE cell types, the plot reflects the maximum normalized coverage across cell types in each tissue. Tissue groups are as taken from the Roadmap metadata. For other cells, coverage is summed over samples within the group and normalized by the mean coverage across ATP2B4 exons. **b)** *ATP2B4* transcripts from the GENCODE^40^ and FANTOM5^43^ transcript models. Transcripts including the alternative first exon are highlighted in red. **c)** detail of association evidence, showing posterior evidence for association with SM under the assumption of a single causal variant. **d)** position of GATA1 binding peaks^27^. **e)** location and size of the expanded regions shown against the full-length transcript, with GATA1 binding peaks shown. **f)** posterior evidence for association with SM as in c), and with mean corpuscular haemoglobin concentrations (MCHC)^35^, under the assumption of a single causal variant in each trait separately. **g)** estimated eQTL effect of rs10751451 against each exon of *ATP2B4*, using a linear regression model of residuals of FPKM against genotype after correcting for cell development stage^31^. For comparable visualisation across exons, we normalized per-exon residual FPKM values by the mean across samples in each exon. **h)** Mendelian randomization analysis of SM and MCHC at 2130 ‘sentinel’ SNPs previously identified as associated with haematopoetic traits^35^ and having association results in our study. Points reflect the posterior effect size estimates on SM (y axis) and MCHC (x axis), conditional on the fitted model. A bivariate Gaussian model of effect sizes is assumed and variants are assumed to act independently. Red solid and dotted lines show the maximum likelihood estimate of the effect of mchc on SM (*ρ*) and its 95% confidence interval. The P-value reflects a likelihood ratio test P-value against the null that ρ=0.

The situation outlined above for *ATP2B4* is reminiscent of the well-known mutation at the Duffy blood group locus (*DARC/ACKR1*), which protects against *P. vivax* malaria by preventing erythrocytic expression of the Duffy antigen receptor through disruption of a GATA1 binding site ^45,46^. However, the mechanism by which *ATP2B4* affects parasite processes remains unknown. The malaria-protective allele at rs10751451 is known to associate with red blood cell indices (notably with increased mean corpuscular haemoglobin concentration (MCHC) ^31,35,47^). We found some evidence that genetic predisposition to high MCHC levels may itself decrease malaria susceptibility (negative observed correlation between MCHC and SM effect size at sentinel SNPs previously identified as associated with haematological indices in European individuals ^35^, *ρ*=−0.15, P=0.01; in a mendelian randomization framework; **Figure 5h**). However, the protective effect at *ATP2B4* appears substantially stronger than this trend and suggests a genuinely pleiotropic effect whose mechanism against malaria is yet to be determined.

Our analysis of functional annotations revealed a number of additional variants with some evidence of function (**Table S4**). These include a potentially regulatory variant near *H6PD*, as well as a variant upstream of *VAC14* which has been implicated in Typhoid fever ^48^ and *S.Typhi* invasion ^34^, detailed further in **Text S4**.

### Imputation reveals a lack of association with classical HLA alleles

Motivated by the modest evidence for association observed in the HLA (**Table S2**), we used a published method ^49^ to impute classical HLA alleles and tested for association with each allele as described above for genome-wide variants (**Figure 6a**). The strongest evidence for association with an HLA antigen was observed at HLA-B*42 (*BF_avg_* = 937; synonymous with HLA-B*42:01 in this panel). By contrast, an order of magnitude stronger evidence was observed at regional SNPs, including rs2523650 (*BF_avg_*=2.2×10^4^; additive model OR=0.88 for the nonreference ‘C’ allele; 95%CI=0.94-0.93; *P_add_*=2.4×10^−6^) which has previously been associated with blood cell traits and expression of regional genes (**Table S4**), but this signal did not replicate in additional samples (*BF_replication_*=0.09). Most notably, the HLA-B*53 allele, which has previously been reported as associated with strong protection against severe malaria in the Gambia ^1^, showed no evidence for association across populations (*BF_avg_*=0.28; OR = 0.99, 95%CI = 0.91-1.07 in fixed-effect meta-analysis under a dominance model where the effect is due to presence of the antigen) or in any individual population (e.g. OR= 0.99, 95%CI = 0.88-1.13 in the Gambia, **Figure S6**).

**Figure 6:**
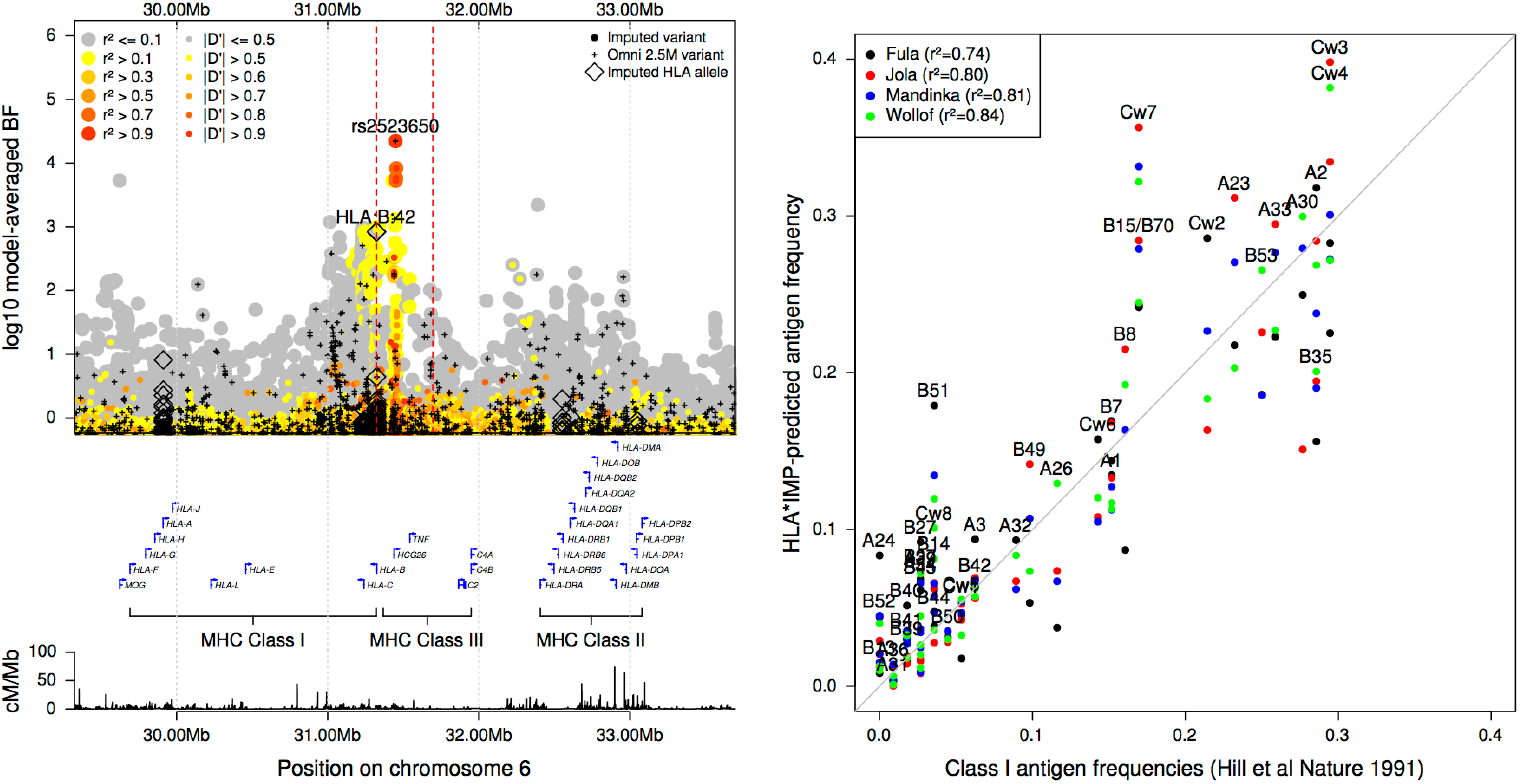
Evidence for association across the HLA. **a)** Evidence for association at genotyped SNPs (black plusses), imputed SNPs and INDELs (circles), and imputed classical HLA alleles (black diamonds) across the HLA region. Points are coloured by LD with rs2523650 as estimated in the African reference panel populations. Selected regional genes and the Hapmap combined recombination map are shown below. **b)** Comparison of HLA Class I antigen frequencies in distinct sample sets from The Gambia. X-axis: antigen frequencies obtained by serotyping of 112 healthy adults ^1^; y-axis: inferred antigen frequencies based on imputation of 2-digit alleles in the four major ethnic groups (colours) in our Gambian dataset. B*15 alleles encode a number of antigens^2^ including B70, and we combine results for B15 and B70 here.

We considered whether imperfect imputation could explain the lack of observed association with HLA alleles. Imputed HLA antigen frequencies were broadly consistent with published frequencies estimated by serotyping in the same populations (**Figure 6b**), and HLA-B*53 was imputed at reasonably high frequency and high confidence (estimated frequency = 7-18%, IMPUTE info score > 0.94 in all African populations). To directly test imputation accuracy, we obtained HLA types of a subset of 31 Gambian case individuals by Sanger sequencing (**Table S5 and Text S5**), and confirmed that HLA B*53 was relatively well imputed in these samples (e.g. correlation = 0.92 between imputed dosage and true number of copies of the allele, reflecting 5 individuals imputed with probability > 0.75 as heterozygote carriers of B*53, of which one appears incorrect). However, the closely related B*35 allele was much less accurately imputed, with four of eight carriers imputed as having non-B*35 genotypes, including one imputed to carry B*53. This may be the reason for the relatively low observed frequency of imputed B*35 alleles (**Figure 6b**), and is notable because B*53 is thought to have arisen from B*35 via a gene conversion^50^. Thus, while our failure to replicate this widely cited association appears robust, future work to improve HLA inference in these populations is needed to provide additional clarity.

### Genome-wide data suggest evidence for polygenic selection

Malaria is regarded as having played a strong role in shaping the human genome through natural selection^51,52^, and observations of allele frequency differences are commonly used to motivate the study of specific mutations (e.g.^53–55^). We sought to assess whether variants associated with malaria susceptibility show evidence of selection by comparing with genome-wide allele frequency variation between populations (**Figure 7** and **Figure S7-S9**). We found that the alleles with the strongest evidence of protection tend to be at lower frequency in non-African populations than expected given the genome-wide distribution (e.g. conditional rank of allele in European populations, given African allele count (rank_EUR_) < 0.5 for 10 of 12 regions with *BF_avg_* >10,000; 46 of 90 regions with *BF_avg_* >1,000; based on reference panel allele counts; **Figure 7a**), consistent with the hypothesis that these alleles have been maintained at high frequency in African populations by positive selection. However, this comparison provides extremely modest levels of evidence - even for individual alleles where selection of this type is well accepted. We similarly found modest evidence for within-Africa differentiation (e.g. *P*=1×10^−3^ against the null model of no differentiation, for the 12 regions with *BF_avg_*>10,000; P=0.02 across 92 regions with *BF_avg_* > 1,000, excluding all but one variant within the HLA region; **Figure 7b-c**). A further discussion of differentiated loci is found in **Text S9**.

**Figure 7:**
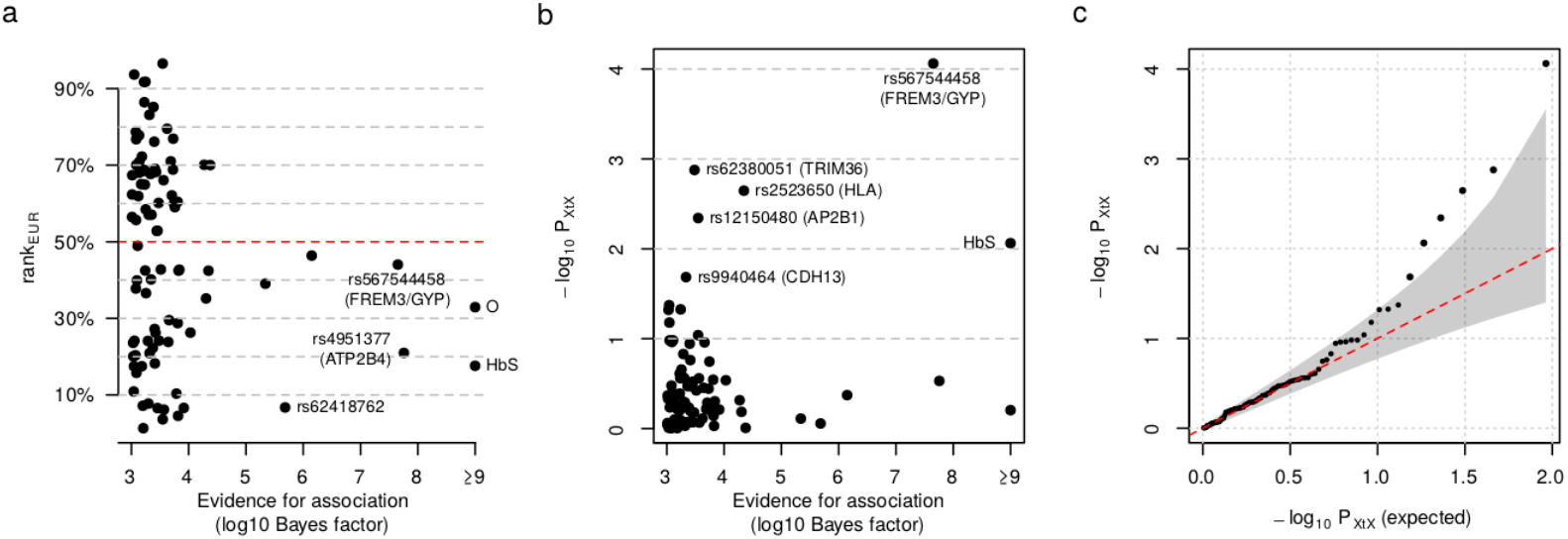
Empirical evidence for frequency differentiation of the most-associated alleles. **a)** The European population rank (rank_EUR_, y axis) plotted against the evidence for association (log10 *BF_avg_*, x axis) for the protective allele at each of the 90 lead variants satisfying *BF_avg_* > 1000 and having assigned ancestral allele. For each allele A, rank_EUR_ is defined as the proportion of alleles genome-wide having lower or equal count than *A* in European populations, conditional on having the same frequency in African populations. In practice we estimate rank_EUR_ empirically from the joint allele count distribution in reference panel samples (**Figure S7**), further stratified according to whether the protective allele is ancestral or derived. On average rank_EUR_ is expected to equal 50% (red dashed line). Points are labelled by the rsid, nearest or relevant gene(s), or by functional variant where known; “O” refers to rs8176719 which determines O blood group. **b)** Evidence for within-Africa differentiation (*P_XtX_* y axis), plotted against the evidence for association (log10 *BF_avg_*, x axis) for each of the 92 lead variants satisfying *BF_avg_* > 1000, after removing all but rs2523650 from within the HLA region. *P_XtX_* is computed from an empirical null distribution of allele frequencies learnt across control samples in the seven largest African populations (**Figures S8-S9**). c) quantile-quantile plot for *P_XtX_* across the top 92 regions in panel b.

These analyses add new weight to the hypothesis that malaria-driven selection has played a polygenic role in shaping human genomes in endemic populations. However, this evidence is rather weak, and it is equally clear that many of the most differentiated alleles, including those within the HLA (e.g. HLA-DPB*01; frequency range 25-54% in African populations; P_XtX_ = 1.3×10^−14^), variants in *CD36* (e.g. rs73711929, frequency range 2-20% in African populations; P_XtX_ = 8×10^−20^), as well as variants in regions associated with skin pigmentation^56^ (e.g. rs1426654; frequency range 83-99% in African populations; P_XtX_=3×10^−15^) are not associated with susceptibility to *P. falciparum* malaria (**Figure S11**). Other selective forces must have driven the evolution of these alleles^57^ and in many cases these remain to be identified.

## Discussion

This analysis identifies five replicable loci with strong evidence for association with resistance to severe malaria: *HBB, ABO, ATP2B4*, the glycophorin region on chromosome 4, and a new locus between *MAP3K7* and *EPHA7* on chromosome 6 whose causal mechanism is unknown. The *HBB, ABO* and glycophorin associations correspond to known phenotypic variants of human red blood cells, namely sickle cell trait and the O and Dantu blood group phenotypes. Building on previous work^31^, our analysis indicates that the associated haplotype in *ATP2B4* also has a specific effect on erythrocyte function. The putative protective allele disrupts a GATA1 binding site upstream of a transcription start site that is only active in erythroid cells, and thus reduces expression of this gene in erythrocyte precursors. This is reminiscent of the protective effect of the Duffy null allele against *P. vivax* malaria, which also involves disruption of a GATA1 binding site causing erythrocyte-specific suppression of the Duffy antigen receptor ^45,46^, and it adds to growing evidence that tissue-specific transcriptional initiation is an important factor in determining human phenotypes ^58,59^. *ATP2B4* encodes the main erythrocyte calcium exporter, and the associated haplotype has been experimentally shown to control calcium efflux and hydration^31,39^. This is consistent with the observed effects on red cell traits^35^ and with the hypothesis that *ATP2B4* modulates parasite growth within erythrocytes, perhaps by affecting calcium signaling mechanisms that are essential to several stages of the parasite lifecycle^60,61^. In theory, this finding raises the possibility of therapeutic blockage of *ATP2B4* in erythrocytes without otherwise affecting physiology. Based on the estimated effect of the associated SNPs, this might provide ~1.5-fold protection against severe malaria and be efficacious in the ~90% of individuals who do not currently carry the protective homozygous genotype at this locus, and it is open to speculation whether this could lead to worthwhile therapy in practice.

The five loci identified so far appear to explain around 11% of the total genetic contribution to variation in malaria susceptibility (**Figure S4**). It is possible that this estimate is confounded, e.g. by unusual LD patterns or by environmental factors such as variation in underlying infection rates, and it should likely be treated with caution. Nevertheless, it raises the question as to why only five loci can be reliably detected at present. Selection due to malaria has been sufficiently strong to maintain alleles such as sickle haemoglobin at high frequency in affected African populations^62^ (**Figure 7**). However, it has evidently also led to complex patterns of variation at malaria resistance loci. This is exemplified by the emergence of haemoglobin C and multiple haplotypes carrying sickle haemoglobin at *HBB*, and by copy number variation observed at the alpha globin and glycophorin loci^14,63^. Additionally there is evidence for long-term balancing selection at the ABO^64^ and glycophorin^65^ loci which may be malaria-related. In theory, the simplest outcome for a strongly protective allele is that it would sweep to fixation, but unlike for *P.vivax*^66^ no sweep of a mutation providing resistance to *P. falciparum* malaria is known. Remaining protective alleles may therefore have effect sizes too small to be subject to strong selection, requiring large sample sizes to detect. Genome-wide association studies of other common diseases have shown the benefit of exceptionally large sample sizes ^67–71^, and although there are many practical obstacles to achieving this for severe malaria, particularly since its incidence has fallen in recent years due to improved control measures^72,73^, efforts to collect new samples and to combine data across studies^61^ are warranted. It is also possible that relevant alleles may have become balanced due to pleiotropic effects, or be subject to compensatory parasite adaptation - both scenarios that might lead to maintenance of allelic diversity. Dissecting such signals is naturally challenging for GWAS approaches and may require other techniques, such as sequencing and joint analysis of host and parasite genomes from infected individuals, to address.

## Author contributions

**Manuscript preparation**: Gavin Band and Dominic P. Kwiatkowski in collaboration with all authors.

**Analysis working group**: Gavin Band, Quang Si Le, Geraldine M. Clarke, Ellen M. Leffler, George B.J. Busby, Jennifer Shelton, Chris C.A. Spencer, Kirk A. Rockett, Dominic P. Kwiatkowski

**Recruitment of subjects, sample and clinical data collection**: Muminatou Jallow, David J. Conway, Kalifa A. Bojang, Fatoumatta Sisay-Joof, Giorgio Sirugo, Umberto d’Alessandro, Ousmane B. Toure, Mahamadou A. Thera, Salimata Konate, Sibiri Sissoko, Ogobara Doumbo, Valentina D. Mangano, Edith C. Bougouma, Sodiomon B. Sirima, David Modiano, Lucas N. Amenga-Etego, Anita K. Ghansah, Kwadwo A. Koram, Abraham V. O. Hodgson, Michael D. Wilson, Anthony Enimil, Daniel Ansong, Jennifer Evans, Tsiri Agbenyega, Olukemi K. Amodu, Subulade A. Ademola, Tobias O. Apinjoh, Eric Achidi, Carolyne M. Ndila, Kevin Marsh, Thomas N. Williams, Alphaxard Manjurano, Eleanor M. Riley, Chris Drakeley, Hugh Reyburn, Malcolm Molyneux, Terrie Taylor, Sarah J. Dunstan, Nguyen Hoan Phu, Nguyen Thi Ngoc Quyen, Cao Quang Thai, Tran Tinh Hien, Jeremy Farrar, Yik Ying Teo, Laurens Manning, Moses Laman, Pascal Michon, Harin Karunajeewa, Peter Siba, Steve Allen, Angela Allen, Melanie Bahlo, Timothy M. E. Davis, Ivo Mueller

**HLA sequencing and imputation**: Alexander Dilthey, Alexander J. Mentzer, Gil McVean

**Genotype and sequence data production**: Kirk A. Rockett, Christina Hubbart, Anna E. Jeffreys, Kate Rowlands, Eleanor Drury, Jim Stalker, Katja Kivinen

**Study site principal investigators**: Kalifa A. Bojang, Ogobara Doumbo, David Modiano, Kwadwo A. Koram, Tsiri Agbenyega, Olukemi K. Amodu, Eric Achidi, Kevin Marsh, Thomas N. Williams, Eleanor M. Riley, Malcolm Molyneux, Terrie Taylor, Sarah J. Dunstan, Jeremy Farrar

**Network coordination and ethics**: Victoria Cornelius, Angeliki Kerasidou

**Project oversight / management**: Chris C.A. Spencer, Kirk A. Rockett, Dominic P. Kwiatkowski

## Acknowledgements

This manuscript is dedicated to the memory of Professor Ogobara Doumbo (1956-2018).

We thank all the study participants and the members of the MalariaGEN Consortial Projects 1 and 3 (https://www.malariagen.net/projects). These MalariaGEN Consortial Projects were supported by Wellcome (WT077383/Z/05/Z) and the Bill & Melinda Gates Foundation through the Foundations of the National Institutes of Health (566) as part of the Grand Challenges in Global Health Initiative. The Resource Centre for Genomic Epidemiology of Malaria is supported by Wellcome (090770/Z/09/Z; 204911/Z/16/Z). This research was supported by the Medical Research Council (G0600718; G0600230; MR/M006212/1). Wellcome also provides core awards to The Wellcome Centre for Human Genetics (203141/Z/16/Z) and the Wellcome Sanger Institute (206194). C.C.A.S was supported by a Wellcome Trust Career Development Fellowship grant (097364/Z/11/Z). O.K.A was supported by the European Community under grant agreement LSHP-CT-2004-503578 and Seventh FrameworkProgramme (FP7/2007-2013) under grant agreement no. 242095. This paper is submitted with the permission of the Director of KEMRI. A.J.M. was supported by an Oxford University Clinical Academic School Transitional Fellowship and a Wellcome Trust Clinical Research Training Fellowship reference 106289/Z/14/Z. We thank Will Rayner at the Wellcome Centre for Human genetics, Oxford, for supplying genotyping platform strand files. Eric Achidi received partial funding from the European Community’s Seventh Framework Programme (FP7/2007-2013) under grant agreement No. 242095 – EVIMalaR and the Central African Network for Tuberculosis, HIV/AIDS and Malaria (CANTAM) funded by the European and Developing Countries Clinical Trials Partnership (EDCTP). Thomas N. Williams is funded by Senior Fellowships from the Wellcome Trust (076934/Z/05/Z and 091758/Z/10/Z) and through the European Community’s Seventh Framework Programme (FP7/2007-2013) under grant agreement No. 242095 – EVIMalaR. The KEMRI-Wellcome Trust Programme is funded through core support from the Wellcome Trust. Carolyne Ndila is supported through a strategic award to the KEMRI-Wellcome Trust Programme by the Wellcome Trust (084538). Tanzania/KCMC/JMP received funding from MRC grant number (G9901439). The Malawi-Liverpool-Wellcome Trust Clinical Research Programme (MLW) is a Major Overseas Programme of Wellcome. Malcolm Molyneux was funded by a Wellcome Trust Research Leave Fellowship. V.D.M. was funded by Istituto Pasteur-Fondazione Cenci Bolognetti, BioMalPar and Evimalar (European Community FP6,FP7). HLA typing was supported by G.M. under Wellcome Trust grant 100956/Z/13/Z We thank Sarah Peacock and Sarah Maxwell at the Tissue Typing Laboratory, Addenbrooke’s hospital, Cambridge, for their assistance with HLA typing.

## Data availability

The following data will be made available prior to publication of this manuscript. Illumina Omni 2.5M genotype data from study samples (**Figure 1a**, right panel), are being made available through the European Genome-Phenome Archive (EGA) under study accession EGAS0000100131. Phased and imputed genotypes from these samples will also be deposited in the EGA. Whole-genome sequence read data for Gambian Genome Variation Project samples (**Figure 1a**, left panel) is available through the European Nucleotide Archive under PRJEB3013 (Fula), PRJEB3252 (Jola), PRJEB1682 (Mandinka), and PRJEB1323 (Wollof). Read data for sequenced samples from Burkina Faso, Cameroon, and Tanzania (**Figure 1a**, left panel) are available through the EGA under study accession EGAS00001003648. Genotypes generated on the Sequenom iPLEX Mass-Array platform for selected variants in discovery and replication samples will be deposited in the EGA. HLA allele genotypes for 31 Gambian individuals (Table S5) will be deposited in the EGA. A full set of association summary statistics underlying our analysis will also be made available to download. Full information on data generated by MalariaGEN is available on our website (www.malariagen.net).

## Methods

### 1. Collection and processing of whole-genome sequence data

#### 1.1. Collection and sequencing of African individuals

Blood samples were collected from a total of 773 individuals from Gambia, Burkina Faso, Cameroon and Tanzania, and sequenced to an average of 10x coverage on the Illumina HiSeq 2000 platform at the Wellcome Trust Sanger Institute, using 100bp paired end reads. In addition, one Gambian trio was additionally sequenced using three alternate library preparation methods (low-quantity and whole-genome amplification pipelines) for a total of 782 sequenced samples.

#### 1.2. Data alignment and QC

Reads were mapped to the GRCh37 human reference genome with additional sequences as modified by the 1000 Genomes Project (hs37d5.fa), using BWA^74^ with base quality score recalibration (BQSR) and local realignment around known indels as implemented in GATK^75^.

#### 1.3. Variant detection

We used GATK UnifiedGenotyper ^75^ to identify potential polymorphic sites on autosomal chromosomes, using all 782 samples as well as the 680 samples of African ancestry from the 1000 Genomes Project, and running in 50kb chunks.

We used GATK VQSR^75^ to separately filter SNPs and indels. For SNPs, we included three training sets: HapMap SNPs, SNPs on the Omni 2.5M array, and 1000 Genomes Phase 3 sites. For indels we used the ‘Mills’ training set as recommended in the GATK documentation. We based filtering on read depth, mapping quality, quality by depth, the MQRankSum and ReadPosRankSum measures of bias in reference versus alternate allele mapping quality or read positions, and tests for strand bias and inbreeding coefficient. We filtered variants applying a sensitivity setting of 99.5% (--ts_filter_level 0.95) for SNPs and 95% for INDELs (--ts_filter_level 0.95). We further restricted to biallelic sites.

#### 1.4. Genotype likelihood computation, genotyping and phasing

We used GATK HaplotypeCaller ^76^to compute genotype likelihoods at all sites passing the filter described above as well as sites in 1000 Genomes Project Phase 3 reference panel. We then used BEAGLE v4.0 ^77^ to generate genotype calls from the genotype likelihoods. We range BEAGLE in windows of 50,000 SNPs, including trio information for Gambian samples. Due to the observed long runtimes when sampling trios, we specified trioscale=3 when running BEAGLE to reduce computation times. We then concatenated results across chromosomes.

To estimate haplotypes, we first removed variants not present in the 1000 Genomes Phase 3 panel or variants that had alleles differing from those in the 1000 Genomes Phase 3 panel, using the ‘check’ mode of SHAPEIT2^78^. We then used SHAPEIT2 to phase each chromosome. We included trio information for Gambian samples, specified a window size of 0.5 (--window), an effective sample size of 17,469 (--effective-size), 200 model states (--states), included 1000 Genomes Phase 3 haplotypes in the phasing process (--input-ref), and used the hapmap combined recombination map in build 37 coordinates (-M).

#### 1.5. Construction of combined imputation reference panel

To construct a combined imputation reference panel, we removed repeated samples. We combined phased haplotypes at the remaining samples with those from the 1000 Genomes Project phase 3 panel. The final panel contains 77,931,101 variants and 3,277 samples, of which 773 are from our set of sequence data and 2,504 are from the 1000 Genomes panel. In total, the panel includes 1,434 individuals with recent African ancestry (including 157 African-american samples), and non-African samples as previously described^12^.

### 2. Collection and processing of GWAS data

#### 2.1. GWAS data collection

Study samples were collected and prepared for genotyping as described previously ^7,10^

#### 2.2. Sample genotyping

Sample genotyping and genotype calling were performed as described previously^10^. In brief, all samples were genotyped on the Illumina Omni 2.5M platform at the Wellcome Trust Sanger Institute. We used three genotype calling algorithms (Illumina Gencall, GenoSNP, and Illuminus) and formed the consensus genotype call by taking the consensus of the three algorithms, or of two algorithms where the third algorithm reported a missing call. Genotypes for which a discrepant call were made were treated as missing.

#### 2.3. Data alignment

We used strand information from array manifest files provided by Illumina, and those from a remapping process implemented by William Rayner at the Wellcome Centre for Human Genetics, Oxford, to determine the strand of each assay on each of the two genotyping platforms used. We omitted variants where mapping or strand information was discrepant, with some adjustments for Omni 2.5M ‘quad’ array as described below. Additionally, we annotated each variant with the reference allele as taken from the build 37 reference sequence FASTA file.

We used these data with QCTOOL to realign study genotypes. The resulting ‘aligned’ datasets are encoded so that the first allele reflects the reference allele, the second allele the non-reference allele, and all alleles are expressed relative to the forward strand of the reference sequence, simplifying downstream analysis.

As a sanity check we plotted the estimated frequency of each non-reference allele against the frequency in the closest reference panel group in each populations. The Kenyan dataset (which was typed using the Omni 2.5M quad ‘D’ version manifest) showed a substantial number of SNPs with frequencies obviously wrongly specified. Comparison of the flanking sequence and alleles reported in the manifest file identified 18,763 SNPs that had alleles coded the incorrectly. We manually fixed these SNPs by recoding these alleles, making them consistent with the flanking sequence, the more recent ‘H’ version of the manifest, and the Omni 2.5M ‘octo’ chip manifest. We also updated the positions of 495 SNPs annotated as lying on the pseudo-autosomal part of the X chromosome (Chr=‘XY’ in the Illumina manifest) but with position = 0. We applied these updates, repeated the alignment process for Kenya, and regenerated the frequency plot. A small number of SNPs in each population still show distinct frequencies compared to reference panel data (**Figure S13**).

#### 2.4. Sample quality control

We used QCTOOL to compute the proportion of missing genotype calls, the average proportion of heterozygous calls per sample, and the average X channel and Y channel intensities across autosomal chromosomes in each population. Values were averaged over a subset of SNPs (’the QC SNP set’) chosen to have low missing rates in all populations. We ran ABERRANT^79^ to identify individuals with outlying mean intensities separately in each population. Considerable variation in the spread of quality between populations was observed, and we picked population-specific values of the ABERRANT lambda parameter based on visual inspection of the plots. We also manually adjusted the ABERRANT-derived exclusions to include a small number of samples in each population which were observed to visually cluster with the main set of included samples, but called as outlying by ABERRANT.

We next plotted logit(missing call rate) against average heterozygosity in each population. We excluded samples with greater than 10% missingness or heterozygosity outside the range 0.2-0.4 in African populations, below 0.175 in Vietnam, or below 0.13 in PNG outright, across the QC SNP set. We then applied ABERRANT to the remaining non-excluded samples choosing a value of the lambda parameter per population based on inspection. As for intensity outliers, we specifically included some groups of samples visually clustering with the main group of samples but called as outlier by ABERRANT. We plotted heterozygosity against missingness annotating the removed samples. Figures **S14** and **S15** show mean intensity, missingness, and heterozygosity, computed across all variants, with excluded samples annotated.

The above process results in a set of 18,515 samples that appear to be relatively well genotyped across the autosomes. In addition, for some purposes described below we used a smaller set 17,664 samples (referred to as the ‘conservative’ set of samples) obtained by repeating the above process with more conservative values of the ABERRANT lambda parameter.

#### 2.5. Gender determination

Array intensity values on the sex chromosomes are informative about the underlying sex of samples and we used these to confirm gender. Specifically, we plotted overall intensity (X channel plus Y channel intensity) averaged across the X chromosome against overall intensity averaged across the Y chromosome (**Figure S16**). To infer sex, we modeled these average intensities as drawn from a bivariate normal distribution, with separate mean and covariance parameters in males and females. Almost all samples had sex determined by direct genotyping of amelogenin SNPs ^7^ and we used these assignments to estimate the model parameters. A small number of samples were observed to lie near a point consistent with female-like intensity values on the X chromosome but male-like intensity values on the Y chromosome; this could potentially indicate males with an extra copy of the X chromosome. We therefore considered an additional ‘mixed’ cluster with parameters computed from the fitted male and female clusters on the X and Y chromosome, and with no covariance between X and Y chromosome intensities.

We used the ‘conservative’ set of included samples to estimate these parameters. Based on the fitted model we then probabilistically assigned gender to all samples sample based on Bayes rule:

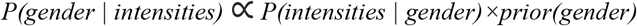

Here the first term is computed using the bivariate normal model described above, and we normalize the formula under the assumption that every sample is either male, female, or ‘mixed’. We specified a 1% prior probability of ‘mixed’ gender, and equal prior probability of 49.5% on each of the male and female clusters. Finally, we assigned each gender having at least 70% posterior probability and treated the gender of other samples as uncalled. Of the 18,515 samples in the QC set, 20 had mixed or missing gender by this assignment, while 38 had assignment discrepant with the amelogenin-determined gender. We removed mixed/missing gender samples from downstream analyses.

We noted a trend for male samples to have slightly higher mean intensity values across autosomes than females on average. We speculate this may be because, lacking one copy of the X chromosome, males carry a higher proportion of autosomal DNA overall.

#### 2.6. Sample relatedness and principal component analysis

We used QCTOOL to compute a matrix of pairwise relatedness coefficients in each population. To do this, we first chose a subset of SNPs with using a preliminary version of the SNP QC process described below, and used inthinnerator (**Online Resources**) to thin this set of SNPs to lie no closer than 0.02cM apart in the Hapmap combined recombination map. We additionally excluded SNPs from regions of known associations, the region chr6:25,000,000-40,000,000 which contains the HLA, and the region of the common inversion on chromosome 8. We used QCTOOL with the resulting set of 157,085 SNPs to compute the normalized allele sharing (‘relatedness’) matrix

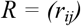

where *r_ij_* reflects the correlation in genotype between sample i and j, normalized by variant frequency. The number *r_ij_* may be treated as an estimate of relatedness with values near 1 reflecting identity and values near 0 reflecting unrelated samples, relative to the average relatedness in the population ^80,81^.

For each pair of samples with relatedness > 0.75 we noted the sample with the highest rate of missing genotype calls. A total of 538 samples were identified in this way, the majority from the Malawi and Kenya datasets, and likely reflect duplicate genotyping. We removed these samples from the datasets.

For each remaining pair of samples with relatedness > 0.2 we again noted the sample with the highest rate of missing genotype calls; these represent samples that are relatively closely related to other samples in the same study sample. A total of 801 samples were identified in this way.

We formed the eigendecomposition of the matrix *R* and used this to compute and plot principal components (PCs) in each population. Based on visual inspection of these plots, we further identified a total of 40 samples outlying on any of the first ten PCs. We recomputed *R* and the PCs after excluding these individuals, to produce a set of PCs for association analysis.

#### 2.7. Summary of sample QC

In total 17,960 samples passed QC criteria, had non-missing / non-mixed gender assignment, and were not identified as duplicates as described above. We took all of these samples through to the phasing and imputation as described below. Of these samples, a total of 17,056 samples were identified as not outlying on principal components, not closely related and having assigned case/control status, and were included in downstream analyses. **Table S7** summarises the sample QC.

#### 2.8. Re-estimating genotype cluster positions

We used intensities and genotype calls for the ‘conservative’ sample set to re-estimate the positions of genotype clusters at each SNP in each population using QCTOOL. QCTOOL fits a multivariate t distribution to each genotype cluster; we specified 5 degrees of freedom. We used these clusters to annotate cluster plots of individual variants and to implement the recall test described below.

#### 2.9. Computation of SNP QC metrics

We based QC on several metrics, computed separately in each population:

- The estimated minor allele frequency (MAF).
- The proportion of missing genotype calls (missingness).
- A P-value for Hardy-Weinberg equilibrium ^82^, computed in control samples (P_HWE_).
- A ‘plate test’ P-value against the null model that genotypes are uncorrelated with the plate on which each sample was genotyped (P_plate_), described further below.
- A ‘recall test’ P-value, which compares the genotype frequencies to frequencies after a recalling process based on the estimated cluster positions across populations. This is described further below.

For the plate test, we implemented a logistic regression model treating the genotype as outcome and an indicator of the 96-well plate on which the sample was supplied for genotyping as a predictor in each population. We included case/control status and an initial set of five principal components as covariates and computed a likelihood ratio test P-value for the effect of including plate as a predictor.

The ‘recall’ test was motivated by the observation that some otherwise well-typed SNPs were poorly genotyped in individual populations. We implemented a re-calling procedure in which the intensity data from each population was matched against a mixture model of intensities based on the cluster positions learnt in all other populations. In each population we then computed a p-value against the null model that the frequency in the original and re-called genotypes were the same using Fisher’s exact test.

We observed study populations to vary considerably both in size and in quality of typing, with the two smallest African sets (Mali and Nigeria) and, to a lesser extent, Malawi, having higher missing data rates, and Kenya having lower rates (**Figure S17**). Inspection of cluster plots suggests that the higher missing rates are largely due to higher levels of noise in intensity values for these populations.

#### 2.10. Choice of SNP QC criteria

We used association test statistics computed under a general logistic regression model, including additive and heterozygote parameters, as a guide to choosing appropriate thresholds for the above metrics. Specifically, we tested for association with severe malaria, including 5 principal components as covariates in each population. For given QC criteria we plotted association test results annotating QC fails, along with qq plots for the variants passing QC.

In choosing SNP QC thresholds, we were motivated by the observation that the combination of consensus genotype calling (which is relatively conservative in calling genotypes) with statistical phasing across populations (which fills in missing genotypes based on inferred haplotype sharing) would likely produce a high-quality set of haplotypes for downstream inference. For this reason we aimed to produce a single set of SNPs with high quality data across populations for input to the phasing process. Based on this we chose a set of criteria applied to the nine largest populations to compute a list of SNPs to include. We removed SNPs with missingness > 2.5% in Kenya and PNG, > 5% in Gambia, Burkina Faso, Ghana, Cameroon and Vietnam, and > 10% in Malawi and Tanzania. We also excluded SNPs with P<1×10^−20^ for HWE, P < 1×10^−3^ for the plate test and P<1×10^−6^ for the recall test in each population. Finally, we included each of the remaining SNPs that was at MAF > 1% in at least two populations and passed the above thresholds in all nine populations considered. Data from the Mali and Nigeria populations, which had the lowest sample counts (484 and 133 samples, respectively) and empirically lower rates of high-quality genotyping (**Figure S17**), was not used to inform the SNP selection process.

The criteria above were observed to lead to essentially uninflated association association test statistics (**Figure S18**). In particular we noted that a set of SNPs close to the ends of chromosomes were excluded by these criteria; more information on this is provided in **Text S2** and **Figure S19**.

#### 2.11. X chromosome SNP QC

To QC the X chromosome we proceeded as for autosomes with the following adjustments. We excluded the pseudoautosomal region (defined as variants with position < 2,699,520 or > 154,931,044; this region contained ~400 typed SNPs). We observed elevated rates of males called as heterozygous within the X transposed region (XTR, defined as chrX: 88,457,462-92,374,313) and excluded this region as well. Approximately 3,400 SNPs were removed this way. In computing summary statistics for males we treated all heterozygous calls as missing and homozygous calls as the corresponding hemizygous calls. We applied the SNP missingness threshold and the plate test separately in males and females; we replaced the Hardy-Weinberg equilibrium test with a test of difference in frequency between males and females (based on a likelihood ratio test P < 1×10^−6^). We did not implement the recall test for X chromosome SNPs.

#### 2.12. Assessment of cluster plots and additional QC

As an additional sanity check, we plotted cluster plots for all SNPs with genotypic association test P < 1×10^−5^ and manually excluded those with obviously problematic cluster plots. Some SNPs are targeted by duplicate assays on the Illumina Omni 2.5M array. For each duplicate assay, we excluded the assay showing highest missingness, on average, across populations. We also restricted to SNPs present on both ‘quad’ and ‘octo’ platforms.

#### 2.13. Summary of SNP QC

Table S8 shows the number of SNPs excluded by each criteria and the SNPs included in the final QC set. We used QCTOOL to subset the data for each population to the set of included SNPs. We then merged data across populations, generating a single GEN format file per chromosome containing all the 17,960 high-quality samples included in the study.

### 3. Phasing and imputation

#### 3.1. Genome-wide phasing

We used SHAPEIT2 ^78^ to jointly phase the 17,960 samples passing QC. In detail, we ran SHAPEIT2 on each chromosome separately specifying an effective sample size of 17,469 and 200 copying states, and using the HapMap combined recombination map. As above we note that in addition to phasing, SHAPEIT2 imputes missing genotypes at typed SNPs with missing data, so that the output of this step contains hard-called phased genotypes with no missing data.

#### 3.2. Recomputation of principal components

Following phasing we recomputed principal components across all populations, across African populations, and in each population separately, using phased genotypes. These PCs are shown in **Figure 1** and **Figure S2**.

#### 3.3. Genome-wide imputation

We used IMPUTE2 (v2.3.2) ^83^ to impute genotypes at all variants present in the combined reference panel. To do this efficiently, we split the data into subsets of 500 samples, and split the genome into a total of 1456 chunks of 2Mb each. We ran IMPUTE2 in each chunk and sample subset specifying a buffer region of 500kb and an effective population size of 20,000. Finally, we used QCTOOL to merge imputation chunks across sample subsets and across chunks, encoding the results in BGEN format ^84^. For post-imputation analysis we additionally merged all impute ‘info’ files across chunks and sample subsets into a single file.

We also repeated the above process using the 1000 Genomes Phase III reference panel (as available from the IMPUTE2 website 3^rd^ August 2015), using the same settings throughout.

#### 3.4. Assessment of imputation performance

To assess per-variant imputation performance we focused on ‘type 0 r^2^’ (which measures correlation between input genotypes and re-imputed genotype dosages), which we refer to here as accuracy. We plotted mean accuracy in 1% minor allele frequency (MAF) bins for each subset of samples imputed (**Figure S1**). We also plotted the proportion of variants meeting a given accuracy threshold for lower-frequency (<10% MAF) and higher-frequency (10-50% MAF) variants. Under the assumption that untyped variants behave similarly to typed variants, these plots suggest that a substantial proportion of variation in the genome is imputed to high accuracy using this panel.

To compare imputation performance between panels we additionally computed the mean difference in accuracy, within minor allele frequency bins, for each imputation sample subset (**Figure S1**).

We also used the per-sample imputation results output by IMPUTE2 to compare imputation performance across ethnic groups. IMPUTE2 outputs an accuracy measure (also called type 0 r^2^, which we refer to as per-sample accuracy) for each sample for each imputation chunk, reflecting the correlation between input genotypes and re-imputed genotype dosages across variants within each chunk. We plotted the distribution of per-sample accuracy, averaged across chunks, for samples in each of the largest ethnic groups in our data (**Figure 1**).

#### 3.5. Visualisation of study and reference panel haplotype similarity

To further understand the effects of additional African reference panel samples on imputation performance, we constructed a joint dataset consisting of phased haplotypes at all 17,960 study individuals and all 3,046 reference panel individuals, subsetted to the overlapping set of 1,492,601 SNPs. For each pair (S,P) of a study panel haplotype S and a reference panel population P, we computed the proportion of 1Mb chunks such that the closest reference panel haplotype to S lies in P. Proximity was measured by absolute number of differences with ties broken by randomly choosing one of the closest haplotypes. We averaged the proportions over samples in each study ethnicity (**Figure 1c**).

#### 3.6. Imputation of classical HLA alleles

We used HLA*IMP:02 to impute HLA classical alleles for all study samples. We based imputation on the unphased, QCd set of genotype data across populations in the region chr6:28Mb-36Mb. This version of HLAIMP outputs imputed diploid allele calls, and posterior probabilities, for alleles at 2-digit and 4-digit resolution, and uses an allele naming scheme that is similar to the pre-2010 IPD-IMGT/HLA naming convention.

For downstream analysis, we split HLA*IMP output into per-allele posterior probabilities. Specifically, for each allele we computed the posterior probability of zero, one or two copies of the allele, against all other alleles at the locus, from the HLA*IMP output. We encoded this data in the GP field of a VCF file with one row per allele. This functionality may be generally useful and we implemented it in QCTOOL. For later reference, we assigned each allele to the midpoint of the corresponding gene.

#### 3.7. Imputation of glycophorin structural variants

We used IMPUTE to impute genotypes into our previously described reference panel in the glycophorin region on chromosome 4^14^. Specifically, we based our imputation on the phased set of haplotypes, but to avoid potential issues with phasing in this region of copy number variation and segmental duplication we treated these genotypes as unphased. We then ran IMPUTE2 in 2Mb chunks across a 10Mb region surrounding the glycophorins, specifying 1000 reference panel haplotypes, a 500kb buffer region, an effective population size of 20,000, and using the HapMap combined recombination map.

### 4. Association testing

#### 4.1. Association with severe malaria status

We used SNPTEST (See **Online Resources**) to test each imputed variant for association with severe malaria status in each study population. Specifically, for each population we ran SNPTEST for each of the 2Mb chunks output by imputation, including either 0, 2, 5 or 10 population-specific PCs as covariates, and under additive, dominant, recessive, heterozygote and general models of association. For the main analyses presented here we used the versions based on including 5 PCs. A total of 17,056 samples had principal components and assigned case/control status, and were included in association analyses. We tested under additive, dominant, recessive and heterozygote inheritance models.

#### 4.2. Association with severe malaria subphenotypes

To test association with the main severe malaria subphenotypes in our data - namely cerebral malaria (CM) and severe malarial anaemia (SMA), and other severe malaria cases (OTHER) - we extended SNPTEST to perform maximum likelihood inference for multinomial logistic regression. A full description of this method is presented in **Text S6**. In brief, in this framework, the likelihood is parameterized by the log-odds of each case phenotype (CM, SMA or OTHER) relative to the baseline phenotype (CONTROL), for the genotypic predictor along with other covariates. The method produces maximum likelihood estimates of these parameters, and estimated parameter standard errors and covariances. Overall evidence for association can then be assessed by computing a likelihood ratio test statistic, and associated P-value, against the null model where all genetic effect size parameters are zero. Additionally, for each case phenotype we compute a Wald test statistic and P-value against the null model that the genetic effect on that phenotype is zero.

Of the 17,056 samples included in association testing, 537 had no subphenotype assignment or were assigned as having both CM and SMA phenotypes. For simplicity we excluded these samples from the genome-wide test against subphenotypes. We tested under additive, dominant, recessive and heterozygote inheritance models.

#### 4.3. X chromosome tests

We additionally tested variants on the X chromosome using a logistic regression model including sex as a covariate, and treating male hemizygote genotypes in the same way as homozygous females.

### 5. Meta-analysis of association results

#### 5.1. Overview of meta-analysis

To efficiently meta-analyse the association results, we further developed our software package BINGWA (see **Online resources**). BINGWA is written in C++ and is available at http://www.well.ox.ac.uk/~gav/bingwa/#overview. BINGWA takes a list of SNPTEST files, along with options that control variant filtering and output, and computes meta-analysis results using both frequentist and bayesian methods.

Per-population results for each variant were included in meta-analysis if the minor allele count (or for nonadditive tests, the minor predictor count as defined below) was at least 25, the IMPUTE info was at least 0.3, and no issues were reported with model fitting. For nonadditive tests we defined the minor predictor count as the minimum of the expected number of individuals having the effect genotype (e.g. AB or BB for dominant model of the B allele; BB for recessive model etc.) and the number having the baseline genotype (e.g. AA for a dominant model of the B allele; AA or AB for recessive model etc).

#### 5.2. Frequentist meta-analysis

For each case/control or subphenotype test, we computed a frequentist fixed-effect meta-analysis estimate β_meta_, its variance-covariance matrix V_meta_, and overall meta-analysis P-value as described in **Text S7**. For subphenotype tests, β_meta_ is 3-dimensional (corresponding to joint estimates for cerebral malaria, severe malarial anaemia, and other severe malaria cases). We additionally computed a Wald test P-value for each estimated parameter.

#### 5.3. Bayesian meta-analysis

To assess evidence for association under a more flexible set of models of association, we used a Bayesian meta-analysis framework similar to that described previously ^7,9,10^ In brief, we implemented Bayesian inference using the *asymptotic* or *approximate Bayes factor* approach ^85^. This approach treats the observed effect sizes (as estimated by logistic or multinomial logistic regression in each population) as arising from a set of true population effect sizes, together with estimation noise that is represented by the standard errors and parameter covariances estimated in each population. The ‘true’ population effects are modeled as being drawn from a multivariate normal distribution with mean zero and a prior covariance matrix *Σ* which is chosen to represent a desired model of true effects. We write this covariance in the form

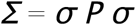

Here *P* is a correlation matrix specifying the prior correlation in true effect sizes between populations and/or between subphenotypes, and *σ* is a scalar (or, in the general case, a diagonal matrix) which determines the prior variance of effect sizes, and thus the overall magnitude of modeled true effects.

Given the vector of per-population estimates *β* and the parameter covariance matrix *V*, a Bayes factor for the model encoded by *Σ* can now be computed as a ratio of multivariate normal densities

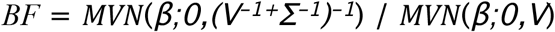

Different choices of *Σ* correspond to different assumptions about the underlying true effect sizes. The true pattern of effects is unknown, so we assess evidence under a collection of plausible assumptions *Σ_1_, Σ_2_*, … by computing the model-averaged Bayes factor

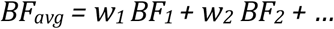

in which *BF_i_* is the BF computed using prior covariance *Σ_i_*, and the prior weights w_i_ are chosen to sum to 1.

The specific choices of prior correlation matrix *P* and weights *w* used for our primary analysis are described in subsequent sections and in **Table S9**. Unless otherwise specified, for each choice of *P* we computed the BF for that model as an equally-weighted average over four values of *σ*– namely 0.2, 0.4, 0.6 and 0.8.

#### 5.4. Mode of inheritance

We repeated all analyses under additive, dominant, recessive and heterozygote modes of inheritance with prior weights 0.4, 0.2, 0.2 and 0.2 respectively.

#### 5.5. Population effects

For case/control tests we considered the following models of effects across all populations:

– *fixed effects* (all off-diagonal entries of *P* set to 0.99)
– *correlated effects* (all off-diagonal entries of *P* set to 0.9)
– *independent effects* (all off-diagonal entries of *P* set to 0)
– *structured effects* (*P* estimated from the correlation in allele frequencies genome-wide)

We placed a total of 60% prior weight on fixed and correlated effects and 4% prior weight on each of independent and structured effects.

In addition to effects across all populations, we included models of effect restricted to population subgroups. Specifically we focused on the following groupings:

- West African populations (Gambia and Mali, or Gambia, Mali, Burkina Faso, and Ghana)
- West and central west African populations (Gambia, Mali, Burkina Faso, Ghana, Nigeria and Cameroon)
- Central west African populations (Burkina Faso, Ghana, Nigeria and Cameroon)
- Central and east African populations (Nigeria, Cameroon, Malawi, Tanzania, and Kenya)
- East African populations (Malawi, Tanzania, and Kenya)
- All African populations (Gambia, Mali, Burkina Faso, Ghana, Nigeria, Cameroon, Malawi, Tanzania and Kenya)
- Non-African populations (Vietnam, PNG)

These subsets are chosen to correspond to geographical groupings, as well as to groups apparent along the first principal component of African populations (**Figure 1e**), and such that no group has fewer than 2,000 samples. We placed 4% prior weight on each of the eight groups, for a total weight of 1 (**Table S9**).

#### 5.6. Subphenotype effects

We placed 80% of prior mass on case/control effects, but also considered effects that vary across subphenotypes (CM, SMA and other SM), as well as effects that are only present for only two or one of the subphenotypes (**Table S9**). Specifically, we considered:

– Effects on all three subphenotypes (between-phenotype entries of *P* set to 0.9 or 0).
– Effects on two of three subphenotypes (*σ* set to 0 for one phenotype, between-phenotype entries of *P* set to 0.9 or 0).
– Effects restricted to one subphenotype (*σ* set to 0 for two phenotypes)

We assigned these categories prior weights of 8%, 6% and 6% respectively. To avoid spurious results, when conducting subphenotype meta-analysis we assumed that effects were fixed across populations. Specifically, between-population within-phenotype entries of *P* were set to 0.99, and between-population between-phenotype entries of *P* were set to 0.99 times the between-assumed phenotype correlation specified above.

#### 5.7. Interpretation of the bayesian model average

The results of using the set of models and prior weights described above to compute *BF_avg_* are shown in **Figure 2**. Conditional on this set of prior assumptions, the Bayes factor can be interpreted directly in terms of a prior odds of association as

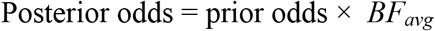

In the absence of other information about a given genetic variant, plausible values of the prior odds might be in the range 10^−5^−10^−7^ as described previously^15,86^. For a Bayes factor of 1×10^5^ this would therefore give posterior odds of association between one and one-hundredth (i.e. posterior probability of 1-50%). This computation thus reflects our belief about the top signals in our data, namely, that there is overwhelmingly strong evidence for signals at *HBB, ABO, ATP2B4* and at the glycophorin region (for which *BF_avg_* >5×10^7^), good evidence at *ARL14* and rs62418762 (*BF_avg_* > 5×10^6^), and some evidence at further loci among the top twelve (*BF_avg_* > 1×10^5^). Except for the association at *ARL14*, these observations are further reinforced by the replication data (**Figure 2c**).

Given the assumptions underlying *BF_avg_*, the relative evidence for different models can also be interpreted as in **Figure 2**. For example, since the Africa-only model has prior weight 0.04 * 0.8 = 0. 032 (averaged over mode of inheritance), the observation of 90% posterior probability on an “Africa-only” model at rs8176719 (**Figure 2**) implies that the Bayes factor for this model must be around 280 times larger than under any other model. Indeed, under a dominance model of case/control effects, the computed Bayes factors are 4.5×10^17^ and 1.2×10^20^ for effects across all populations and across Africa only, a relative difference of 265. (The Bayes factor for fixed effects across Africa and Vietnam combined is 2.6×10^19^; though we did not include this model in *BF_avg_*).

To allow further assessment of the dependency of *BF_avg_* on the model assumptions, the full metaanalysis output produced by BINGWA includes

– A full set of BFs, computed under a larger set of models, including those making up the *BF_avg_*.
– The model with the highest BF.
– The model with the highest posterior weight (i.e. highest BF after the weighting), and the model second highest posterior weight.
– A per-population BF reflecting the evidence from each individual population. (As for *BF_avg_* these are computed by model averaging over parameters *σ = 0.2,0.4, 0.6, 0.8*. For subphenotype tests we assume independence between effects in different subphenotypes).
– Meta-analysis effect size estimates and standard errors, computed under the frequentist fixed-effect model, with associated P-values.
– A summary of the set of populations and number of samples which were included in metaanalysis.
– Additional information, including genotype counts and INFO scores, taken from the SNPTEST result files.

Together these quantities allow a detailed assessment of the underlying signals. In our implementation, BINGWA was used to store meta-analysis for each mode of inheritance in a sqlite database, with results indexed by variant position and identifiers. We used sqlite features to construct the final meta-analysis.

#### 5.8. Bayesian replication analysis

The Bayesian framework outlined above naturally extends to a discovery/replication setting; this is described further below and in **Text S5**.

### 6. Selection, annotation, and interpretation of associated regions

#### 6.1. Variant filtering

The combined meta-analysis results contain many variants of low frequency or low imputation quality, potentially leading to spurious association results. To identify a set of variants with high-quality evidence, we first filtered based on minor allele count and imputation quality as follows. In each population we computed an effective minor allele count by the formula

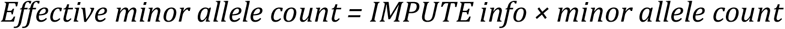

where the ‘minor allele count’ refers to the number of the less frequent allele seen in that population. For imputed variants, this is the expected number given the imputed genotype distribution. We then formed an overall effective minor allele count (EMAC) by summing this quantity across all eleven populations in our study (or, for variants where individual populations were omitted from meta-analysis, across all populations included in the meta-analysis.)

For downstream analyses we considered all variants with EMAC >= 250. This is a relatively relaxed criterion: for example, for a variant with IMPUTE info=1 in each population, this corresponds to an overall minor allele frequency of 250/(2*17056) = 0.75%. Using the combined genome-wide reference panel, a total of 26,035,208 variants passed this filter.

#### 6.2. Definition and annotation of association regions

We ranked the filtered list of variants by *BF_avg_* and used inthinnerator (see **Online resources**) to define a set of ‘lead’ variants and association regions. Inthinnerator works by iteratively picking the variant with the highest rank and excluding all other variants in a recombination interval around them. We specified a recombination interval of 0.125cM plus a margin of 25kb on either side of each picked variant, with distances determined by interpolating the HapMap combined recombination map.

Association regions around *HBB* and around the glycophorin gene cluster on chromosome 4 are especially extensive. We treated these regions specially, defining them as the regions chr11:3.5Mb-6.5Mb and chr4:143.5Mb-146Mb respectively based on visual inspection of the association signal, and we excluded these regions from the inthinnerator region definition process. A total of 95 regions defined in this way contained a variant with *BF_avg_* > 1,000. These regions are listed in **Table S2**.

#### 6.3. Visual assessment of signals of association

We created a set of plots showing the evidence for association in each region of association. Specifically, we created ‘hit plots’ showing:

– The association signal (log10 *BF_avg_*), colouring points by LD with the lead variant computed using the African reference panel haplotypes, and annotating variants that were directly typed and included in our phased set.
– Recombination rates from the HapMap combined recombination map.
– Regional genes and pseudogenes, taken from the UCSC Genome Browser refGene track downloaded on 9^th^ June 2016.
– A set of annotations reflecting functional information (described further below).

For each lead variant we created forest plots showing, for each mode of inheritance,

– the effect size estimate and confidence interval estimated in each population,
– the number of samples, variant frequency and IMPUTE info in each population
– the frequentist meta-analysis results
– a bar plot summarising the Bayesian meta-analysis results.

As described above, we also produced and inspected cluster plots of typed variants with evidence of association.

To annotate hitplots with functional information, we considered:

– Variants with impact rating ‘MODERATE’ or ‘HIGH’ according to Variant Effect Predictor ^26^ as described below.
– Variants previously reported as associated with traits in the NHGRI/EBI GWAS catalog ^37^ and a previously reported GWAS of haematopetic traits ^35^.
– Variants previously identified as having evidence for ancient balancing selection^65^
– RNA expression levels, transcription factor binding sites, and chromatin state in erythrocyte precursors and other cells^28–30^.
– Topologically associated domains ^87^
– Reported eQTLs across cell types ^31–33^

#### 6.4. Prediction of variant functional consequence

We used Variant Effect Predictor ^26^ from Ensembl tools release 75 to predict the functional consequence of variants in each association region. Specifically, we used our meta-analysis database to generate a VCF file containing all typed and imputed variants within each association region, and ran VEP using the –everything flag. We then included VEP results into the combined meta-analysis database. In addition to functional annotation, VEP has the useful property that it returns common identifiers (e.g. dbSNP ‘rs’ IDs) for variants that may have specialized IDs in the imputation reference panel.

#### 6.5. Processing of RNA-seq data from erythrocyte precursors

We obtained raw RNA-seq data from erythrocyte precursors ^30,88^ from the Gene Expression Omnibus using the sra toolkit. In total data on eight cell types was processed: CD34+, BFU-E, CFU-E, early basophilic, late basophilic, proerythroblast, polychromatic, and orthochromatic erythrocyte progenitor cells, with three replicates of each^30,88^. We also downloaded data on 12 fetal and 12 adult experimentally differentiated erythroblasts reported by Lessard et al^31^. To process these data, we extracted FASTQ files from the SRA archives and mapped reads to the build 37 reference sequence using TopHat v2.0.14, using the Gencode v19 release of the human reference sequence and gene annotations. For the Lessard et al data we included the options --library-type fr-firststrand and --microexon-search to match the original processing^31^.

We also downloaded RNA-seq data from circulating erythrocytes ^44^ from the Gene Expression Omnibus. We mapped these data to the build 37 reference sequence using bwa-mem ^74^ and marked duplicates using Picard (https://broadinstitute.github.io/picard).

To visualize coverage profiles across *ATP2B4*, we restricted attention to reads with mapping quality at least 50, and used bedtools genomecov^89^ to compute coverage across the gene and across the genome. We then plotted per-base coverage divided by the total coverage across ATP2B4 exons, summed over samples within each dataset and developmental stage (**Figure 5**).

To compute eQTL results in *ATP2B4* exons in the 24 samples from Lessard et al^31^, we computed FPKM in each exon as

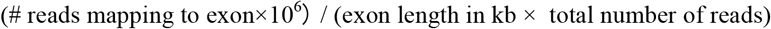

We then used a linear model to compute residual FPKM after regressing out the developmental stage (fetal or adult) ^31^. For comparison across exons we additionally standardised this residual FPKM across samples (i.e. by subtracting the mean and dividing by the standard error across samples). We then computed eQTL results by fitting a linear model of genotype on the standardized, residual FPKM values.

#### 6.6. Processing of normalised RNA-seq coverage profiles

We downloaded normalized RNA-seq coverage profiles for 56 cell types, representing 12 tissues and the ENCODE cell lines, from the roadmap epigenomics browser^28^ at http://egg2.wustl.edu. We used the stranded normalized coverage files.

For visualization of coverage profiles (**Figure 5**) we combined coverage across strands by summing the two strands. To summarise coverage across tissues, we used the tissue assignments in the “jul2013.roadmapData.qc - Consolidated_EpigenomeIDs_summary_Table.csv” file, and computed the maximum per-base normalized coverage across all cells within each tissue. We treated ENCODE cell lines as a single group here, except that we treated the K562 cell line separately as it has an expression programme similar to erythroid cells.

### 7. Generation and analysis of replication data

#### 7.1. Selection and genotyping of SNPs for replication

For validation of imputed results and additional replication, SNPs were genotyped on the Sequenom® iPLEX Mass-Array platform (Agena Biosciences, Hamburg, Germany) as described elsewhere ^90^. We used a preliminary version of the analysis described above to select association regions to target for replication. In brief, we identified the 100 regions with the most evidence for association under a case/control model of association, using inthinnerator to identify lead variants within regions. In each region we considered up to 5 SNPs for replication, including the lead SNP and others with substantial evidence for association. We further ranked regions using *BF_avg_* restricted to non-case/control models of association and selected additional SNPs making up to 5 in each region, where not already selected by the case/control scan.

We designed primer sets as described previously ^7^, omitting variants in regions previously assayed ^7,10^ Altogether we designed two multiplexes targeting 76 variants. We used these multiplexes to genotype all samples collected in the project ^7^, including those included in discovery and additional replication samples. Three variants failed genotyping (as determined by > 20% rate of missing genotypes across samples) and were excluded from downstream analysis, leaving data for 73 variants in 35 genomic regions. Genotype calls for these assays, along with data for the SNPs previously assessed ^7,10^ were combined in a single database for downstream processing.

#### 7.2. Curation of Sequenom results

We processed Sequenom genotypes as follows. Data was available for a total of 38,155 samples, most of whom represent severe malaria cases and population controls. All samples have a gender assignment by genotyping of amelogenin SNPs^7^. A small number of individuals were typed from multiple samples. For these individuals, we first removed samples with discrepancies between the clinical record of gender and the amelogenin-based gender assignment, and then merged genotypes across the remaining samples by taking a consensus call across the repeat typing (i.e. treating discordant calls as missin). Finally, we treated samples as low quality, and excluded from downstream analysis, if greater than 10% of genotype calls were missing across variants for which an attempt to type the sample had been made.

#### 7.3. Validation of imputation results

For each imputed lead variant, we computed the correlation between imputed genotypes and Sequenom-typed genotypes, for all variants within 100kb of the lead variant, within each population and across samples. We treated signals as potentially replicable if a SNP with r^2^>0.75 with the lead imputed SNP was present (**Table S2**).

#### 7.4. Frequentist replication analysis

We used SNPTEST to compute association test results based on the Sequenom genotypes, in discovery samples, replication samples, and across all samples in each population, including ethnic group as a covariate. We then used BINGWA to meta-analyse these results. A commonly-used frequentist assessment of replication is to look for a nominally significant one-tailed P-value in the direction of effect observed in discovery. These values are reported in **Table S2** for the regions identified in discovery with *BF_avg_* > 1,000, where a sequenom tag SNP exists.

#### 7.5. Bayesian replication analysis

We extended the Bayesian analysis described above to a full discovery / replication analysis. A full description of this method can be found in **Text S5**, but in brief, this analysis produces a discovery BF (*BF_avg_* as described above), an overall BF across discovery and replication samples (*BF_overall_*), and a replication BF (*BF_replication_*) which is computed under the posterior effect size distribution learnt from discovery samples. These quantities satisfy

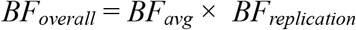

Intuitively, BF_*replication*_ measures the evidence for association under the set of models and effect size distribution learnt from discovery samples.

In practice, requiring replication effect sizes to match the distribution learnt from discovery may be too restrictive - for example this could happen if discovery effects sizes at chosen SNPs are affected by Winner’s curse, or because the distribution of phenotypes in replication samples differs from that in discovery. We therefore assessed replication data assuming true effect sizes in discovery and replication samples had a correlation of 0.9. The effects of this assumption are discussed in **Text S5**.

#### 7.6. Replication evidence at rs62418762

We used both frequentist and Bayesian analyses to assess overall and replication evidence at rs62418762 (**Figure 3**), as described above. Discovery data suggests a positive (i.e. risk) effect of the nonref allele at rs62418762 on both CM and SMA susceptibility. We therefore also computed a replication P-value for the alternative model ‘CM and SMA effects are positive’ (**Figure 3**). To do this, we simulated 10,000,000 parameter values under the null model, using a 2-dimensional normal distribution with mean zero and covariance set to the estimated covariance matrix of the replication loglikelihood from meta-analysis. We estimated the P-value as the proportion of simulated parameter values which lay in the positive quadrant and were more extreme than the observed quantities (in the sense that they are points at lower density than the observed effect sizes under this normal distribution.)

### 8. Modelling effects at replicating loci

We fit a joint model of effect of the 5 replicating variants on CM, SMA and nonspecific cases, and on samples having both CM and SMA phenotypes. Additionally, two other variants (rs33930165, encoding haemoglobin C (HbC), and rs8176746, which is indicative of A/B blood type, were included as they have been previously implicated in malaria susceptibility ^7,51^. We fit the joint model across all populations, using the nnet package in R and including 5 principal components from each population and the study population indicator as covariates. We treated effects as additive, except that variants with a clearly nonadditive effect (as in **Figure 2c**) were encoded using the predicted protective dosage. We used the estimated effect on CM to determine the protective allele and for this analysis we assumed effects were fixed across populations. Results are shown in **Figure 4a**. This analysis suggests little evidence of association with rs8176746, indicating that observed effects at this SNP are likely due to linkage disequilibrium with the O blood group mutation rs8176719.

We used control samples to compute the frequency of the protective dosage (or the allele frequency for variants with additive effects) across populations. To visualize frequencies, we computed the minimum, maximum and mean frequency for each variant across populations, and plotted frequencies as circles with width proportional to the maximum frequency (**Figure 4b**).

To visualise the potential for interaction between the variants, we computed the combined genotype for each sample across these variants, omitting rs8176746 (**Figure 4c**). Combined genotypes with less than 50 samples were collapsed into a single, ‘other genotype’ category. We refit the model across populations using these genotypes as predictors and plotted the estimated effect sizes and confidence intervals, against the expectation given the estimates from individual variants and the assumption that variants contribute independently to the log-odds of disease.

### 9. Mendelian randomization with blood cell traits

We conducted mendelian randomization (MR) analysis (which, under restrictive assumptions, can provide evidence of a causal link between two traits^91^) using summary statistics for 36 red blood cell (rbc), white blood cell (wbc), and platelet traits previously published ^35^. To do this, we treated each rbc, wbc or platelet trait separately. For each trait T, we assumed a model of an effect of *T* on SM, such that a unit increase in *T* increases the log-odds of SM by a quantity *ρ*. If this is the case then for any genetic variant affecting *T* we should have

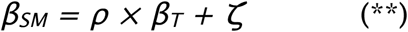

where *β_T_* and *β_SM_* denote the effect of the variant on *T* and on severe malaria, and *ζ* represents any additional effect of the variant on SM, over and above the effect through *T*. Under the assumptions traditionally underlying mendelian randomization analysis ^91^, *ζ=0* and we assumed this throughout.

We used a Gaussian likelihood formulation of MR^92^ under which the effect size estimates (*β’_T_* and *β’_SM_*) are assumed distributed around the true effects according to the estimated standard errors, as:

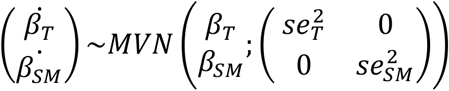

and true effects are assumed to follow:

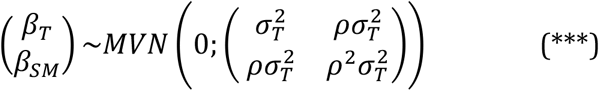

The latter equation reflects equation (**) when *ζ=0*, and the assumption that effects follow a Gaussian distribution.

To estimate ρ we focused on the 2,706 ‘sentinel’ variants identified as associated with blood traits by Astle et al^35^. A subset of 2,130 of these variants were included in the meta-analysis in our study after filtering by effective minor allele count. We removed 89 variants lying in any of the 95 regions of association containing a variant with *BF_avg_* > 1×10^3^ in our discovery analysis (including the *ATP2B4* region). We treated the remaining variants as independent, and estimated ρ and σ^2^_T_ by minimizing (***) using the optim() function in R. To compute a P-value for model fit, we refit the model assuming no correlation (ρ=0) and computed the likelihood ratio P-value.

We visualized effect size and the model fit by plotting the effect size estimates on *T* and on SM across variants, overlaid with a line indicating the estimated value of *ρ* and its 95% confidence interval. To avoid visualization problems stemming from large estimates at variants with high standard errors, we used effect size estimates regularized using a *N(0,0.1)* prior in plots. These estimates are plotted in **Figure 5h** (for mean corpuscular haemoglobin) and **Figure S12** (for all traits tested).

For *T*=mchc, our estimate of effect variance was *σ^2^_T_* = 0.013^2^. We plotted the empirical distribution of observed effects overlaid with the Gaussian density and observed some evidence that effects are overdispersed relative to the Gaussian distribution with this variance. To verify that this did not adversely affect results, we refit the model assuming a fixed prior value *σ^2^_T_* = 0.1^2^, with similar results (ρ=-0.13, *P* = 7×10^−3^).

### 10. HLA region analysis

#### 10.1. Association analysis

For each imputed HLA allele we tested for association and conducted meta-analysis as described above for genome-wide SNPs and indels.

#### 10.2. Comparison with previous data on West African populations

For each 2-digit HLA allele in our HLA*IMP-imputed data in Gambia, we computed the corresponding antigen frequency as the proportion of individuals heterozygote or homozygote for the allele. We plotted these frequencies in control samples from each Gambian ethnic group against previously published HLA Class I antigen frequencies in the same populations (**Table 1** of Hill et al ^1^).

We noted that a pair of alleles were particularly discordant (the B*15 allele, which is at relatively high frequency in our imputation and the B70 antigen, which is at relatively high frequency in the serotyped data. We used the IPD-IMGT/HLA dictionary to confirm that the B70 antigen is expressed by B*15 alleles. We therefore computed the combined B70/B15 frequencies in **Figure 6b**. For each ethnic group we then computed squared correlation between imputed and serotyped antigen frequencies.

#### 10.3. HLA typing in 32 Gambian children

31 Gambian case samples for which both parents have previously been genotyped (EGA dataset EGAD00000000019) were selected based on available DNA quantities. These samples were typed at 11 HLA gene loci through exon sequencing. Sequenced regions included exons 2-4 of HLA-A, - B, -C, -DPB1; exons 2-3 of HLA-DUPA1, -DQA1, DQB1 and DRB1, and exon 2 of HLA-DRB3, -DRB4, and -DRB5. Typing was performed by the Accredited Tissue Typing Laboratory at Addenbrooke’s Hospital, Cambridge University Hospitals NHS Foundation Trust, using the proprietary uTYPE software version 7 (Fisher Scientific. Pittsburgh, USA). The list of possible ambiguous calls were minimised by using the ‘allele pair’ export function in this software, which lists all possible and permissible allele pair possibilities for each locus for each individual. Alleles were defined using the IMGT/HLA Release: 3.22.0 2015 October 10. Best-call allele pairs for each locus in each individual were determined based on local guidelines prioritising alleles that were “Common and Well-Documented” (CWD).

### 11. Analysis of Glycophorin structural variants

We tested for association and conducted meta-analysis for glycophorin region variants in the same way as described above for other variants. Four CNVs were imputed with reasonable frequency and imputation quality (DUP1; the glycophorin B deletions DEL1 and DEL2; and DUP4).

Glycophorin region SNPs appear in our meta-analysis results twice - once using the genome-wide combined reference panel, and once based on this imputation of glycophorin variants. In our main presentation for consistency we refer to the genome-wide panel imputation results, except for glycophorin CNVs which are imputed from the glycophorin panel.

### 12. Analysis of polygenicity of severe malaria

We aimed to assess the degree to which additional polygenic effects explain severe malaria susceptibility. The epidemiology of malaria differs between sub-Saharan Africa and outside, and we chose to restrict attention to the African populations in our data for this analysis. We further restricted attention to a subset of 13,088 samples having pairwise relatedness less than 0.05 (estimated in each population based on the kinship matrix used to compute principal components from phased genotypes, as described above). We used the genotypes at directly typed SNPs in the phased dataset for heritability estimation.

We used two previously published methods (GCTA^93^ and PCGC ^20^, in the implementation by Gaurav Bhatia (https://github.com/gauravbhatia1/PCGCRegression)) to estimate the heritability of SM. These methods are both based on the “infinitesimal” model of genotype-trait association, under which all variants contribute equally, in expectation, to trait variation. However, they differ in how estimation is performed. In particular, PCGC is thought to be more robust for case/control traits and we use it for our main results.

We estimated heritability based on the phased dataset in several ways:

– in each chromosome separately
– in a joint model in which all chromosome were included as separate components
– in a joint model in which the four previously confirmed association regions, and the rest of the genome, were included as separate components
– in a joint model in which regions near protein-coding genes, and the rest of the genome were included as separate components
– in a joint model in which variants in different minor allele frequency bins were allocated to different components.

For our main analysis we included an indicator of callset and 20 PCs, as computed across the 13,088 samples, as covariates. GCTA takes covariates into account directly; to adjust for covariates in PCGC, we used the --adjust-grm option in LDAK^22^ to subtract covariates from the relatedness matrix. (However, we did not further explore using the LDAK model for heritability estimation). For some analyses we also included the protective dosages of risk alleles at HbS, rs8176719 (*ABO*), rs4951377 (*ATP2B4*) and DUP4 as covariates.

We also explored including 10 or 50 PCs as covariates, with little difference in results (data not shown). We observed slightly smaller estimates of heritability using GCTA than PCGC, consistent with the previously reported theory and observations^20^. Although our populations are highly structured, estimated of per-chromosome heritability when fitting chromosomes independently or jointly were similar (**Figure S4**). This suggests that any between-chromosome correlations between variants, which can occur as a result of population structure, are being adequately controlled for by our inclusion of covariates.

### 13. Analysis of allele frequency differentiation

#### 13.1. Overview

We used the reference panel data to construct a joint derived allele count spectrum between African and non-African populations (**Figure S7**), and we used the genotypes of control samples at a set of randomly selected sites to construct an empirical model of allele frequencies across African populations (**Figures S8-9 and S11**). We then annotated variants across the genome with metrics of differentiation between continents and within Africa based on these models.

#### 13.2. Between-continent differentiation

To assess between-continent differentiation, we first computed the empirical allele count distribution between African and each non-African group G, based on all reference panel variants which had an ancestral allele assignment and which were not masked by the 1000 Genomes strict mask. Let D = D_ij_ be the matrix of allele counts, where i ranges from 0 to the number of haplotypes in G and j ranges from 0 to the number of haplotypes in African reference panel individuals. Given a derived allele x with counts a=G(x) and b=AFR(x), we then compute upper and lower ranks of the allele in G as

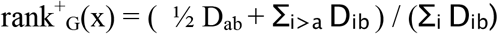

and

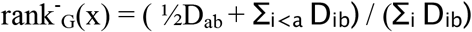

These quantities reflect the rank of the allele x among all alleles with the same African allele count, and satisfy rank^+^_G_(x) + rank^−^_G_(x) = 1. Visually, they correspond to the position of the allele in the vertical ‘slice’ through the joint derived allele count matrix (**Figure S7**).

To compute the African allele counts in D we used only samples from non-admixed African populations (i.e. excluding ACB and ASW populations).

The matrix D is computed from data on genome-wide variants. If these are primarily neutrally evolving mutations, then D forms an estimate of the distribution of allele counts under neutrality, andrank^+^_G_(x) can be interpreted as an empirical P-value reflecting the alternative hypothesis that the allele count G(x) is higher than expected, given AFR(x). Similarly we interpret rank^−^_G_(x) as an empirical P-value reflecting the alternative hypothesis that G(x) is lower than expected, given AFR(x).

For a malaria-associated variant, the malaria hypothesis suggests that natural selection will maintain the protective allele at higher frequency in populations of high childhood mortality (such as African populations in our data) due to malaria. To avoid confounding by selection of variants through GWAS, we instead condition on the African allele frequency and ask whether the allele is at lower frequency in non-endemic populations. For any variant v with derived allele x, we therefore define rank_G_(v) = rank^−^_G_(x), if x is the estimated protective allele, or rank_G_(v) = rank^+^_G_(x) if x is the estimated risk allele. We will therefore have rank_G_(v) < 0.5 if the protective allele is at below-median rank in group G among all variants with the same African allele count AFR(x). Where unclear we determine the direction of effect of x based on the effect size estimated for cerebral malaria.

#### 13.3. Within-Africa differentiation

To assess differentiation within Africa we implemented a model similar to that underlying Bayenv ^94^ but with simplifications that we now describe. The Bayenv model assumes that the vector of underlying allele frequencies in sampled populations (denoted *F*) follows a multivariate normal distribution of the form

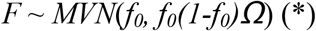

Here *f_0_* denotes the conceptual ancestral allele frequency (assumed to reflect a time point *T* before the populations from which study samples were drawn separated), and the matrix *Ω* captures the effects on allele frequency of the coancestry of populations since *T*. In particular, diagonal entries of *Ω* reflect levels of genetic drift (relative to the population at time point *T*) and off-diagonal entries reflect shared ancestry between populations.

The model above is expected to hold approximately provided *f_0_* is not too small and the levels of genetic drift are not too large^94^. Because the coancestry matrix *Ω* is assumed to be the same across all neutrally evolving variants, it can be estimated from a set of neutral SNPs across the genome. Specifically, for each variant, a vector of scaled frequencies can be computed as

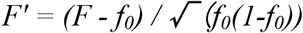

Under the model above, *Ω* can now be estimated as the covariance of scaled allele frequencies,

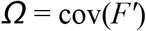

To implement this in practice, we made the following simplifying assumptions. First, we restricted attention to the 7 African study populations with > 200 control samples (namely Gambia, Burkina Faso, Ghana, Cameroon, Malawi, Tanzania, and Kenya), and assumed that the population allele frequency vector *F* is accurately estimated by the observed allele frequencies. (We used only control samples in frequency estimates). Second, we assumed that the ancestral allele frequency *f_0_* is well estimated as the mean of the per-population frequency estimates. Thirdly, we restricted attention to directly typed SNPs with mean allele frequency in the range 2-98%, and ignored the effects of allele frequencies reaching the boundary at frequency 0 or 1. We sampled a subset of 100,000 such SNPs randomly across the genome with the above properties and computed *Ω* as the empirical covariance in scaled estimated allele frequencies. The estimated matrix *Ω′* is visualized in **Figure S8**. We note that the Bayenv method avoids these assumptions by integrating over the uncertainty in ancestral and current allele frequencies^94^, and by modeling the population frequencies as bounded between 0 and 1, but requires a computationally expensive MCMC process.

In practice, systematic effects in *F′* may arise due to the selection of variants used in estimation. In addition to *Ω′* we therefore additionally estimated the mean scaled frequency, denoted *μ*.

The estimated covariance matrix *Ω* provides a model for allele frequency variation of putatively neutral alleles in the populations considered. We visualized this model by plotting the joint distribution of observed and simulated allele frequencies. Specifically, we simulated 1,000,000 variants by sampling f_0_ from the observed list of mean allele frequencies, and then sampling allele frequencies by drawing from the model (*). We then plotted the joint allele frequency distribution in the two most divergent populations, Gambia and Kenya (**Figure S8**).

We tested each typed or imputed variant against this model as follows. First, we computed the mean allele frequency and scaled allele frequency vector F′. The model above assumes that

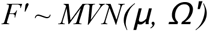

Or equivalently,

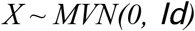

where *X = L^−1^ (F’-μ)* and *LL^t^ = Ω* denotes the Cholesky decomposition of *Ω*. In practice, the use of mean allele frequency implies that *Ω* has rank one less than the number of populations ^94^ and we therefore remove one population from the vector F′ and from *Ω* before computing this decomposition. In the results presented here, we chose to remove Cameroon, which is the most central population both geographically and within *Ω* (**Figure S8**). Under the model, the sum of squared entries of *X*, i.e. *X^t^X*, is now distributed as χ^2^ distribution with 6 degrees of freedom. We denote the corresponding P-value as P_XtX_.

**Figure S9** compares the distribution of the X^t^X test statistic against the χ^2^ distribution for typed and imputed variants at different frequencies. **Figure S11** depicts −log10 *P_XtX_* for all imputed variants with mean allele frequency in the range 2-98%.

### 14. Online resources

- QCTOOL is available at http://www.well.ox.ac.uk/~gav/qctool
- Inthinnerator is available at http://www.well.ox.ac.uk/~gav/inthinnerator
- BINGWA is available at http://www.well.ox.ac.uk/~gav/bingwas
- SNPTEST is available at https://mathgen.stats.ox.ac.uk/genetics_software/snptest/snptest.html
- More information on our study and data is available at http://www.malariagen.net.

## References

1. Hill, A.V. et al. Common west African HLA antigens are associated with protection from severe malaria. Nature 352, 595–600 (1991).

2. Robinson, J. et al. The IPD and IMGT/HLA database: allele variant databases. Nucleic Acids Res 43, D423–31 (2015).

3. Casanova, J.L. & Abel, L. The genetic theory of infectious diseases: a brief history and selected illustrations. Annu Rev Genomics Hum Genet 14, 215–43 (2013).

4. Hall, M.D. & Ebert, D. The genetics of infectious disease susceptibility: has the evidence for epistasis been overestimated? BMC Biol 11, 79 (2013).

5. Hill, A.V. Evolution, revolution and heresy in the genetics of infectious disease susceptibility. Philos Trans R Soc Lond B Biol Sci 367, 840–9 (2012).

6. Malaria Genomic Epidemiology, N. A global network for investigating the genomic epidemiology of malaria. Nature 456, 732–7 (2008).

7. Band, G. et al. Imputation-based meta-analysis of severe malaria in three African populations. PLoS Genet 9, e1003509 (2013).

8. Jallow, M. et al. Genome-wide and fine-resolution association analysis of malaria in West Africa. Nat Genet 41, 657–65 (2009).

9. Malaria Genomic Epidemiology, N., Band, G., Rockett, K.A., Spencer, C.C. & Kwiatkowski, D.P. A novel locus of resistance to severe malaria in a region of ancient balancing selection. Nature 526, 253–7 (2015).

10. Malaria GEN. Reappraisal of known malaria resistance loci in a large multi-centre study. Nature Genetics (2014).

11. World Health Organization. Guidelines for the treatment of malaria, xi, 194 p. (World Health Organization, Geneva, 2010).

12. The 1000 Genomes Project Consortium et al. A global reference for human genetic variation. Nature 526, 68–74 (2015).

13. Crosnier, C. et al. Basigin is a receptor essential for erythrocyte invasion by Plasmodium falciparum. Nature 480, 534–7 (2011).

14. Leffler, E.M. et al. Resistance to malaria through structural variation of red blood cell invasion receptors. Science 356(2017).

15. The Wellcome Trust Case Control, C. Genome-wide association study of 14,000 cases of seven common diseases and 3,000 shared controls. Nature 447, 661–78 (2007).

16. Holmberg, J., Clarke, D.L. & Frisen, J. Regulation of repulsion versus adhesion by different splice forms of an Eph receptor. Nature 408, 203–6 (2000).

17. Kaushansky, A. et al. Malaria parasites target the hepatocyte receptor EphA2 for successful host infection. Science 350, 1089–92 (2015).

18. Ahmeti, K.B. et al. Age of onset of amyotrophic lateral sclerosis is modulated by a locus on 1p34.1. Neurobiol Aging 34, 357 e7–19 (2013).

19. Li, Z. et al. Genome-wide association analysis identifies 30 new susceptibility loci for schizophrenia. Nat Genet 49, 1576–1583 (2017).

20. Golan, D., Lander, E.S. & Rosset, S. Measuring missing heritability: inferring the contribution of common variants. Proc Natl Acad Sci U S A 111, E5272–81 (2014).

21. Evans, L.M. et al. Comparison of methods that use whole genome data to estimate the heritability and genetic architecture of complex traits. Nat Genet 50, 737–745 (2018).

22. Speed, D. et al. Reevaluation of SNP heritability in complex human traits. Nat Genet 49, 986–992 (2017).

23. Mackinnon, M.J., Mwangi, T.W., Snow, R.W., Marsh, K. & Williams, T.N. Heritability of malaria in Africa. PLoS Med 2, e340 (2005).

24. Rowe, J.A. et al. Blood group O protects against severe Plasmodium falciparum malaria through the mechanism of reduced rosetting. Proc Natl Acad Sci U S A 104, 17471–6 (2007).

25. Rowe, J.A., Opi, D.H. & Williams, T.N. Blood groups and malaria: fresh insights into pathogenesis and identification of targets for intervention. Curr Opin Hematol 16, 480–7 (2009).

26. McLaren, W. et al. The Ensembl Variant Effect Predictor. Genome Biol 17, 122 (2016).

27. Consortium, E.P. An integrated encyclopedia of DNA elements in the human genome. Nature 489, 57–74 (2012).

28. Roadmap Epigenomics, C. et al. Integrative analysis of 111 reference human epigenomes. Nature 518, 317–30 (2015).

29. Xu, J. et al. Combinatorial assembly of developmental stage-specific enhancers controls gene expression programs during human erythropoiesis. Dev Cell 23, 796–811 (2012).

30. An, X. et al. Global transcriptome analyses of human and murine terminal erythroid differentiation. Blood 123, 3466–77 (2014).

31. Lessard, S. et al. An erythroid-specific ATP2B4 enhancer mediates red blood cell hydration and malaria susceptibility. J Clin Invest 127, 3065–3074 (2017).

32. GTEx Consortium. Human genomics. The Genotype-Tissue Expression (GTEx) pilot analysis: multitissue gene regulation in humans. Science 348, 648–60 (2015).

33. Westra, H.J. et al. Systematic identification of trans eQTLs as putative drivers of known disease associations. Nat Genet 45, 1238–1243 (2013).

34. Alvarez, M.I. et al. Human genetic variation in VAC14 regulates Salmonella invasion and typhoid fever through modulation of cholesterol. Proc Natl Acad Sci U S A 114, E7746–E7755 (2017).

35. Astle, W.J. et al. The Allelic Landscape of Human Blood Cell Trait Variation and Links to Common Complex Disease. Cell 167, 1415–1429 e19 (2016).

36. Gilchrist, J.J. et al. Genetic variation in VAC14 is associated with bacteremia secondary to diverse pathogens in African children. Proc Natl Acad Sci U S A 115, E3601–E3603 (2018).

37. MacArthur, J. et al. The new NHGRI-EBI Catalog of published genome-wide association studies (GWAS Catalog). Nucleic Acids Res 45, D896–D901 (2017).

38. Merika, M. & Orkin, S.H. DNA-binding specificity of GATA family transcription factors. Mol Cell Biol 13, 3999–4010 (1993).

39. Zambo, B. et al. Decreased calcium pump expression in human erythrocytes is connected to a minor haplotype in the ATP2B4 gene. Cell Calcium 65, 73–79 (2017).

40. Harrow, J. et al. GENCODE: the reference human genome annotation for The ENCODE Project. Genome Res 22, 1760–74 (2012).

41. O’Leary, N.A. et al. Reference sequence (RefSeq) database at NCBI: current status, taxonomic expansion, and functional annotation. Nucleic Acids Res 44, D733–45 (2016).

42. Flicek, P. et al. Ensembl 2014. Nucleic Acids Res 42, D749–55 (2014).

43. Consortium, F. et al. A promoter-level mammalian expression atlas. Nature 507, 462–70 (2014).

44. Doss, J.F. et al. A comprehensive joint analysis of the long and short RNA transcriptomes of human erythrocytes. BMC Genomics 16, 952 (2015).

45. Miller, L.H., Mason, S.J., Clyde, D.F. & McGinniss, M.H. The resistance factor to Plasmodium vivax in blacks. The Duffy-blood-group genotype, FyFy. N Engl J Med 295, 302–4 (1976).

46. Tournamille, C., Colin, Y., Cartron, J.P. & Le Van Kim, C. Disruption of a GATA motif in the Duffy gene promoter abolishes erythroid gene expression in Duffy-negative individuals. Nat Genet 10, 224–8 (1995).

47. van der Harst, P. et al. Seventy-five genetic loci influencing the human red blood cell. Nature 492, 369–75 (2012).

48. Dunstan, S.J. et al. Variation at HLA-DRB1 is associated with resistance to enteric fever. Nat Genet 46, 1333–6 (2014).

49. Dilthey, A. et al. Multi-population classical HLA type imputation. PLoS Comput Biol 9, e1002877 (2013).

50. Allsopp, C.E. et al. Sequence analysis of HLA-Bw53, a common West African allele, suggests an origin by gene conversion of HLA-B35. Hum Immunol 30, 105–9 (1991).

51. Hedrick, P.W. Population genetics of malaria resistance in humans. Heredity (Edinb) 107, 283–304 (2011).

52. Kwiatkowski, D.P. How malaria has affected the human genome and what human genetics can teach us about malaria. Am J Hum Genet 77, 171–92 (2005).

53. Ma, S. et al. Common PIEZO1 Allele in African Populations Causes RBC Dehydration and Attenuates Plasmodium Infection. Cell 173, 443–455 e12 (2018).

54. Opi, D.H. et al. Two complement receptor one alleles have opposing associations with cerebral malaria and interact with alpha(+)thalassaemia. Elife 7(2018).

55. Zhang, D.L. et al. Erythrocytic ferroportin reduces intracellular iron accumulation, hemolysis, and malaria risk. Science 359, 1520–1523 (2018).

56. Crawford, N.G. et al. Loci associated with skin pigmentation identified in African populations. Science 358(2017).

57. Aitman, T.J. et al. Malaria susceptibility and CD36 mutation. Nature 405, 1015–6 (2000).

58. Reyes, A. & Huber, W. Alternative start and termination sites of transcription drive most transcript isoform differences across human tissues. Nucleic Acids Res 46, 582–592 (2018).

59. Simon, L.M. et al. Integrative Multi-omic Analysis of Human Platelet eQTLs Reveals Alternative Start Site in Mitofusin 2. Am J Hum Genet 98, 883–897 (2016).

60. Brochet, M. & Billker, O. Calcium signalling in malaria parasites. Mol Microbiol 100, 397–408 (2016).

61. Timmann, C. et al. Genome-wide association study indicates two novel resistance loci for severe malaria. Nature (2012).

62. Piel, F.B. et al. Global distribution of the sickle cell gene and geographical confirmation of the malaria hypothesis. Nat Commun 1, 104 (2010).

63. Lam, K.W. & Jeffreys, A.J. Processes of copy-number change in human DNA: the dynamics of {alpha}-globin gene deletion. Proc Natl Acad Sci U S A 103, 8921–7 (2006).

64. Segurel, L. et al. The ABO blood group is a trans-species polymorphism in primates. Proc Natl Acad Sci U S A 109, 18493–8 (2012).

65. Leffler, E.M. et al. Multiple instances of ancient balancing selection shared between humans and chimpanzees. Science 339, 1578–82 (2013).

66. McManus, K.F. et al. Population genetic analysis of the DARC locus (Duffy) reveals adaptation from standing variation associated with malaria resistance in humans. PLoS Genet 13, e1006560 (2017).

67. Bycroft, C. et al. The UK Biobank resource with deep phenotyping and genomic data. Nature 562, 203–209 (2018).

68. Khera, A.V. et al. Genome-wide polygenic scores for common diseases identify individuals with risk equivalent to monogenic mutations. Nat Genet 50, 1219–1224 (2018).

69. Mahajan, A. et al. Fine-mapping type 2 diabetes loci to single-variant resolution using high-density imputation and islet-specific epigenome maps. Nat Genet 50, 1505–1513 (2018).

70. Schizophrenia Working Group of the Psychiatric Genomics, C. Biological insights from 108 schizophrenia-associated genetic loci. Nature 511, 421–7 (2014).

71. Tian, C. et al. Genome-wide association and HLA region fine-mapping studies identify susceptibility loci for multiple common infections. Nat Commun 8, 599 (2017).

72. Bhatt, S. et al. The effect of malaria control on Plasmodium falciparum in Africa between 2000 and 2015. Nature 526, 207–211 (2015).

73. Snow, R.W. et al. The prevalence of Plasmodium falciparum in sub-Saharan Africa since 1900. Nature 550, 515–518 (2017).

74. Li, H. & Durbin, R. Fast and accurate short read alignment with Burrows-Wheeler transform. Bioinformatics 25, 1754–60 (2009).

75. DePristo, M.A. et al. A framework for variation discovery and genotyping using next-generation DNA sequencing data. Nat Genet 43, 491–8 (2011).

76. Van der Auwera, G.A. et al. From FastQ data to high confidence variant calls: the Genome Analysis Toolkit best practices pipeline. Curr Protoc Bioinformatics 43, 11 10 1–33 (2013).

77. Browning, S.R. & Browning, B.L. Rapid and accurate haplotype phasing and missing-data inference for whole-genome association studies by use of localized haplotype clustering. Am JHum Genet 81, 1084–97 (2007).

78. Delaneau, O., Zagury, J.F. & Marchini, J. Improved whole-chromosome phasing for disease and population genetic studies. Nat Methods 10, 5–6 (2013).

79. Bellenguez, C. et al. A robust clustering algorithm for identifying problematic samples in genome-wide association studies. Bioinformatics 28, 134–5 (2012).

80. Astle, W. & Balding, D.J. Population Structure and Cryptic Relatedness in Genetic Association Studies. Statistical Science 24, 451–471 (2009).

81. Powell, J.E., Visscher, P.M. & Goddard, M.E. Reconciling the analysis of IBD and IBS in complex trait studies. Nat Rev Genet 11, 800–5 (2010).

82. Wigginton, J.E., Cutler, D.J. & Abecasis, G.R. A note on exact tests of Hardy-Weinberg equilibrium. Am J Hum Genet 76, 887–93 (2005).

83. Howie, B., Marchini, J. & Stephens, M. Genotype imputation with thousands of genomes. G3 (Bethesda) 1, 457–70 (2011).

84. Band, G. & Marchini, J. BGEN: a binary file format for imputed genotype and haplotype data. bioRxiv (2018).

85. Wakefield, J. Bayes factors for genome-wide association studies: comparison with P-values. Genet Epidemiol 33, 79–86 (2009).

86. Stephens, M. & Balding, D.J. Bayesian statistical methods for genetic association studies. Nat Rev Genet 10, 681–90 (2009).

87. Javierre, B.M. et al. Lineage-Specific Genome Architecture Links Enhancers and Noncoding Disease Variants to Target Gene Promoters. Cell 167, 1369–1384 e19 (2016).

88. Li, J. et al. Isolation and transcriptome analyses of human erythroid progenitors: BFU-E and CFU-E. Blood 124, 3636–45 (2014).

89. Quinlan, A.R. & Hall, I.M. BEDTools: a flexible suite of utilities for comparing genomic features. Bioinformatics 26, 841–2 (2010).

90. Malaria Genomic Epidemiology Network. Reappraisal of known malaria resistance loci in a large multicenter study. Nat Genet 46, 1197–204 (2014).

91. Davey Smith, G. & Hemani, G. Mendelian randomization: genetic anchors for causal inference in epidemiological studies. Hum Mol Genet 23, R89–98 (2014).

92. Burgess, S., Butterworth, A. & Thompson, S.G. Mendelian randomization analysis with multiple genetic variants using summarized data. Genet Epidemiol 37, 658–65 (2013).

93. Yang, J., Lee, S.H., Goddard, M.E. & Visscher, P.M. GCTA: a tool for genome-wide complex trait analysis. Am J Hum Genet 88, 76–82 (2011).

94. Gunther, T. & Coop, G. Robust identification of local adaptation from allele frequencies. Genetics 195, 205–20 (2013).

## References

7. Malaria GEN. Reappraisal of known malaria resistance loci in a large multi-centre study. Nature Genetics (2014).

8. World Health Organization. Guidelines for the treatment of malaria, xi, 194 p. (World Health Organization, Geneva, 2010).

9. Band, G. et al. Imputation-based meta-analysis of severe malaria in three African populations. PLoS Genet 9, e1003509 (2013).

10. Malaria Genomic Epidemiology, N., Band, G., Rockett, K.A., Spencer, C.C. & Kwiatkowski, D.P. A novel locus of resistance to severe malaria in a region of ancient balancing selection. Nature 526, 253–7 (2015).

11. Jallow, M. et al. Genome-wide and fine-resolution association analysis of malaria in West Africa. Nat Genet 41, 657–65 (2009).

26. McLaren, W. et al. The Ensembl Variant Effect Predictor. Genome Biol 17, 122 (2016).

42. Flicek, P. et al. Ensembl 2014. Nucleic Acids Res 42, D749–55 (2014).

43. Consortium, F. et al. A promoter-level mammalian expression atlas. Nature 507, 462–70 (2014).

57. Aitman, T.J. et al. Malaria susceptibility and CD36 mutation. Nature 405, 1015–6 (2000).

77. Browning, S.R. & Browning, B.L. Rapid and accurate haplotype phasing and missing-data inference for whole-genome association studies by use of localized haplotype clustering. Am J Hum Genet 81, 1084–97 (2007).

