## Supplementary Information for "New insights into malaria susceptibility from the genomes of 17,000 individuals from Africa, Asia, and Oceania"

### Supplementary Figures

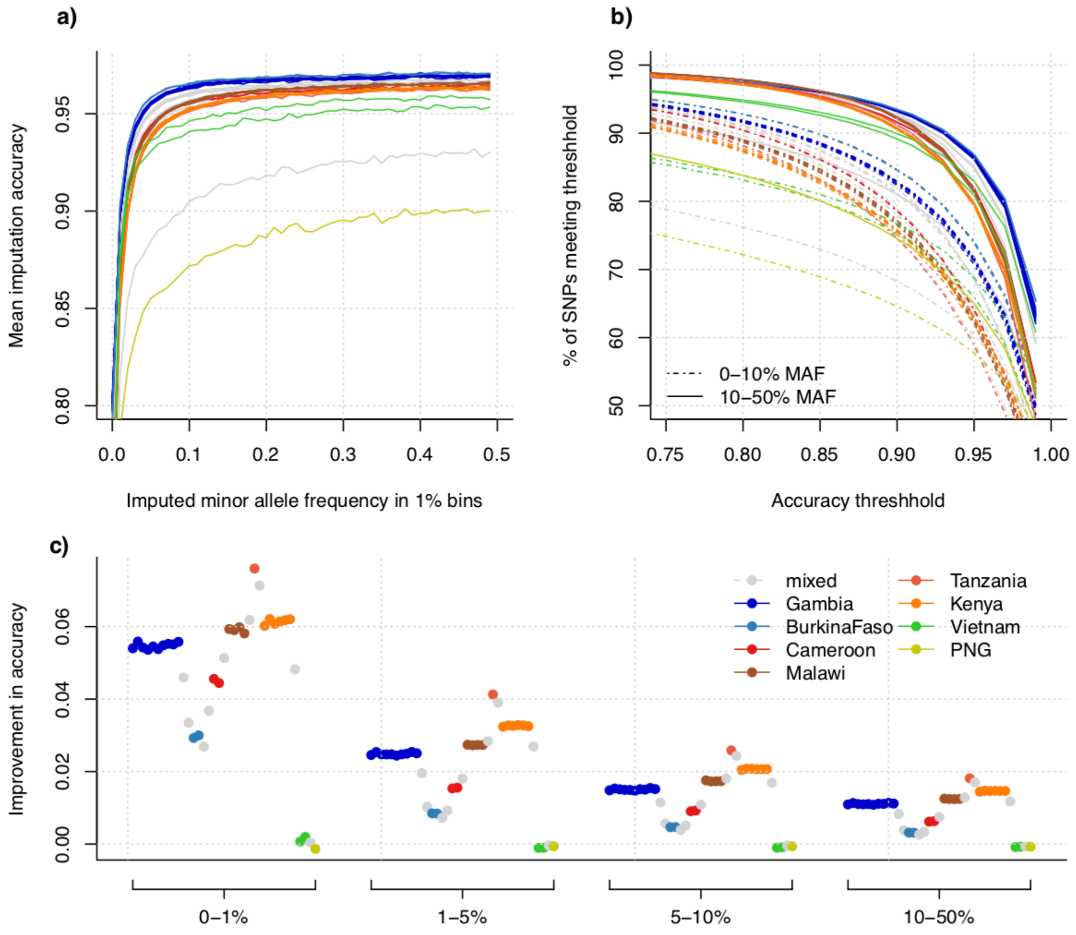

#### Figure S 1 - Detail of imputation accuracy.

a) Imputation accuracy (computed as the mean squared correlation between directly typed and reimputed genotypes, evaluated at Omni 2.5M SNPs included in the imputation) against minor allele frequency for all samples in our study. Samples were imputed in sets of 500 samples; each line represents a single sample set. Lines are coloured by population (if all samples in the set were from a single population) or grey if samples from a mixture of populations was included, according to the legend in panel c). b) The proportion of variants at or above a given imputation accuracy, for accuracies in the range 0.75-1, as computed by masking and re-imputing typed SNPs. For example, in African sample sets over 90% of common variants were reimputed with at least 90% accuracy. c) Improvement in per-variant imputation accuracy between the combined panel and the 1000 Genomes reference panel. Improvement is computed as the mean difference in accuracy for variants in the given minor allele frequency bin (x axis); each point represents a single imputation run of 500 samples.

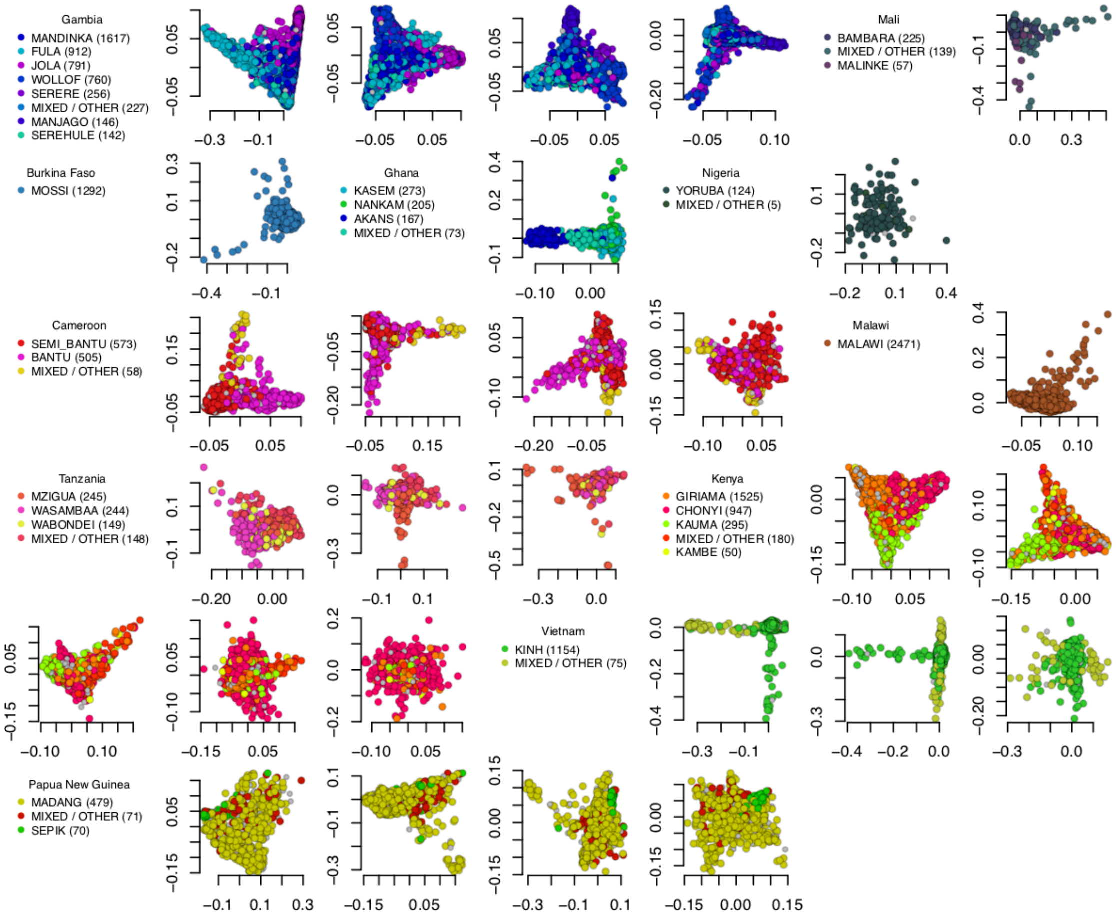

#### Figure S 2 - Principal components (PCs) across all study samples.

Selected PCs computed in each population using the phased genotype calls. For each study population, a summary of the ethnic makeup of the study samples is given, followed by a plot of the first PC (x axis) versus the second PC (y axis). For populations displaying more structure we also plot the 2dn versus 3rd PC, the 3rd versus 4th PC, etc. Colours are chosen to distinguish ethnic groups, with the largest ethnic group being set to the population colour, as shown in **Figure 1**, and other ethnic groups given a spread of hues around the population colour. In each population we plot samples in random order to avoid visual overrepresentation of specific ethnicities.

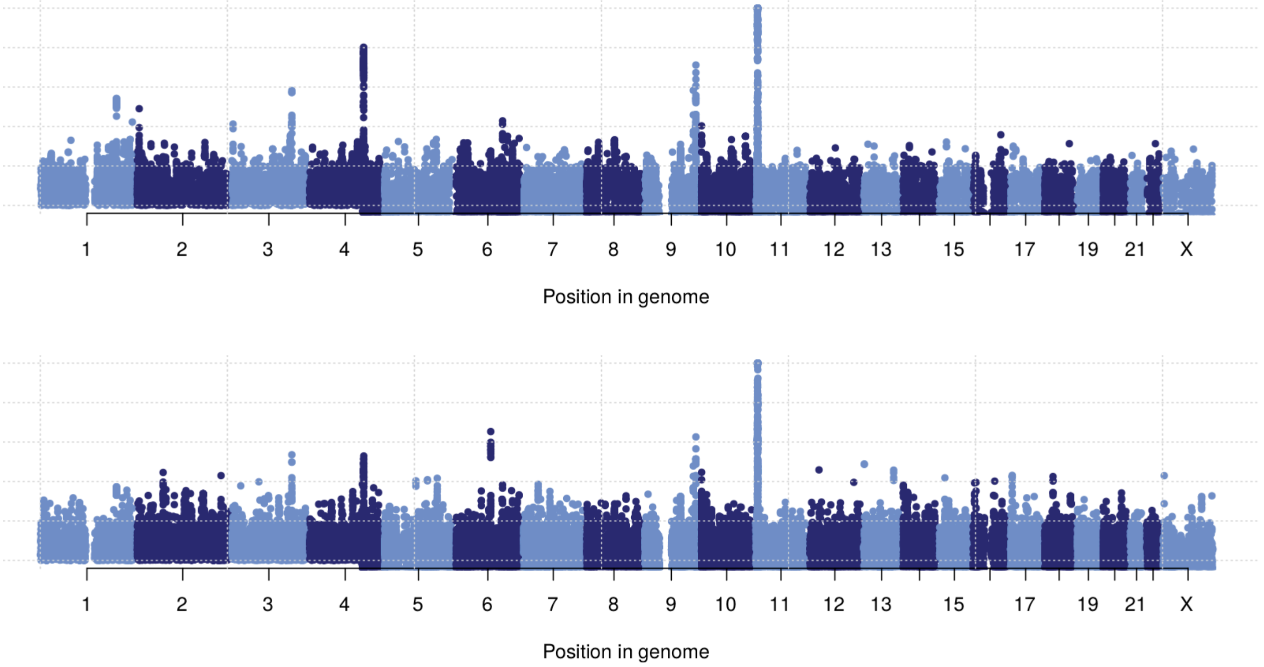

**Figure S 3 - Manhattan plot for case-control and subphenotype tests.**

**a)** -$\log_{10}$ P-value for an additive model of association of the genotype on SM status, versus controls, for each SNP included in our analysis. Effect size estimates and standard errors are computed in each study population using logistic regression (as implemented in SNPTEST) including 5 principal components of population structure. Results are then meta-analysed across populations using fixed-effect meta-analysis. **b)** -$\log_{10}$ P-value for an additive model of association of the genotype with CM, SMA, or OTHER phenotypes, relative to controls. Results are computed using multinomial logistic regression in each study population including 5 principal components. Results are then meta-analysed across populations using fixed-effect meta-analysis.

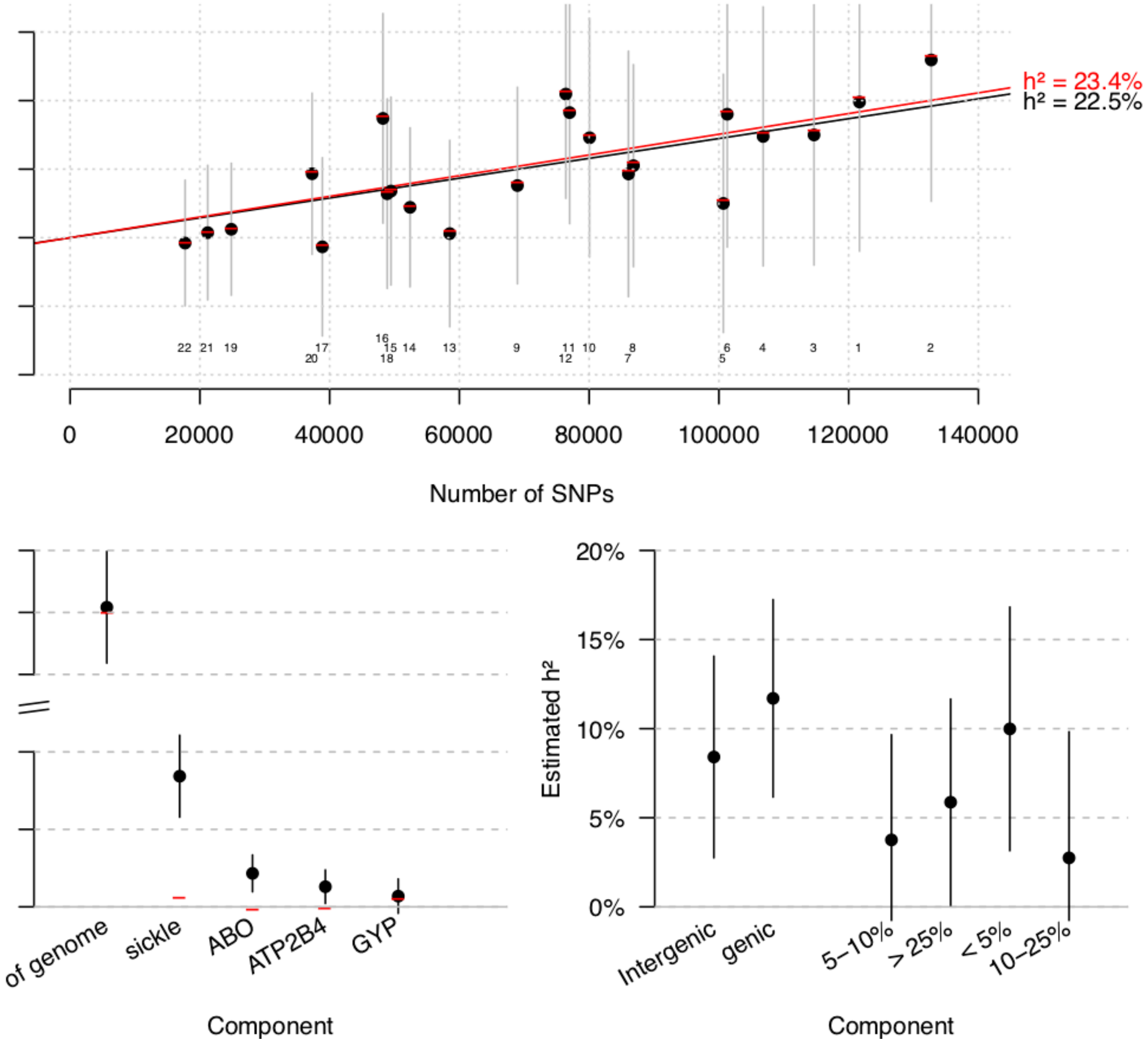

**Figure S 4 - Analysis of the heritability of SM across African study populations**

Results are based on a joint analysis of 13,038 individuals, selected to have <5% pairwise relatedness within each African study site, using the phased dataset of 1,550,514 SNPs. We included 20 PCs computed across these individuals, as well as an indicator of study site as fixed effects to control for population structure. Results are estimated using PCGC. a) Estimated heritability attributed to each chromosome when including all chromosomes jointly in the model (black points, with grey bars indicating 95% confidence intervals; small text indicates chromosome number) and when estimating for each chromosome seperately (red horizontal lines). Sloping lines indicate the overall estimated heritability for each model. A small degree of inflation is seen when using separate estimates, indicating there may be some residual confounding by population structure. b) Estimates for heritability across previously identified regions of association and to the rest of the genome, when fit jointly (black points and line segments). Red bars indicate estimates after including the dosage of the lead SNP in each of the four association regions as a covariate. c) Residual estimates for heritability partitioned into genic and intergenic regions (left two points), and into minor allele frequency bins. Estimates are made excluding the four association regions in b) and are conditional on the protective dosage at these variants.

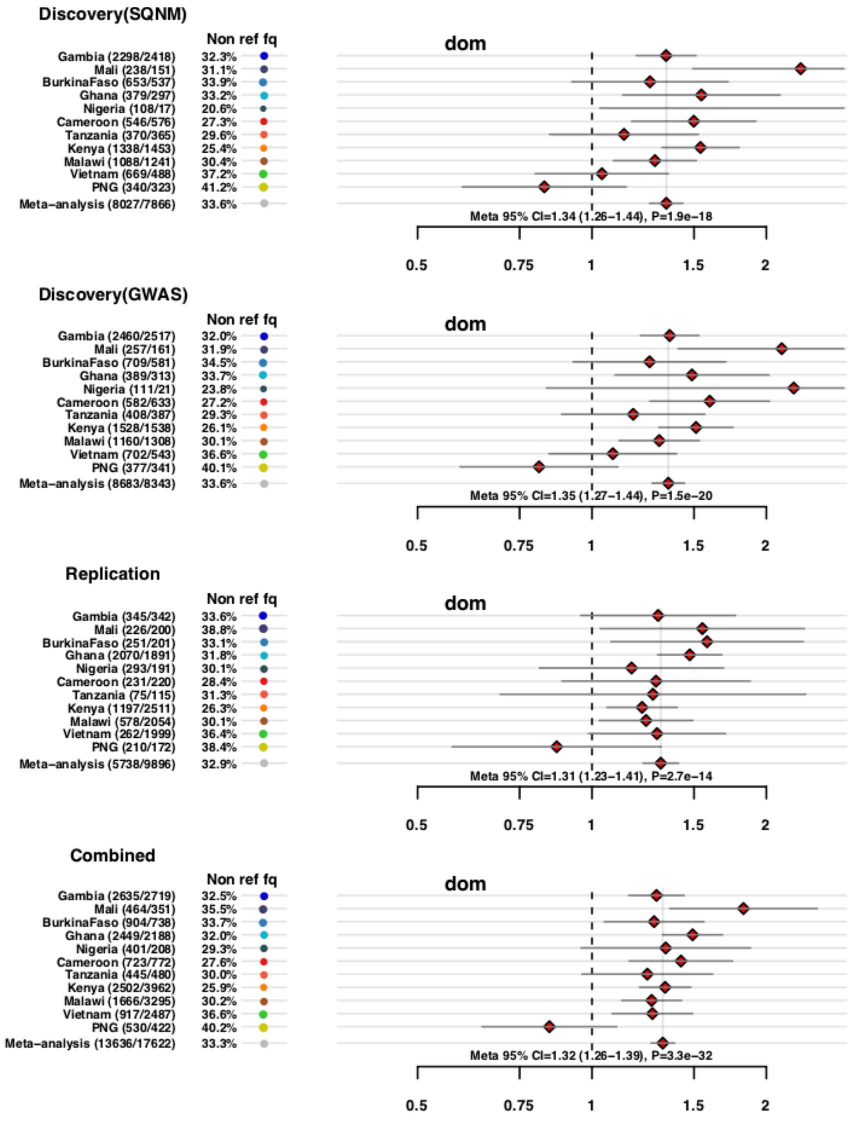

#### Figure S 5 - Discovery and replication effect sizes for the O blood group deletion rs8176719

Estimated odds ratios (points) and 95% confidence intervals (line segments) for rs8176719, under additive, dominant, recessive and heterozygote models of association with severe malaria. The four rows reflect analysis across discovery samples using imputed data; discovery samples using directly-typed genotypes; replucation samples not included in the discovery analysis; and across the combined sample.  Text on left indicates the study population, and case/control counts.  In each row the bottom line reflects fixed-effect meta-analysis across populations.

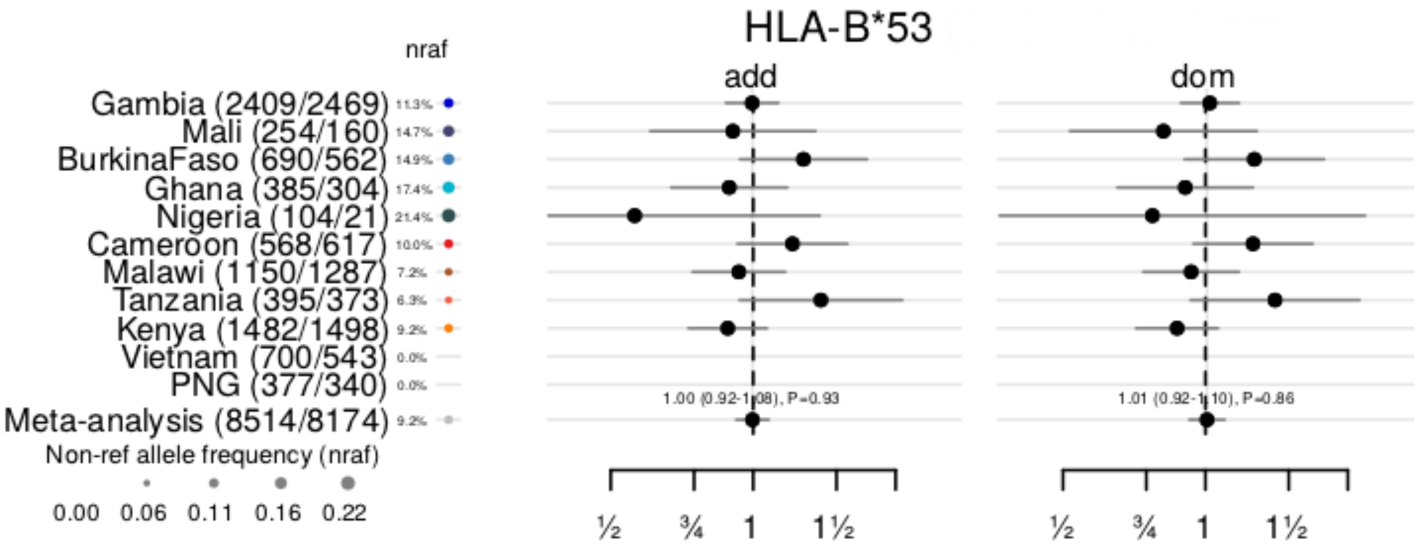

**Figure S 6 - Forest plot showing evidence for association with HLA-B*53**

Plot shows estimated odds ratios (black dots) and 95% confidence intervals (black line segments) for imputed genotypes of HLA-B*53 against severe malaria status. Estimates are presented under additive, dominant, recessive and heterozygote models of association (labels). The bottom row indicates the result of fixed-effect meta-analysis across populations. Coloured circles and small text indicate the allele frequency in control samples, according to the scale on the lower left. The left side of the plot indicates the number of cases and controls included in the analysis in each study population.

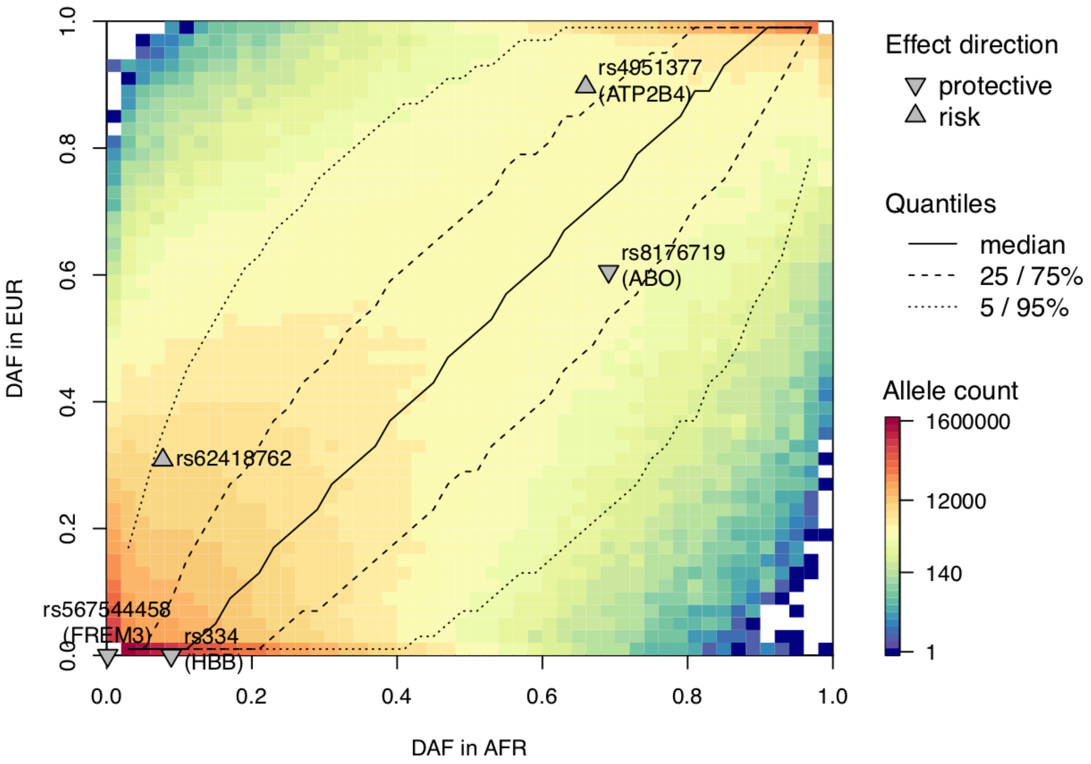

**Figure S 7 - joint derived allele frequency distribution of African and European populations**

Plot shows the empirical distribution of allele counts for derived (i.e. non-ancestral) alleles in African (x axis) and European (y axis) reference panel samples. Counts are aggregated into 1% frequency bins for visualization; colours indicate the number of alleles in each bin according to the scale on the right. Only variants not masked by the 1000 Genomes 'strict' mask, and having an ancestral allele assignment in the 1000 Genomes ancestral allele sequence are included. Black lines indicate the empirical median, 25%, and 5% quantiles of the distribution in Europe conditional on the African allele frequency (i.e. in each vertical 'slice' through the plot). Triangles indicate the position of the five replicating associations on the distribution, with 'up' arrows indicating that the risk allele is derived, and 'down' arrows indicating that the protective allele is derived. For each SNP, $rank_{EUR}$ can be computed as the tail of the vertical slice above (for protective derived alleles) or below (for risk derived alleles) the variant, in the direction of the triangle. We include half of the count of the bin containing the SNP so that both tails sum to one.

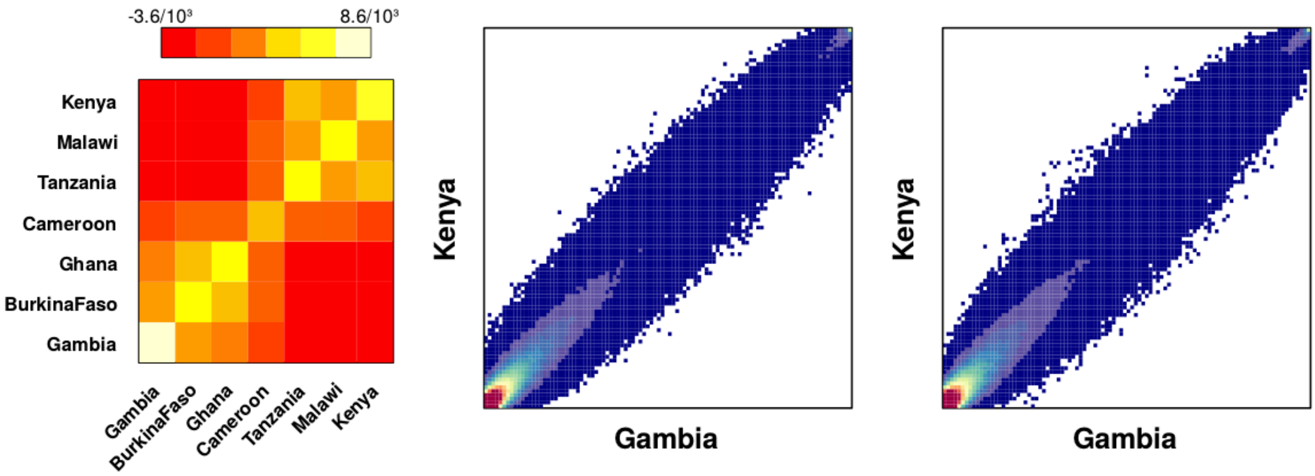

#### Figure S 8 - Empirical model of allele frequencies across African populations

Visualisation of an empirical model of allele frequencies estimated across the seven African study sites with at least 500 samples. Left: covariance matrix of scaled allele frequencies estimated at 100,000 SNPs, randomly chosen from among those have mean frequency in the range 2-98%. Covariance is computed form allele frequencies after subtracting the mean frequency (f) and dividing by the expected standard deviation $\sqrt{f(1-f)}$. b) the empirical joint distribution of allele frequencies in Kenya and the Gambia. c) illustration of the modelled distribution of allele frequencies, for 1,000,000 SNPs simulated from the model in panel a) based on mean frequency sampled from the true mean frequency distribution. Given the root frequency f, we simulated a scaled frequency vector v by sampling based on the estimated covariance matrix. We then plot the frequencies computed as $f+v\sqrt{f(1-f)}$ in Gambia and Kenya.

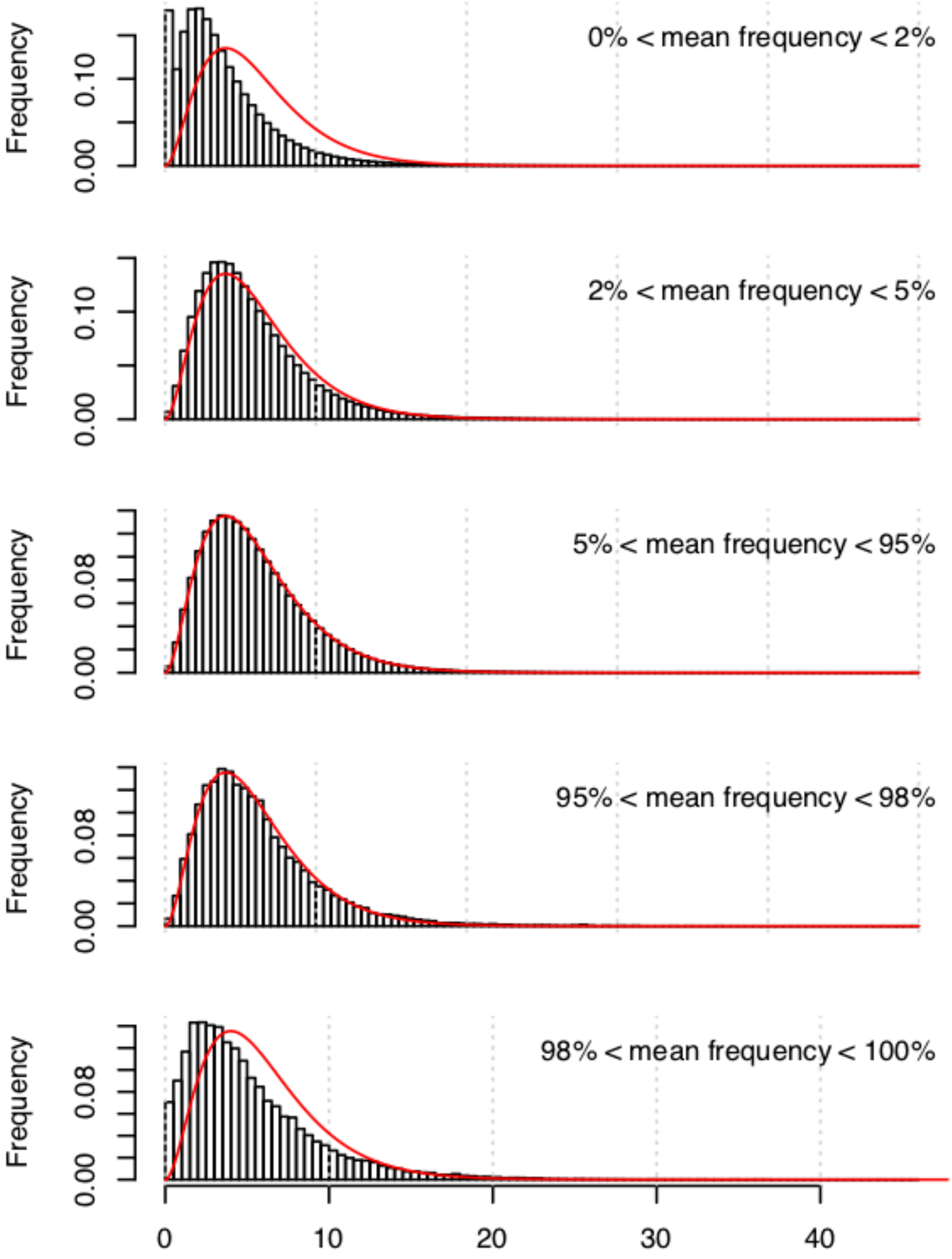

**Figure S 9 - Illustration of the empirical model of allele frequencies across African populations at different frequencies**

Bars show the empirical distribution of the $XX^t$ statistic for all SNPs in the frequency bins indicated by the text. Red lines show the density of the $\chi^2$ distribution used to compute $P_{XXt}$. The test reflects a 6-degree-of-freedom $\chi^2$ distribution; we remove one population (Cameroon) from the test to account for subtracting the mean frequency.. The theoretical distribution matches the true distribution closely, except for rarer variants where the theoretical distribution appears conservative.

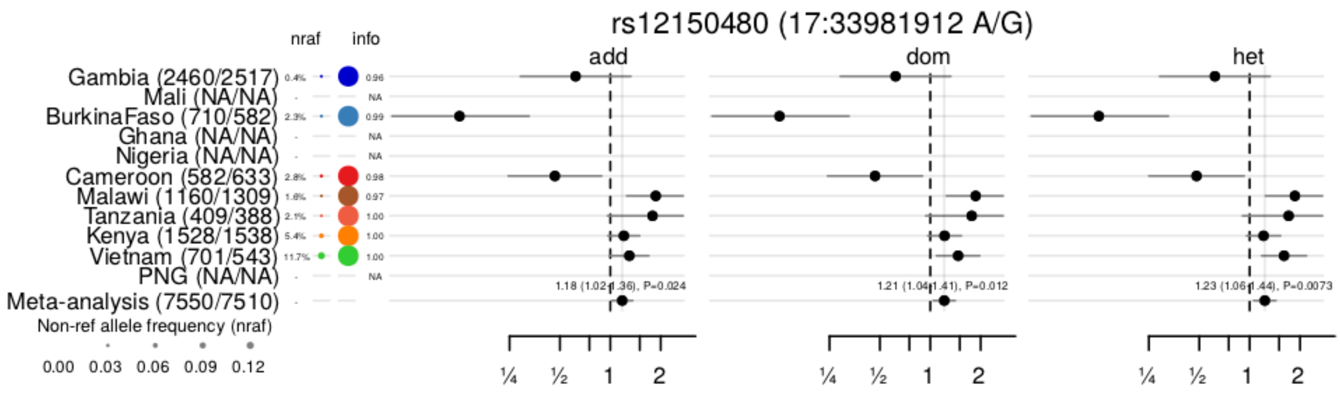

#### Figure S 10 - Estimated effect sizes for rs12150480 (AP2B1)

Plot shows estimated odds ratios, confidence intervals, and meta-analysis results for rs12150480 under additive, dominance and heterozygote models. Blank rows indicate those where an allele count less than 25 was observed. The bottom row indicates the result of fixed-effect meta-analysis across populations. Coloured circles and small text indicate the allele frequency in control samples, according to the scale on the lower left. The left side of the plot indicates the number of cases and controls included in the analysis in each study population.

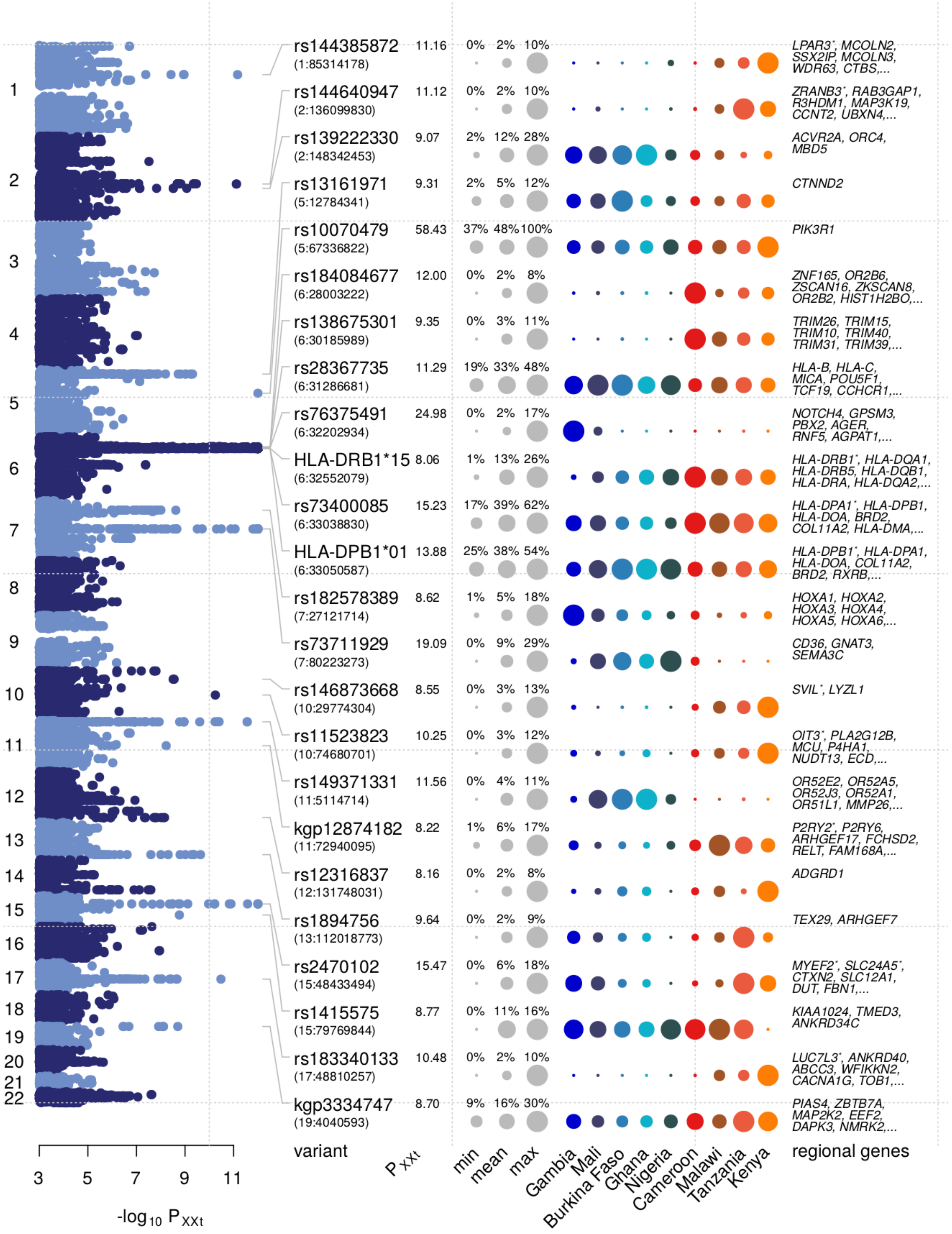

**Figure S 11 - Evidence for within-Africa differentiation at variants across the genome**

Plot shows the evidence for differentiation (-$\log_{10}$ $P_{XXt}$, computed using controls from the 7 African study populations with at least 500 samples) at all variants with at least 2% mean allele frequency across the genome. Variants include genome-wide imputed SNPs, imputed HLA alleles, and glycophorin CNVs. Table on right shows a set of variants selected to have $P_{XXt} < 1E\text{-}8$, and thinned such that no two variants lie within a distance of 0.5cM with a 25kb margin of each other (thinned separately for imputed HLA alleles and genome-wide variants). For each variant we show the ID, chromosome and position and the $-\log_{10} P_{XXt}$. Circles show the allele frequencies, with the area of each circle reflectin the minor allele frequency in proportion to the maximum minor allele frequency across African populations (shown as grey circle). Genes shown are the closest protein-coding genes that appear within the corresponding thinning region; asterisk denotes that the variant lies in the gene, and ellipsis indicates that further genes (not shown) lie within the region.

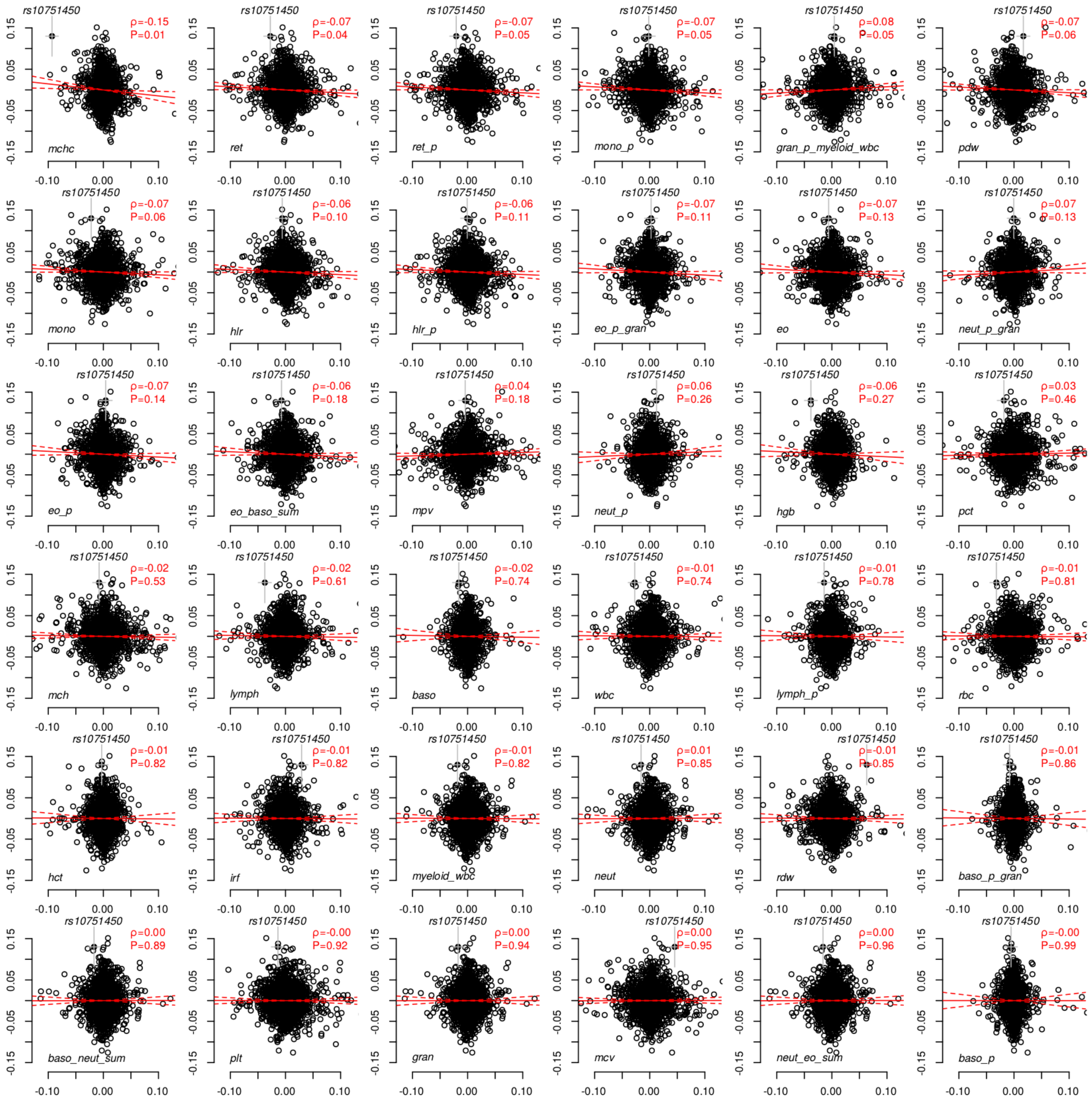

#### Figure S 12 - Mendelian randomization analysis with 36 haemotopoetic traits

Results depict mendelian randomization analysis against each of 36 traits analysed in Astle et al, Cell (2017). Points show the Mendelian randomization analysis of SM and each trait, at 2130 'sentinel' SNPs from Astle et al and having association results in our study. The estimated correlation and P-value are shown in red. Traits are ordered by p-value from lowest to highest. For further details see **Methods** and the legend for **Fig. 4**.

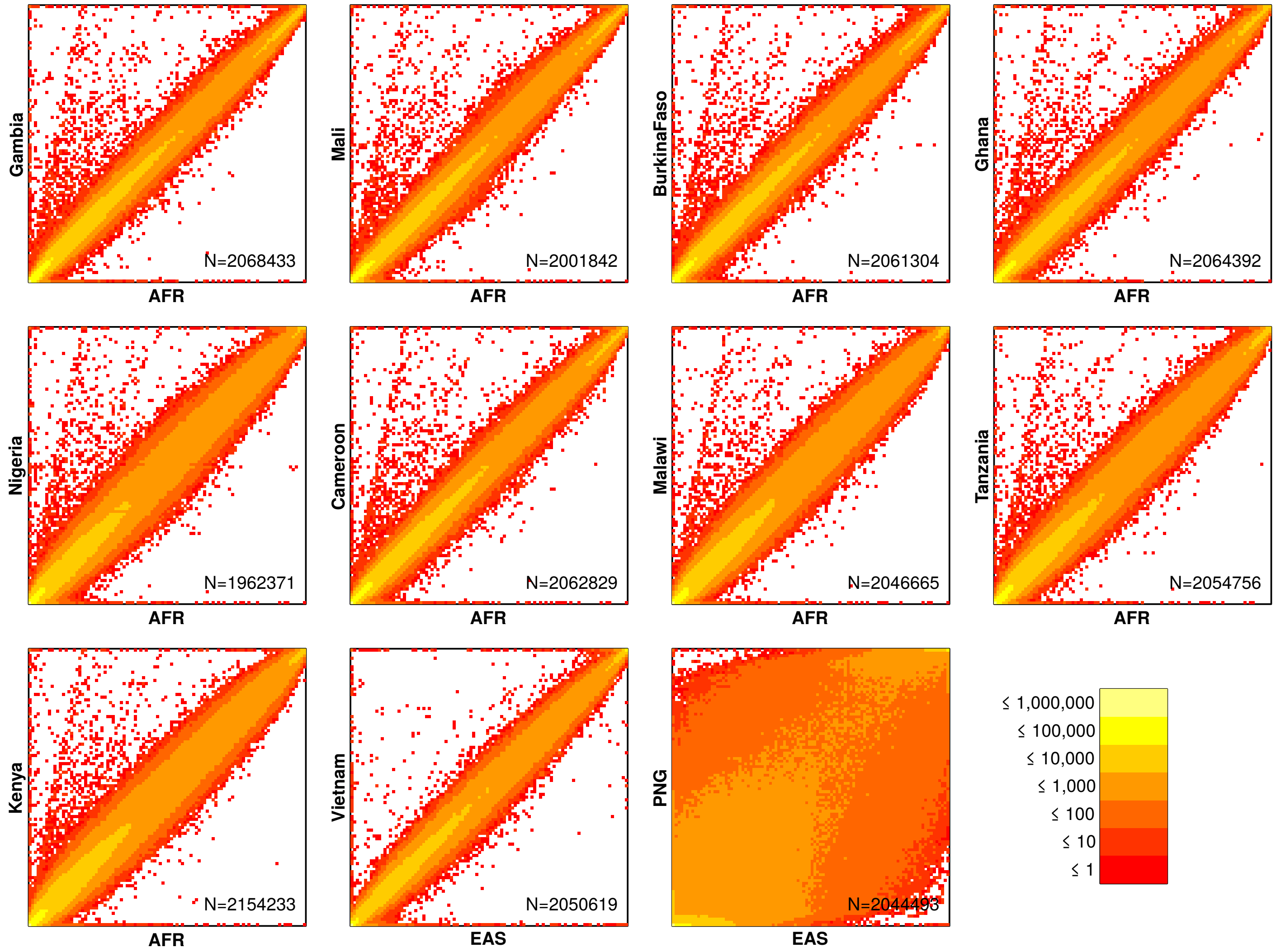

#### Figure S 13 - Comparison of reference panel and study frequencies at typed SNPs

Figure shows the distribution of non-reference allele frequencies in the aligned genotype data for each SNP. The X axis denotes the frequency in the closest reference panel group, which is AFR for African populations and EAS for non-African study populations. Only SNPs with missing proportion < 10% in each study population, and that are also present in the combined reference panel (as identified by genomic position and alleles) are shown. Numbers in text denote the total number of SNPs plotted, and colours denote the counts in each cell according to the legend at the bottom right.

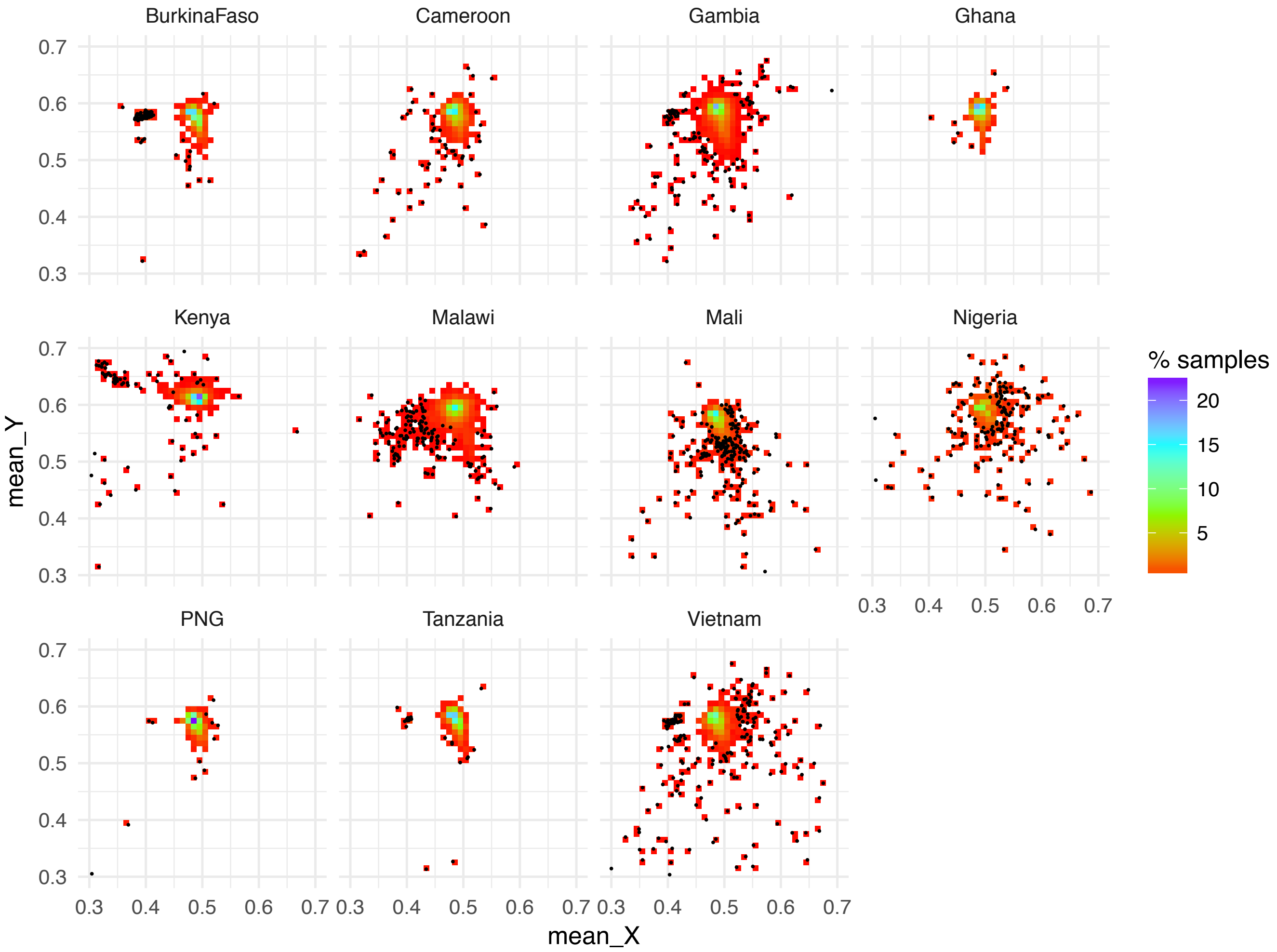

**Figure S 14 - Average X and Y channel intensity values per sample in each population, and intensity exclusions.**

Heatmaps show the proportion of samples in each site with the given average intensity values. Excluded samples were computed using ABERRANT and are shown as black dots.

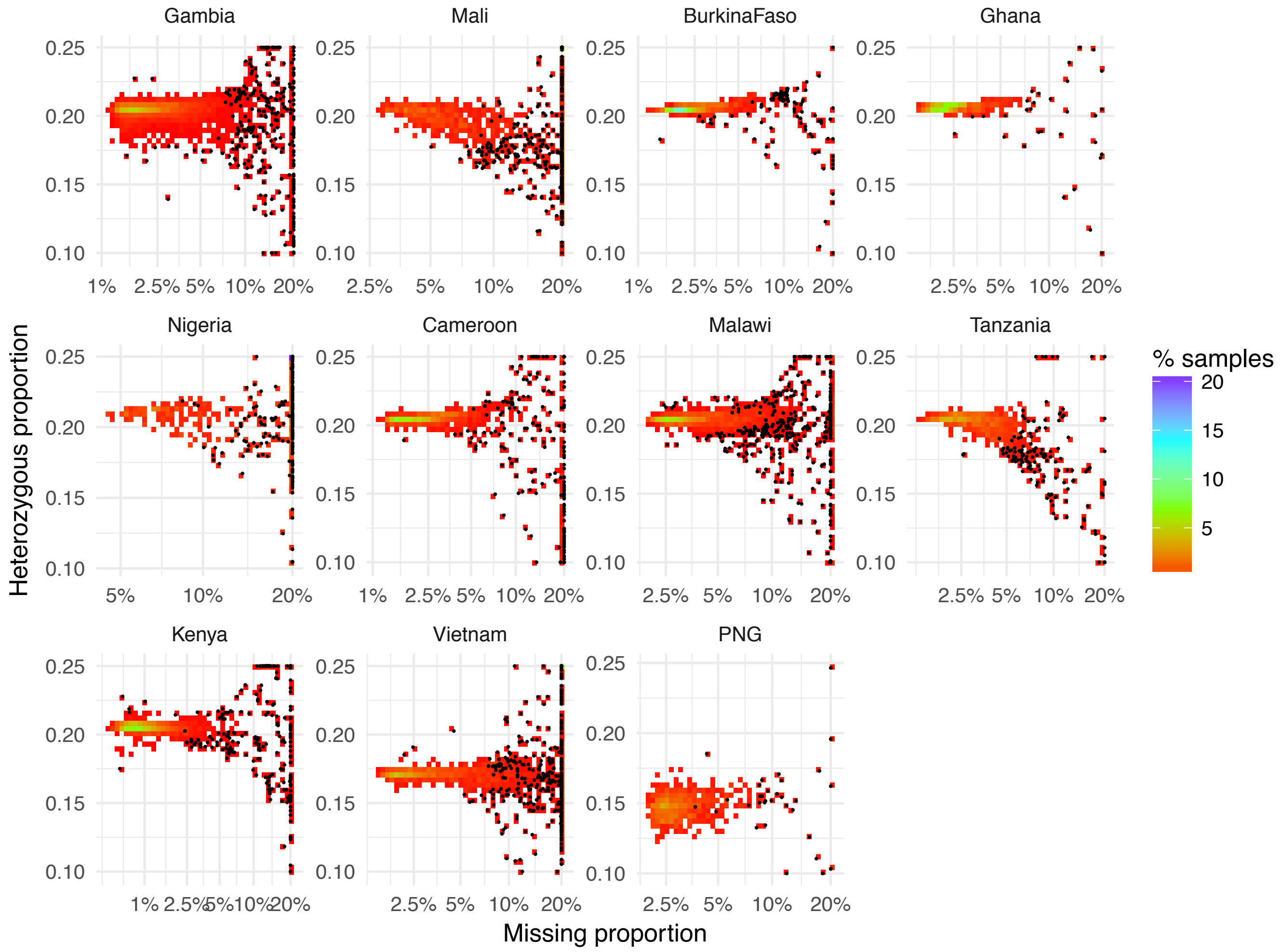

**Figure S 15 - average missing call and heterozygosity rates per sample, and sample exclusions.**

Heatmaps show the proportion of samples in each population with the given average missing call rate (on a logit scale, x axis) and average heterozygosity (y axis). Exclusions were computed using ABERRANT and are shown in black dots.

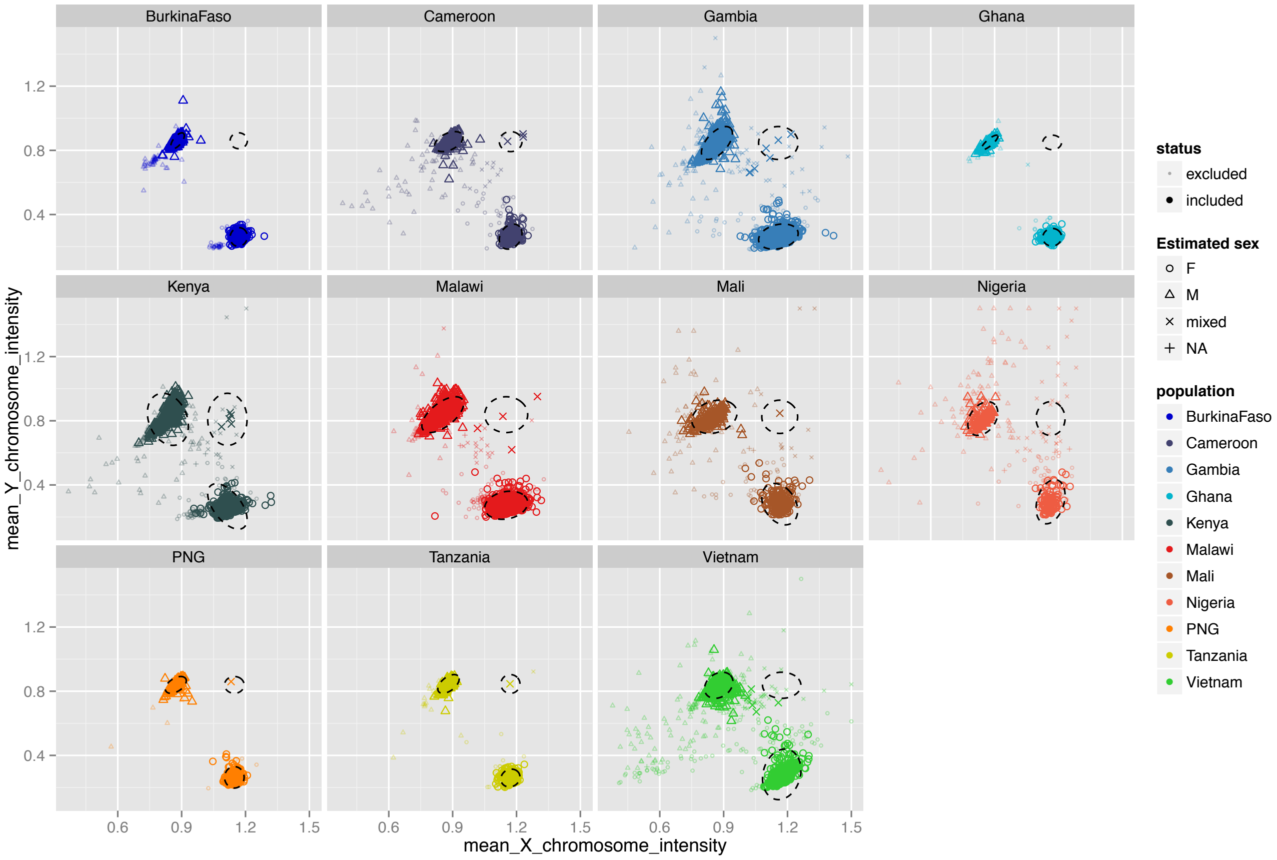

**Figure S 16 - Average X+Y channel intensities across sex chromosomes, and gender assignment.**

Plot shows total (X+Y) channel intensity across X chromosome (x axis) and Y chromosome (y axis) for each sample. Shapes denote the gender assignment of each sample, and are estimated using the cluster positions denoted with dashed ellipses. Excluded samples are shown as smaller, transparent points.

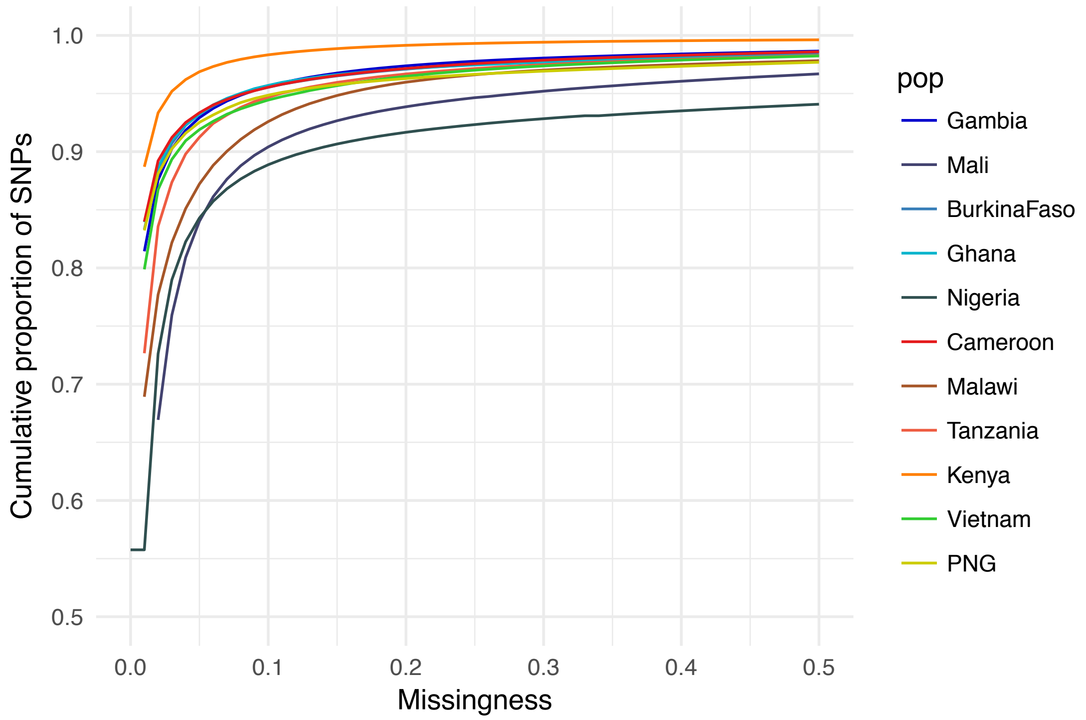

**Figure S 17 - cumulative distribution of per-sample missingness.**

Plot shows the proportion of samples in each study population (y axis) that has less than a given rate of missingness (indicated by the x axis), across all at SNPs on the Omni 2.5M platform after data alignment.

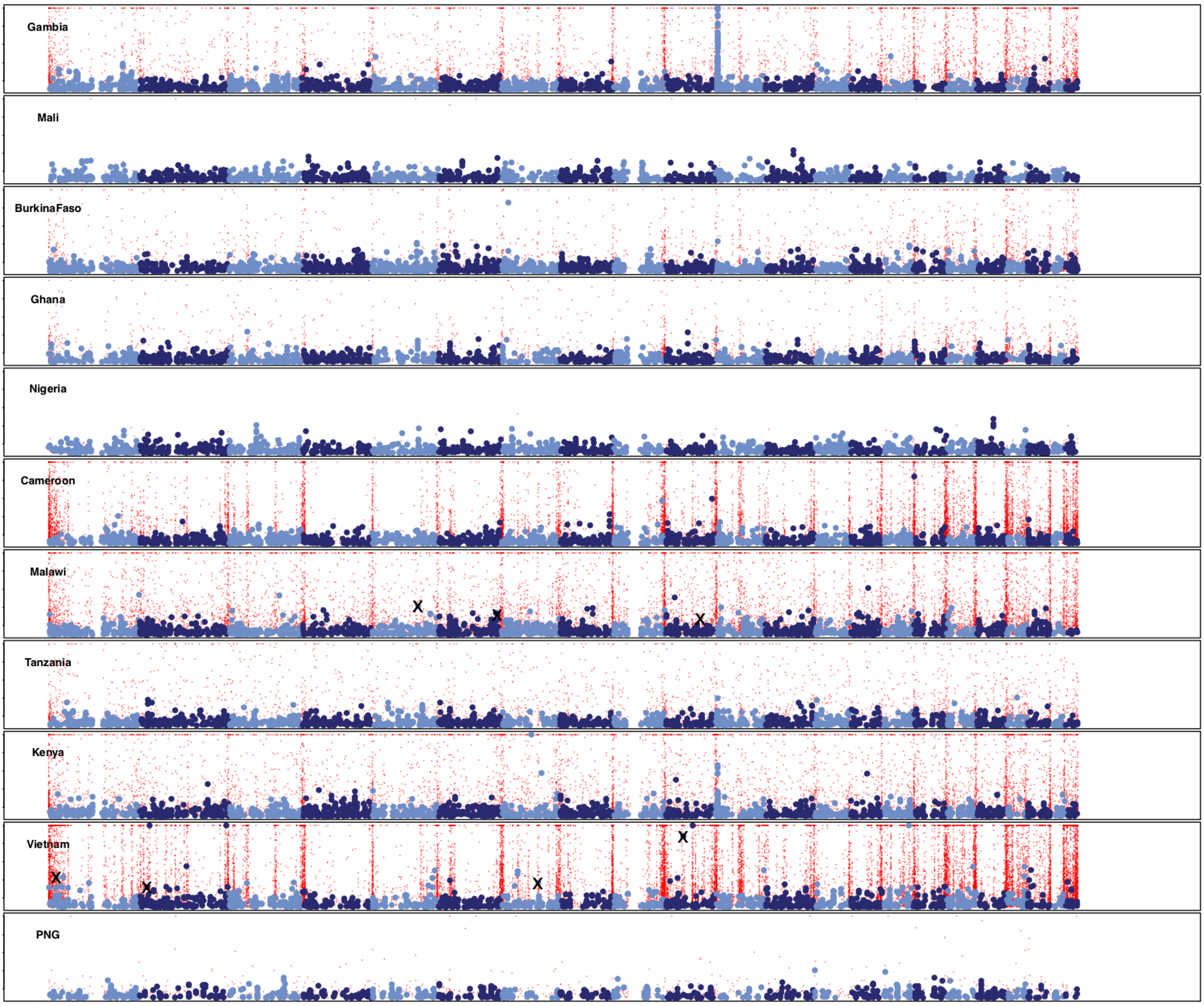

#### Figure S 18 - Detail of QC: manhattan plots.

Rows show manhattan plots for association of severe malaria with directly-typed SNPs, under a general model of association. Red points denote SNPs that were removed during our QC process. Black crosses denote SNPs that were manually removed after inspection of cluster plots.

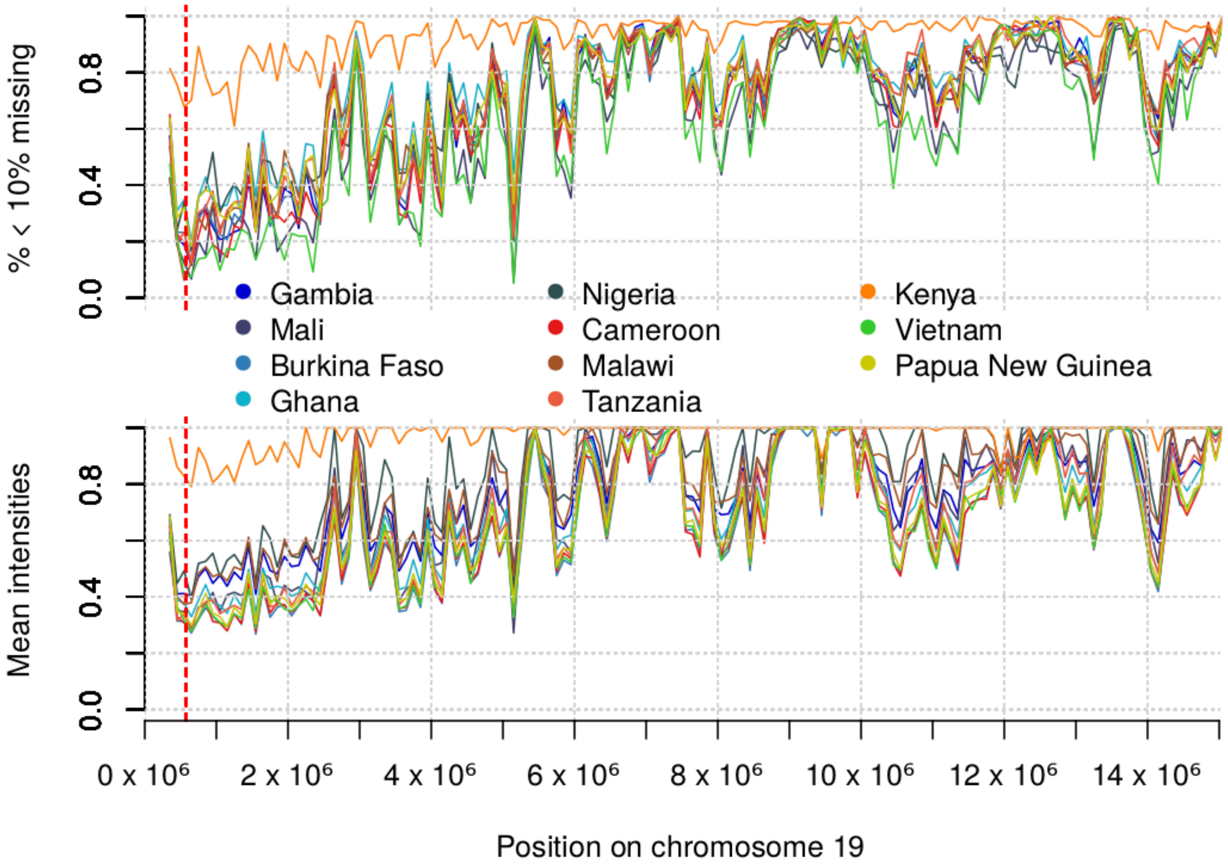

**Figure S 19 Array intensities and genotyping near the end of chromosome 19**

Top panel shows the percentage of variants on the Omni 2.5M microarray which had at least 10% missing calls in our data, for each study population (colours), computed in 100kb bins across the chromosome. Second panel shows the total (X channel+Y channel) intensity for the same SNPs, averaged over 100kb bins. The vertical dashed red line depicts the location of the *BSG* gene.

### Supplementary tables

Note: for supplementary tables not listed here, please see the external file 'Supplementary tables.xlsx'.

| Country | Site | Previous publications (candidate variant typing) | Previous publications (microarray data) | Previous publications (sequencing data) |
|---|---|---|---|---|
| **Gambia** | MRC Laboratories, Banjul | MalariaGEN[1,2], Shah et al[3], Clarke et al [4] | Jallow et al[5]; Band et al[6]; MalariaGEN[1]; Leffler et al[7] | Leffler et al[7] |
| **Mali** | University of Bamako, Bamako | MalariaGEN[1,2], Toure et al[8], Clarke et al [4] | - | |
| **Burkina Faso** | Centre National de Recherche et de Formation sur le Paludisme, Ouagadougou | MalariaGEN[1,2], Clarke et al [4] | - | Leffler et al[7] |
| **Ghana** | Kwame Nkrumah University of Science and Technology | MalariaGEN[1,2], Clarke et al [4] | - | - |
| | Navrongo Memorial Institute for Medical Research with Navrongo Health Research Centre | MalariaGEN[1,2], Clarke et al [4] | - | - |
| **Nigeria** | University of Ibadan | MalariaGEN[1,2], Olaniyan et al[9], Clarke et al [4] | - | - |
| **Cameroon** | University of Buea | MalariaGEN[1,2] Apinjoh et al[10,11], Clarke et al [4] | - | Leffler et al[7] |
| **Malawi** | Blantyre Malaria Project with Malawi-Liverpool–Wellcome Programme | MalariaGEN[1,2], Clarke et al [4] | Band et al[6] MalariaGEN[1]; Leffler et al[7] | - |
| **Tanzania** | Joint Malaria Programme, Kilimanjaro Christian Medical Centre, Moshi | MalariaGEN[1,2]; Manjurano et al[12-14], Clarke et al [4] | Ravenhall et al[15] | Leffler et al[7]; Ravenhall et al[15] |
| **Kenya** | KEMRI–Wellcome Research Programme, Kilifi | MalariaGEN[1,2]; Opi et al[16]; Ndila et al[17], Shah et al [18], Ugoya et al [19], Clarke et al [4] | Band et al[6] MalariaGEN[1]; Leffler et al[7] | Leffler et al[7] |
| **Vietnam** | Oxford University Clinical Research Unit, Ho Chi Minh City | MalariaGEN[1,2], Dunstan et al [20], Clarke et al [4] | - | - |
| **Papua New Guinea** | Papua New Guinea Institute for Medical Research, Madang | MalariaGEN[1,2], Manning et al [21], Clarke et al [4] | - | - |

#### Table S 1 - study sites and previous publications

The first two columns lists study site countries and the address of lead research institutions. The last three columns refer to papers published on data generated by this project, including publications using direct typing of candidate variants, publications on genome-wide microarray data, and publications using short-read sequencing data.

#### Table S 2 - The 95 regions with $BF_{avg}$ > 1,000

This table can be found in the external file "Supplementary tables.xlsx". The table reflects the 95 regions with the greatest evidence for association in our data, as determined by inthinnerator ranking variants by $BF_{avg}$. Columns represent the id, chromosome, position, rsid, alleles of the lead variant; the nearest gene and the distance to the nearest gene; P-values computed by fixed-effect meta-analysis across populations for case/control test and for test with SM subphenotypes; the $BF_{avg}$; the model with the highest posterior weight; the number of directly-typed SNPs having $r^2$ >= 0.75 across populations; the rsid of the directly-typed SNP with the highest $r^2$; the replication and overall BF; a two-sided replication P for the case/control model.

#### Table S 3 - heritability estimates.

This table can be found in the external file "Supplementary tables.xlsx". The table shows heritability estimates made using PCGC {Golan, 2014 #168} and GCTA {Yang, 2011 #198} based on directly-typed genotypes in our QCd set of data. Estimates are made based on 13,030 samples from African study populations chosen to have relatedness < 0.05 within populations. We computed principal components across this set of samples and include an indicator of study site and 10, 20, or 50 PCs as covariates to allow for potential confounding by major axes of population structure. We present results for estimates across the whole genome, joint estimates across all chromosomes, estimates of contributions from chromosomes estimated independently, estimates split into regions of replicable associations and the remainder of the genome, and split by variant frequency, as denoted by the first column. Additional columns indicate the covariates included, the subset of SNPs for which the estimate applies (where 'combined' denotes a sum over all other components in the analysis), the number of SNPs included in the subset, the estimated liability scale heritability from PCGC and GCTA and the corresponding standard errors, and the estimated proportion of heritability per SNP.

#### Table S 4 - functional annotation for variants in 95% credible set of top 95 association regions.

This table can be found in the external file "Supplementary tables.xlsx". The table reflects identified functional annotations of variants in the 95% credible set of the regions **Table S2**, where the credible set is computed assuming a single causal variant is present (i.e. by reweighting $BF_{avg}$ across variants). Only variants with $BF_{avg}$ >100 are included. Columns reflect the id, rsid and $BF_{avg}$ of the lead variant in the region; the id, chromosome, position, rsid, $BF_{avg}$, and best posterior model at the annotated variant; indicators of whether the variant lies in a protein coding gene and/or in an exon of a protein coding gene; the gene name where applicable; the output of Variant Effect Predictor; ENCODE transcription factor binding sites the variant lies in; an indicator of whether the variant lies in a GATA1 or TAL1 motif; inferred chromatin states at the variant in selected cell types, from Roadmap Epigenomics Project data; the mean allele frequency and P-value for the XtX test of population differentiation; genes for which the variant has evidence of eQTL in peripheral blood {Westra, 2013 #135}, GTEx tissues {GTEx Consortium, 2015 #110}, or erythrocyte precursors {Lessard, 2017 #121}; RBC trait associations {Astle, 2016 #142}; and GWAS trait associations {MacArthur, 2017 #136}.

#### Table S 5 - Summary of HLA typing

This table can be found in the external file "Supplementary tables.xlsx". Table presents a comparison of HLA classical allele genotypes determined by HLA typing and by imputation, in 31 Gambian children who are cases in our study. Columns represent the HLA locus and allele; counts of samples typed with each genotype for the allele (which we consider as 'true' genotypes); counts of samples imputed with each genotype for the allele (using a cutoff of 0.75 probability where imputation is uncertain); counts of samples of each true genotype wrongly imputed; counts of samples of each imputed genotype wrongly imputed; and the correlation, recall and precision of the imputed genotypes.

#### Table S 6 - variants showing heterogeneous patterns of association

This table can be found in the external file "Supplementary tables.xlsx". The table shows a list of variants having heterogenous patterns of estimated effects across populations, identified as having maximum *BF* ($BF_{max}$) > 25,000 and at least 100 times greater than the *BF* under a fixed-effect model. Only the additive model is considered here. The maximum is computed across all models tested, which include those in **Table S9** and additional population and group-specific models, including individual population-specific effects. We restrict to variants with an effective minor allele count of at least 1,720 (corresponding to a minor allele frequency of 5% across all study samples for well-imputed SNPs). Variants other than the lead variant in the *HBB* and glycophorin regions are removed and we thin results using inthinnerator to identify a single variant within each region. Columns represent the identifier, chromosome, position, rsid, additive component of the $BF_{avg}$, best posterior model, $BF_{max}$, the model attaining the maximum *BF*, the *BF* under structured, correlated and fixed-effects; the containing gene where relevant; the effective minor allele count; the XtX mean allele frequency and P-value; Variant Effect Predictor functional results; eQTL results in peripheral blood {Westra, 2013 #135}, GTEx tissues {GTEx Consortium, 2015 #110}, and erythrocyte precursors {Lessard, 2017 #121}.

| Collection | | Pre-phasing | | | | | Post-phasing | |
|---|---|---|---|---|---|---|---|---|
| Population | Total samples analysed (repeats) | Intensities | Miss / het | Gender | Duplicates / repeats | ***TOTAL*** | Relatedness (PCs, non-case-control sample) | ***TOTAL*** |
| Gambia | 5594 | 137 | 182 | 4 | 99 | *5171* | 192 (1) | *4979* |
| Mali | 900 | 220 | 191 | | 2 | *484* | 52 (3, 10) | *422* |
| Burkina Faso | 1446 | 69 | 28 | | 20 | *1325* | 31 (4) | *1294* |
| Ghana | 782 | 7 | 25 | | 14 | *732* | 11 (3, 15) | *706* |
| Nigeria | 419 | 192 | 84 | | 8 | *133* | 0 (2) | *133* |
| Cameroon | 1471 | 64 | 118 | 3 | 9 | *1272* | 55 (5) | *1217* |
| Malawi | 2791 (297) | 150 | 250 | 3 | 186 | *2495* | 24 (4) | *2471* |
| Tanzania | 979 | 22 | 126 | 1 | 11 | *814* | 16 (5) | *798* |
| Kenya | 3769 (96) | 120 | 116 | 6 | 156 | *3462* | 355 (5, 39) | *3068* |
| Vietnam | 1690 (38) | 263 | 174 | 1 | 26 | *1260* | 13 (4) | *1247* |
| PNG | 815 | 12 | 21 | 1 | 5 | *772* | 51 (4) | *721* |
| **TOTAL** | *20,656 (431)* | 1,257 | 1,315 | 19 | 536 | *17,960* | 800 (40, 64) | *17,056* |

#### Table S 7 - Detail of Sample QC

'Collection' columns show: the population label; total samples analysed per study population (all samples, including repeat samples were included in the QC process). 'Pre-phasing' columns show: samples removed due to outlying intensities, outlying missingness/heterozygosity, unassigned gender, genetic due to being a genetic duplicate, and the total number of samples included in phasing.  'Post-phasing' columns show: the number of samples excluded from association testing due to being closely related to other samples, identified as outlying on principal components, or lacking case/control status (e.g. for parent samples); and the total included in association testing.

| Population | Total SNPs | Missingness | frequency | HWE / MF | plate | Recall | Cluster / platform | TOTAL |
|---|---|---|---|---|---|---|---|---|
| *Autosomes* | | | | | | | | |
| **Gambia** | 2,312,228 | 163,463 | 457,765 | 2,129 | 5,082 | 646 | | |
| **Burkina Faso** | 2,312,228 | 156,296 | 464,801 | 75 | 3,242 | 34 | | |
| **Ghana** | 2,312,228 | 156,576 | 446,825 | 14 | 2,720 | | | |
| **Cameroon** | 2,312,228 | 153,740 | 451,465 | 69 | 2,824 | 22 | | |
| **Malawi** | 2,312,228 | 171,305 | 461,725 | 1,022 | 12,771 | 2361 | | |
| **Tanzania** | 2,312,228 | 123,024 | 458,999 | 45 | 4,003 | 155 | | |
| **Kenya** | 2,374,031 | 132,070 | 480,658 | 382 | 3,296 | 20 | | |
| **Vietnam** | 2,312,228 | 187,266 | 927,210 | 30 | 1,953 | 17,126 | | |
| **Papua New Guinea** | 2,312,228 | 246,303 | 1,017,977 | 19 | 1,929 | | | |
| *Combined* | 2,383,648 | 426,545 | 289,371 | 1,315 | 28,026 | 36,823 | 6/51,048 | 1,550,514 |

| | | | | | | | | |
|---|---|---|---|---|---|---|---|---|
| *X chromosome* | | | | | | | | |
| **Gambia** | 55,510 | 5,138 | 8093 | 753 | 249 | | | |
| **Burkina Faso** | 55,510 | 4,154 | 8,313 | 441 | 234 | | | |
| **Ghana** | 55,510 | 4,301 | 8,120 | 221 | 439 | | | |
| **Cameroon** | 55,510 | 4,191 | 8,547 | 587 | 135 | | | |
| **Malawi** | 55,510 | 6,174 | 8,419 | 830 | 216 | | | |
| **Tanzania** | 55,510 | 3,746 | 8,186 | 185 | 354 | | | |
| **Kenya** | 57,044 | 6,351 | 8,563 | 569 | 179 | | | |
| **Vietnam** | 55,510 | 6,185 | 19,489 | 278 | 247 | | | |
| **Papua New Guinea** | 55,510 | 6,851 | 21,632 | 251 | 219 | | | |
| *Combined* | 57,104 | 13,740 | 4,719 | 224 | 2,331 | - | 2,695 | 33,395 |

#### Table S 8 - Detail of SNP QC

Columns show: the population label, total number of SNPs genotyped, SNPs removed due to missingness in each population, SNPs additionally identified as low frequency, out of Hardy-Weinberg equilibrium, or failing the plate or recall test in each population. The 'Combined' row shows the total number of SNPs and the number failing each combined filter, including the criteria of at least two populations with frequency > 1%. For the X chromosome, missingness and plate test were computed separately in males and females, and a test of difference in frequency between males and females was used in place of HWE.

| **Category / model name** | **Short name** | **Detail** | **Prior weight in $BF_{avg}$** |
|---|---|---|---|
| *Mode of inheritance* | | | |
| Additive | add | Encoded as AB+2*BB | 0.4 |
| Dominant | dom | Encoded as AB+BB | 0.2 |
| Recessive | rec | Encoded as BB | 0.2 |
| Heterozygote | het | Encoded as AB | 0.2 |
| | | ***Total*** | 1 |
| *Population* | | | |
| Fixed effects | fix | All off-diagonal entries of $P$ are set to 0.99 | 0.4 |
| Correlated effects | cor | All off-diagonal entries of $P$ are set to 0.9 | 0.2 |
| Independent effects | Ind | All off-diagonal entries of $P$ are set to 0 | 0.04 |
| Structured effects | str | Uses $P$ estimated from the correlation in allele frequencies genome-wide | 0.04 |
| Population group-specific effects | | Effects assumed to be restricted to a subset of populations. | 0.04 per subset; 8 subsets in total. |
| | | ***Total*** | 1 |
| *Subphenotype effects* | | | |
| Case/control effects | | (models as described above) | 0.8 |
| Correlated between subphenotypes | cm-sma-other-cor | Between-phenotype entries of $P$ set to 0.9 | 0.04 |
| Independent across subphenotypes | cm-sma-other-ind | Between-phenotype entries of $P$ set to 0 | 0.04 |
| Effects restricted to two subphenotypes | | $\sigma$ nonzero for two phenotypes | 0.02 per pair of subtypes |
| Effects restricted to one phenotype | | $\sigma$ nonzero for one phenotype | 0.02 per subtype |
| | | *Total* | 1 |

##### Table S 9 - Detail of models included in $BF_{avg}$

The table shows the component models included in the model-averaged Bayes factor. Columns specify the model name and shortened mnemonic name, detail of the implementation, and the prior weight in the $BF_{avg}$. Prior weights are given per category; to compute the full prior weight for a model, weights should be multiplied across categories. For example, the model of dominant effects on cm and sma is weighted as 0.2*0.02 = 0.004. For subphenotype analysis, between-population correlation for each phenotype was set to 0.99, and between-population between-phenotype correlation was set to 0.99 times the assumed between-phenotype correlation.

### Supplementary text

#### Text S1 - Investigation of the *HBA1-HBA2* region

Variation in the genes encoding alpha globin (*HBA1*, chr16:226679-227520; and *HBA2*, chr16:222846-223709) have been linked to malaria susceptibility as reviewed previously [22,23]. In particular, the $-\alpha^{3.7}$ deletion, which deletes a 3.7kb sequence forming a hybrid *HBA2-HBA1* gene, is a cause of alpha thalassaemia and is found at nontrivial frequency in African populations. Alpha thalassaemia is thought to be protective against malaria, but direct evidence for this hypothesis is currently somewhat limited. Available evidence comes both from observation of the distribution across populations (reviewed in [24]) and from direct testing of this variant in case/control samples (e.g. OR=0.83 (0.76–0.90); $P=2\times10^{-6}$; observed using direct typing of N=6193 children in Kilifi, Kenya [17]; these samples are also included in our study).

We did not observe strong signal of association across the *HBA1-HBA2* region in our GWAS study (e.g. maximum $BF_{avg}$ = 67 within the first megabase of chromosome 16; at rs186272033, chr16:83520; maximum $BF_{avg}$ = 12 within 100kb of *HBA1-HBA2*). The 1000 Genomes reference panel contains a variant identified from sequence reads that corresponds to the $-\alpha^{3.7}$ deletion (appearing in the panel as EM_DL_DEL34404, chr16:223678±150-227490±150, length = 3812; combined frequency = 5.5% across African ancestry samples in the 1000 Genomes project). In our data, EM_DL_DEL34404 was imputed at high frequency in all African populations (allele frequency = 8.2%, 10.6%, 13.4%, 16.2%, 24.9%, 19.7%, 29.4%, 26.4%, 30.2% in Gambia, Mali, BurkinaFaso, Ghana, Nigeria, Cameroon, Malawi, Tanzania and Kenya respectively; frequency = 1.4% in Vietnam and <1% Papua New Guinea). Imputation confidence was also nontrivial (IMPUTE info = 0.73-0.86 in African study populations). However, we observed only modest evidence for association with EM_DL_DEL34404 ($BF_{avg}$ = 10.3; fixed-effect additive $P_{add}$ = 0.002; OR = 0.90; 95%CI = 0.84-0.96). The strongest evidence for association in individual populations was observed in Malawi (OR=0.83; 95%CI = 0.73-0.95; $P_{add}$=0.007).

These findings may be taken to confirm that $-\alpha^{3.7}$ confers a modest protective effect. However, we also noted reasons that suggest these results, based on imputation, should be treated with caution. First, the *HBA1-HBA2* region lies near the start of chromosome 16, and is susceptible to an observed end-of-chromosome effect on study genotype quality (described further in Text S2 below). This limits the amount of data informing on imputation, such that only 115 SNPs in our post-QC set lie in the region chr16:0-1,000,000.

Second, LD between EM_DL_DEL34404 and regional SNPs appears relatively weak and differs substantially between populations (max $r^2$ = 0.53 between EM_DL_DEL34404 and all other reference panel variants in the first megabase of chromosome 16; this maximum is attained at rs76462751 in the ESN and YRI populations; $r^2$ = 0.001 at the same SNP in GWD and < 0.25 in MSL and LWK; $r^2$ < 0.4 for all other SNPs except in ESN; no SNPs with $r^2$ > 0.1 in all African populations). This pattern of LD suggests that the $-\alpha^{3.7}$ deletion may be carried on several distinct haplotypes across populations, a situation which is naturally challenging for imputation-based approaches. (In particular, this appears incompatible with a hypothesis of a single recent origin and subsequent positive selection of $-\alpha^{3.7}$, but does appear consistent with the high observed de novo mutation rate of this deletion [25]).

Finally, comparison of imputed $-\alpha^{3.7}$ genotypes to previously reported direct-typing in Kenya [17] reveals a relatively low accuracy that is overestimated by the IMPUTE info score ($r^2$=0.42 between expected number of copies of $-\alpha^{3.7}$ inferred from imputation and copies inferred from direct typing, in N=2913 Kenyans; IMPUTE info = 0.86). These results do not take account of other, globally rarer thalassaemia-causing alleles. Greater accuracy of inference in this region will be of interest.

#### Text S2 - Investigation of the basigin region

Basigin has been identified as a receptor for *P.falciparum* malaria during invasion of red blood cells, and the interaction between basigin and the Pf ligand PfRh5 is thought to be essential to the invasion process[26]. It is therefore plausible that the gene encoding basigin (*BSG*, chr19:572454-583493) might harbour genetic mutations that affect invasion and hence malaria infection outcomes. However, we found little signal of association across the region containing *BSG* (e.g. maximum $BF_{avg}$ = 29 at rs10425584, approximately 150kb upstream of *BSG*). On closer inspection we noted that few typed SNPs in the region are contained in our set of QCd haplotypes (e.g. 67 typed SNPs after QC in the first megabase of chromosome 19, compared to approximately 420 per Mb across the whole of chromosome 19). We plotted SNP QC metrics in the first 15Mb of chromosome 19 (**Figure S19**) and noted generally poor genotyping across the region. Specifically, we noticed lower-than average normalised array intensities and low rates of genotype calling across the first ~5Mb of chromosome 19. Similar, but less extreme issues were seen on other chromosomes and we suggest that this likely reflects issues with preparation of samples via whole genome amplification. The Kenyan study population, which was typed on the Omni 2.5M 'quad' platform, was affected by this issue but in a less extreme way than other populations.

#### Text S3 - Investigation of association in *G6PD* and *CD40LG*

We have previously reported evidence of association within *G6PD*[2,4] and upstream of *CD40LG*[2], both of which lie on the X chromosome, using direct typing of variants which are common in African populations. Here we compare these results to those based on our imputed data. Both rs1050828 (*G6PD c.*202C>T, chrX:153,764,217) and rs3092945 (chrX:135,729,609, upstream of *CD40LG*) were imputed with high confidence (info > 0.9) in all African populations. However, under our bayesian model average, evidence for association at both variants was weak ($BF_{avg} < 1$) and there was little evidence for association at variants across these regions ($BF_{avg} < 10$ within 100kb of rs1050828 or rs3092945), except for $BF_{avg}$ = 36 at rs369388464, which lies in an intron of *ARHGEF6*).

We note two explanations for these results. First, the observed effects at rs1050828 in *G6PD* have been noted to be complex, with putative opposing effects in males and females (which would not be picked up by our analysis, which treats males like homozygous females for the purpose of association testing) and in severe malaria subtypes. We did note weak evidence for an SMA-only effect of rs1050828 (*BF* = 6 for SMA-only model; $OR_{SMA}$ = 1.3, 95% CI 1.07-1.71, $P$ = 0.01; $OR_{CM}$ = 0.95). These results are thus consistent with previous estimates based on direct typing in these samples [2,4], though we did not observe evidence for a protective effect on CM in this analysis.

Second, we noted some evidence at both SNPs of discrepancies between direct typing and imputation in specific populations. Specifically, at rs1050828, correlation between imputed and directly-typed genotypes was > 0.9 in all African study populations except The Gambia ($r^2$ = 0.73). At rs3092945 we also found discrepancies in The Gambia ($r^2$ = 0.73) and Kenya ($r^2$ = 0.8). This is particularly notable because the reported signal of association is driven by strong and opposing observed effects in The Gambia and in Kenya, with little evidence in other populations. Inspection of directly-typed data suggests that the directly-typed rs3092945 is out of Hardy-Weinberg equilibrium in the Gambia and other populations. Our tentative interpretation is that the imputation is likely accurate, and the observed association may be at least in part driven by typing artifacts.

#### Text S4 - Analysis of functional annotations

We reasoned that functional annotations might provide clues to further true associations among our list of most associated regions, as well as to the likely causal variants within these regions. To assess this, we examined functional annotations of all variants (described in Methods) with modest or strong evidence of association (defined as having $BF_{avg}$ >= 100 and lying in the 95% credible set of a lead variant with $BF_{avg}$ >= 1000, under the assumption of a single variant within the association region[35]). These results are listed in **Table S4**.

Four protein-altering mutations lie in this list: the mutations encoding HbS and O blood group, which are the sole members of their 95% credible sets, and SNPs in the aminoacyl-tRNA synthetase *IARS* (rs2070053, $BF_{avg}$=1.2x$10^3$) and in *ANKRD30B (*rs73427575, $BF_{avg}$=385). The derived allele at rs2070053 is at only 1-2% frequency and is predicted to have a strong risk effect in our data ($OR_{add}$=1.65, 95% CI=1.32-2.05), but its effect on the protein is reported to be benign [27]. rs73427575 may be deleterious, but has a complex observed pattern of effects across populations. We note also that the copy number variant DUP4, which we have found to underly the association in the glycophorin region, affects protein coding sequence through structural rearrangement of the underlying region.

A number of regions contain SNPs with evidence of potentially regulatory effects. We note here variants with multiple lines of evidence - namely those with evidence of eQTL effects, which also lie in an annotated transcription factor binding site, and which have previously been associated to other traits. This list includes the associations in *ATP2B4,* and at rs2523650 in the human leukocyte antigen region (HLA) on chromosome 6, which are described in main text. Also on this list is a variant in the first exon of the gene *H6PD*, which encodes hexose-6-phosphate dehydrogenase (rs12144045, $BF_{avg}$=1.2x$10^3$*)* and is evolutionarily and functionally related to *G6PD*. Finally, an associated eQTL for *VAC14* (rs8060947, $BF_{avg}$= 550), which encodes a component of the PIKFYVE complex, has recently been associated with *S.Typhi* invasion in vitro [36], with susceptibility to typhoid [37] and with some forms of bacteraemia [38], putatively through altering expression of *VAC14* with downstream effects on cholesterol.

The combined evidence of function and association with susceptibility to malaria at these loci may be considered compelling. However, complicating the results above is that neither the main signal in the HLA region nor that at *VAC14* appear to replicate in our additional replication samples (**Table S2).** We did not attempt to replicate the association at *H6PD*.

### Supplementary Text 5 - Bayesian analysis of replication

##### Replication analysis for a single association model

Consider assessing the evidence for association under a single model of association (denoted $M_1$, parameterised by a vector of parameters $\theta$) versus the model of no association (denoted $M_0$, corresponding to $\theta \equiv \theta_0$), and suppose we have two tranches of data - a discovery set $D_1$ and a replication set $D_2$. We assume these are sampled independently i.e. are conditionally independent given the true parameter value. The overall evidence in the data for $M_1$, relative to $M_0$, is expressed in the Bayes factor

$$BF^{\text{overall}} = \frac{P(D_1, D_2|M_1)}{P(D_1, D_2|M_0)} = \frac{\int_\theta P(D_1|\theta)P(D_2|\theta)P(\theta|M_1)}{P(D_1|M_0) \cdot P(D_2|M_0)} \tag{1}$$

where $\theta$ represents the parameters of $M_1$ (i.e. the genetic effect sizes). Since the Bayes factor based only on discovery data is

$$BF^{\text{discovery}} = \frac{\int_\theta P(D_1|\theta)P(\theta|M_1)}{P(D_1|M_0)}$$

multiplying and dividing by $BF^{\text{discovery}}$ gives

$$BF^{\text{overall}} = BF^{\text{discovery}} \cdot \int_\theta \left( \frac{P(D_2|\theta)}{P(D_2|M_0)} \cdot d_1(\theta) \right) \tag{2}$$

where $d_1(\theta)$ is the posterior mass on parameter value $\theta$ given the discovery data,

$$d_1(\theta) = P(\theta|D_1, M_1) = \frac{P(D_1|\theta)P(\theta|M_1)}{\int_{\theta'} P(D_1|\theta')P(\theta'|M_1)}$$

The second term in equation (2) can be interpreted as the evidence for $M_1$ versus $M_0$ in the replication data, given the effect size distribution learnt from the discovery data. We denote this quantity by $BF^{\text{replication}}$ so that

$$BF^{\text{overall}} = BF^{\text{discovery}} \cdot BF^{\text{replication}} \tag{3}$$

Formula(3) is a basic reflection of the consistency of bayesian reasoning, in the sense that inference is unaffected by whether all data is treated together (as in the left hand side), or in tranches (as in the right hand side). This is an intuitively obvious property but we note that no similar property holds for commonly-used frequentist assessments of replication, since there is no simple relationship between the P-value computed across all data and those computed in discovery and replication samples seperately. (We note that $BF^{\text{replication}}$ is not the same as the Bayes factor that would be computed in replication data using the original prior effect size distribution, i.e. ignoring the discovery data.)

##### Replication analysis using model averaging

Now suppose $M_1, \cdots, M_K$ are $K$ models of association with prior weights

$$w_i = P(M_i|\text{variant is associated}) \qquad \sum_i w_i = 1$$

Then the overall evidence for association can be assessed by summing over models,

$$BF^{\text{overall}}_{\text{avg}} = \sum_i w_i \cdot BF^{\text{discovery}}_i \cdot BF^{\text{replication}}_i$$

Again, multiplying and dividing by the discovery model-averaged Bayes factor $BF^{\text{discovery}}_{\text{avg}}$ gives

$$BF^{\text{overall}}_{\text{avg}} = BF^{\text{discovery}}_{\text{avg}} \cdot \sum_i \left( \frac{w_i \cdot BF^{\text{discovery}}_i}{\sum_j w_j BF^{\text{discovery}}_j} \right) \cdot BF^{\text{replication}}_i$$

The term in the bracket is the posterior weight on model $M_i$ given the discovery data, conditional on one of the models of association being true. We write $w'_i$ for this posterior weight and note it is simply computed by renormalising discovery data Bayes factors. With this convention, the discovery and replication evidence can be summarised in three quantities:

1. A model-averaged discovery Bayes factor based on a chosen set of prior weights,

$$BF_{\text{avg}}^{\text{discovery}} = \sum_i w_i BF_i^{\text{discovery}}$$

2. A model-averaged replication Bayes factor based on posterior weights and effect size distributions learnt from discovery data,

$$BF_{\text{avg}}^{\text{replication}} = \sum_i w_i' BF_i^{\text{replication}}$$

3. An overall model-averaged Bayes factor, which decomposes as a product of the two terms above

$$BF_{\text{avg}}^{\text{overall}} = BF_{\text{avg}}^{\text{discovery}} \cdot BF_{\text{avg}}^{\text{replication}} \tag{4}$$

Specifically, $BF_{\text{avg}}^{\text{overall}}$ may be interpreted as the overall evidence for association (conditional on the prior assumptions), while $BF_{\text{avg}}^{\text{replication}}$ may be interpreted as the evidence that the effect, as learnt in the discovery data, replicates in the independent replication samples.

In our implementation we use the approximate Bayes factor formulation to compute $BF_{\text{avg}}^{\text{discovery}}$ and $BF_{\text{avg}}^{\text{overall}}$, given the maximum likleihood estimate and standard error computed seperately in discovery and replication samples in each population, as described in Methods. We then use (4) to compute $BF_{\text{avg}}^{\text{replication}}$ as the ratio of the two. However, as described below, we additionally modify this computation to be more lenient about observed differences in effect size between discovery and replication data.

**Allowing for deviation in effect size between discovery and replication**

The method outlined above assumes that true replication effect sizes are identical to those learnt in discovery, and may be too restrictive in practice for several reasons. Firstly, phenotyping may differ between discovery and replication samples. This is the case in our data, where samples with strict phenotype definitions (CM and SMA) were preferentially picked for discovery typing, subject to sufficient DNA quantities. Second, in the context of GWAS, Winner's curse will lead to preferential choice of variants with large observed effect sizes, leading to over-estimation of effect sizes in discovery. Third, the potential for differences in genotyping behaviour between discovery and replication cohorts, e.g. due to technology differences or imputation, may also lead to discrepancies.

To allow for this, we modify formula (2) by additionally allowing the true replication effects to differ from those in discovery. Formally, we split the parameter $\theta$ into a parameter $\theta_1$ (the true effects in discovery) and $\theta_2$ (the true effects in replication), and write

$$\begin{aligned} BF^{\text{overall}} &= \frac{P(D_1|M_1)P(D_2|D_1, M_1)}{P(D_1|M_0)P(D_2|M_0)} \\ &= BF^{\text{discovery}} \cdot \frac{\int_{\theta_2} P(D_2|\theta_2)P(\theta_2|D_1)}{P(D_2|M_0)} \\ &= BF^{\text{discovery}} \cdot \int_{\theta_2} \left( \frac{P(D_2|\theta_2)}{P(D_2|M_0)} \cdot \int_{\theta_1} P(\theta_2|\theta_1) d_1(\theta_1) \right) \end{aligned} \tag{5}$$

For replication analysis, we further assume that $\theta_1$ and $\theta_2$ have a prior joint multivariate normal distribution with zero mean and variance of the form

$$\begin{pmatrix} \Sigma & \rho\Sigma \\ \rho\Sigma & \Sigma \end{pmatrix}$$

where $\Sigma$ reflects the association model for discovery phase, as described in Methods, and $\rho$ is a correlation coefficient. This implies that

$$P(\theta_2|\theta_1) = \mathcal{MVN}\left(\rho\theta_1; (1-\rho^2)\Sigma\right)$$

In our approximate framework the posterior distribution of effect sizes given discovery data, $d_1(\theta_1)$, is also multivariate normal, with distribution

$$d_1(\theta_1) = \mathcal{N}(x^*; A) \quad \text{where} \quad A = \left(\Sigma^{-1} + V^{-1}\right)^{-1} \quad \text{and} \quad x = A\left(V^{-1}\hat{\theta_1}\right)$$

Here as above $\hat{\theta}_1$ and $V$ are the effect size and covariance estimated in discovery data.

To assess the implications of choosing different value of $\rho$ for inference, we compute the joint distribution of parameters given discovery data. This is

$$P(\theta_1, \theta_2 | D_1) = \mathcal{MVN}(y; B)$$

where $B = \left( \begin{pmatrix} \Sigma & \rho\Sigma \\ \rho\Sigma & \Sigma \end{pmatrix}^{-1} + \begin{pmatrix} V^{-1} & 0 \\ 0 & 0 \end{pmatrix} \right)^{-1}$ and $y = B \cdot \begin{pmatrix} V^{-1}\hat{\theta}_1 \\ 0 \end{pmatrix}$. The matrix $B$ can be solved using block inversion, giving

$$\begin{aligned} B &= \begin{pmatrix} \frac{1}{1-\rho^2}\Sigma^{-1} + V^{-1} & -\frac{\rho}{1-\rho^2}\Sigma^{-1} \\ -\frac{\rho}{1-\rho^2}\Sigma^{-1} & \frac{1}{1-\rho^2}\Sigma^{-1} \end{pmatrix}^{-1} \\ &= \begin{pmatrix} A & \rho A \\ \rho A & (1-\rho^2)\Sigma + \rho^2 A \end{pmatrix} \end{aligned}$$

Thus, the marginal distribution on the replication parameters $\theta_2$, given the discovery data $D_1$, is

$$\theta_2 | D_1 \sim \mathcal{MVN}\left( \rho A \cdot V^{-1}\hat{\theta}_1; (1-\rho^2)\Sigma + \rho^2 \cdot A \right)$$

Note that when $\rho = 0$ (no assumed correlation between discovery and replication effect sizes), this says that $\theta_2|D_1$ is distributed according to the prior effect size distribution, while when $\rho = 1$, $\theta_2|D_1$ reduces to the expression for $\theta_1|D_1$, as expected if these parameters are perfectly correlated. In the one-dimensional case, writing $\sigma$ and $v$ for the corresponding variances, the expression becomes

$$\theta_2 | D_1 \sim \mathcal{N}\left( \frac{\rho\sigma\hat{\theta}_1}{v+\sigma}; (1-\rho^2)\sigma + \frac{\rho^2 v\sigma}{v+\sigma} \right)$$

The figure below shows the distribution $\theta_2|D_1$ (y axis, solid and dashed lines) for a fixed observed discovery effect size, a range of values of the discovery standard error $\sqrt{v}$, and four choices of $\rho$. In the main replication analysis presented here we use $\rho = 0.9$.

Figure : Distribution of replication effect size, given an estimated log odds ratio of $\hat{\theta}_1 = 0.396 = \log(1.49)$ observed in discovery (solid horizontal grey line). The discovery standard error is indicated by the x axis. Solid / dashed coloured lines indicate the mean and 95% credible interval of the true effect $\theta_2$ in replication. A marginal prior distribution with mean 0 and standard deviation 0.2 is assumed for discovery and replication effects, with correlation given by $\rho$.

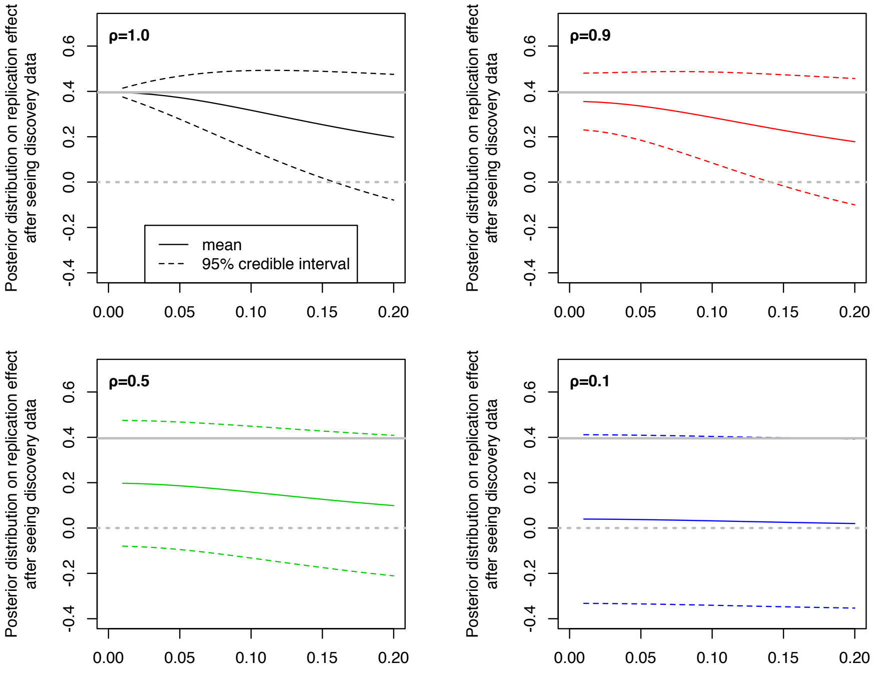

### Supplementary Text 6 - Genome-wide implementation of multinomial logistic regression

##### Introduction

We consider the problem of fitting a regression model for each of a large number of SNPs against a single categorical phenotype. The method described here is implemented in the software SNPTEST (http://www.well.ox.ac.uk/~gav/snptest).

##### Multinomial logistic regression for association testing

Let $\bar{Y} = (Y_i)_{i=1}^N$ denote a set of measurements of a categorical outcome variable on $N$ samples. Each $Y_i$ is assumed to take on one of $M+1$ possible values labelled $0, 1, \cdots, M$. Also, let $\bar{X} = (X_i)$ denote a set of measured covariates for each sample and $\bar{G} = (G_i)$ a predictor of interest. In the context of our study, $Y_i$ is the severe malaria phenotype of sample $i$, with levels 0=Control, 1 = Cerebral malaria, 2 = Severe malarial anaemia, 3 = Other severe malaria, and $G_i$ denotes the genotype of sample $i$ at the variant under consideration.

Multinomial logistic regression models the log-odds of outcome level $i$, relative to the baseline level 0, as a linear combination of predictor and covariates. For a single sample this can be written as

$$\text{logodds}\,(Y = i | G = g, X = x, \theta) = z(g, x) \cdot \theta_i$$

or in terms of outcome probabilities as

$$P(Y = i | G = g, X = x, \theta) = \frac{e^{z(g,x)\cdot\theta_i}}{1 + \sum_{j=1}^{J} e^{z(g,x)\cdot\theta_j}} \tag{1}$$

where

1. $z(g, x)$ denotes the row vector $\left[\begin{array}{ccc} 1 & g & x \end{array}\right]$, consisting of a single 1, the genotype $g$, and the row vector $X$ of measured covariates. More generally we will write $z(g, x) = \left[\begin{array}{ccc} 1 & F(g) & x \end{array}\right]$ where $F(g)$ is a function of the predictor used to model nonadditive effects, as described below.

2. $\theta_j$ denotes a column vector of parameters for outcome $j > 0$. (We always treat $j = 0$ as the baseline outcome, which in the above corresponds assuming that $\theta_0 \equiv 0$ .)

The term $F(g)$ in our study is used to encode different models of effect, as follows. We assume variants are biallelic and let $a(g)$ and $b(g)$ be the counts of the first and second alleles carried by genotype $g$. The table below specifies how different common models of association are encoded in this scheme.

| **Model** | **Encoding** |
|---|---|
| Additive | $F(g) = b(g)$ |
| Dominant | $F(g) = \begin{cases} 1 & \text{if } b(g) > 0 \\ 0 & \text{otherwise} \end{cases}$ |
| Recessive | $F(g) = \begin{cases} 1 & \text{if } b(g) = 2 \\ 0 & \text{otherwise} \end{cases}$ |
| Heterozygote | $F(g) = \begin{cases} 1 & \text{if } b(g) = 1 \\ 0 & \text{otherwise} \end{cases}$ |
| General | $F(g) = \begin{cases} \left[\begin{array}{cc} 0 & 0 \end{array}\right] & \text{if } b(g) = 0 \\ \left[\begin{array}{cc} 1 & 1 \end{array}\right] & \text{if } b(g) = 1 \\ \left[\begin{array}{cc} 2 & 0 \end{array}\right] & \text{if } b(g) = 2 \end{cases}$ |
| Null model | $F(g)$ empty, i.e. no genotype predictor |

**Multinomial regression for directly-typed genotypes**

For $N$ samples, the full likelihood can be written as

$$\begin{aligned} P(\bar{Y}|\bar{G},\bar{X},\theta) &= \prod_{n=1}^{N} P(Y=Y_n|G=G_n,X_n,\theta) \\ &= \prod_{n} f(Y_n;G_n,X_n,\theta) \end{aligned}$$

up to a multiplicative constant, where

$$f(y;g,x,\theta) = \frac{e^{z(g,x)\cdot\theta_i}}{1+\sum_{j=1}^{J} e^{z(g,x)\cdot\theta_j}} \tag{2}$$

The log-likelihood is then

$$\ell(\theta) = \log P(\bar{Y}|\bar{G},\bar{X},\theta) = \sum_{n=1}^{N} \log f(Y_n;G_n,X_n,\theta) \tag{3}$$

For each SNP, we first fit model (3) iteratively starting from $\theta^{\text{null}} \equiv 0$. We then fit the full model starting from the parameters fit under the null model. We use Newton-Raphson iterations to fit both models. This requires computing the first and second derivatives of (3), as described below.

A key assumption underlying the product in (3) is that the outcome for each sample, given its covariates and genotype, is independent of all the other data, i.e.

$$P(Y=Y_n|\bar{Y}_{-n},G,\bar{I},\bar{X},\theta) = f(Y_n;G_n,X_n,\theta) \tag{4}$$

For this to be reasonable in practice, this implies that the covariates $X$ must capture relevant confounding effects, such as environmental effects on the phenotype that are shared between samples. In practice we use principal components computed from genome-wide genotypes as covariates, thus capturing geographic and population structure.

**Multinomial regression for imputed genotypes**

We now consider the case where the predictors are not directly observed but are probabalistically inferred from other, observed quantities. In the GWAS context this corresponds to the situation where genotypes at a variant are imputed from surrounding SNPs, giving a probability distribution over genotypes.

Write $p$ for the function giving the distribution of genotypes at the untyped SNP across the $N$ samples, given the other quantities,

$$p(g_1,\cdots,g_N) = P\left(G=g_1,\cdots,g_N|I,X,\theta,s\right)$$

Here $I$ is used to denote the directly observed genotype data i.e. the genotypes at all directly typed SNPs. The symbol $s$ denotes the fact of having been sampled in the study; we omit this from further notation but return to it below. The full likelihood can now be written by summing over the unobserved genotypes,

$$P(Y=\bar{Y}|I,X=\bar{X},\theta) = \sum_{g_1,\cdots,g_N} P(\bar{Y}|G=g_1,\cdots,g_n,\bar{X},\theta)p(g_1\cdots,g_N) \tag{5}$$

We make additional assumptions that make (5) tractible.

First, we assume that the genotypes at the chosen SNP for each sample are independent of all the genotypes and covariates of all other samples. Namely we write

$$p_n(g_n) = P(G_n=g|G_{-n}=g_1,\cdots,g_N,I,X,\theta) = P(G_n=g|I_n,X_n,\theta) \tag{6}$$

This assumption lets us split the likelihood over samples

$$
\begin{aligned}
P(Y = \bar{Y}|I, X = \bar{X}, \theta) &= \sum_{g_1,\cdots,g_N} P(\bar{Y}|G = g_1, \cdots, g_n, \bar{X}, \theta) p(g_1 \cdots, g_N) \\
&= \sum_{g_1,\cdots,g_N} \prod_n f(Y_n; G_n, X_n, \theta) \cdot p_n(g_n) \\
&= \prod_n \sum_g f(Y_n; g, X_n, \theta) \cdot p_n(g)
\end{aligned}
$$

The last row holds because the $n$th term in the product does not involve $g_m$ for any $m \neq n$, allowing us to reverse the order of the sum and product.

We make a further approximation by taking $p_n$ as the probabilty distribution estimated by genotype imputation - i.e. using IMPUTE2 in our study. IMPUTE2 uses a reference panel of known haplotypes to infer genotypes based on surrounding typed SNPs. We note two ways in which this approximation may become inaccurate. First, if the reference panel populations and study populations are not well matched, i.e. if haplotypes are at substantially different frequencies in the panel and study, then imputation is likely to be inaccurate. (We address this in main text by incorporating population-specific haplotypes into our reference panel.) Second, as described above, $p(g)$ is conditional on covariates, on the effect size parameter $\theta$, and on the fact of having been sampled. IMPUTE2 does not take into account these factors. In particular, dropping covariates from the notation, and writing $\kappa_j$ for the frequency of phenotype level $j$ in the study and $K_j$ for its frequency in the study population, we have

$$
\begin{aligned}
P(G = g|\theta, s) &= \sum_{j=0}^{M} P(G = g|Y = j, \theta) \cdot \kappa_j \\
&= \sum_{j=0}^{M} \frac{P(Y = j|G = g, \theta) P(G = g|\theta)}{P(Y = j|\theta)} \kappa_j \qquad (7) \\
&= \left( \sum_{j=0}^{M} f(j; g, \theta) \cdot \frac{\kappa_j}{K_j} \right) \cdot \text{freq}(g) \qquad (8)
\end{aligned}
$$

Where $\text{freq}(g)$ is the frequency of genotype $g$ in the study population. Thus if $\theta$ is substantially nonzero, then differences between study and population phenotype frequencies, e.g. due to upsampling of disease cases, lead to differences in genotype distributions that are not modelled by imputation. In practice we focus attention on well-imputed genotypes, and we expect most GWAS effect sizes to be small so that these inaccuracies are minor.

##### Derivatives of the complete data log-likelihood

To implement Newton-Raphson iterations for the model described above we need to compute the loglikelihood and its first and second derivatives. We first do this for the case when all genotypes are known (3) and then turn to the full model (5).

Write $f_y(\theta) = f(y; g, x, \theta)$, for given outcome level $y$, considered as a function of $\theta$. We suppress $g$ and $x$ from the notation for a moment. Also we write $z = z(g, x)$ and $D_j$ denotes the operation of taking the derivative with respect to the column vector $\theta_j$. With this notation

$$
f_i = \frac{e^{z^t \theta_i}}{1 + \sum_{j=1}^{J} e^{z^t \theta_j}}
$$

By the quotient rule,

$$D_j f_i = \frac{D_j e^{z^t\theta_i}}{1+\sum_{j=1}^J e^{z^t\theta_l}} - \frac{e^{z^t\theta_{\cdot i}}\cdot D_j e^{z^t\theta_j}}{\left(1+\sum_{l=1}^J e^{z^t\theta_l}\right)^2}$$
$$= z^t \cdot \begin{cases} f_i\,(1-f_i) & \text{if } i=j \\ -f_i f_j & \text{if } i\neq j \end{cases}$$
$$= z^t f_i(\psi_{i,j} - f_j) \quad \text{where} \quad \psi_{i,j} = \begin{cases} 1 & \text{if j=i} \\ 0 & \text{otherwise} \end{cases} \tag{9}$$

since $D_j e^{z^t\theta_l} = 0$ if $l \neq j$. (Note that $f_i$ are scalar-valued functions, so the result of each of these expressions is a row vector reflecting the derivative of $f_i$ with respect to the entries of $\theta_j$.)

Next we take second partial derivatives by applying the product rule to the expressions in (9):

$$D_k D_j f_i = z \otimes z^t \cdot (D_k f_i(\psi_{ij} - f_j) - f_i \cdot D_k f_j)$$
$$= z \otimes z^t \cdot f_i\left((\psi_{i,j} - f_j)(\psi_{i,k} - f_k) - f_j(\psi_{j,k} - f_k)\right) \tag{10}$$

Here $z \otimes z^t$ denotes the Kronecker product (i.e. the matrix of all pairwise multiples of elements of $z$).

The above expressions show that, even though each individual has single assigned outcome, it is useful to compute $f_l$ over all possible outcome levels, since these values can be reused when computing the derivatives.

Remark: the log-likelihood and derivatives are now given by

$$\ell = \sum_n \log f(Y_n; G_n, X_n, \theta) = \sum_n \log f \tag{11}$$
$$D\ell = \sum_n \frac{Df}{f} \tag{12}$$
$$D^2\ell = \sum_n \left(\frac{D^2 f}{f} - \frac{Df^{\otimes 2}}{f^2}\right) \tag{13}$$

The derivative over all parameters can be computed blockwise for given outcome levels using (9) and (10).

##### Derivatives of the full log-likelihood

Now consider the full likelihood (5), i.e. the likelihood allowing for imputed predictor variables. The log-likelihood is

$$\ell^{\text{full}}(\theta) = \sum_n h_n \quad \text{where} \quad h_n = \log \sum_g f(Y_n; g, X_n, \theta) \cdot p_n(g)$$

Consider one term of the outer sum, say the $n-$th term $h = h_n$. Similar to what we did before, write $f_{i,g}(\theta) = f(i; g, X_n, \theta)$ where $i$ is an outcome level. Then

$$h = \log \sum_g f_{Y_n,g} \cdot p_n(g)$$

$$D_j h = \frac{\sum_g D_j f_{Y_n,g} \cdot p_n(g)}{\sum_g f_{Y_n,g} \cdot p_n(g)} \tag{14}$$

(Here we have used the simplifying assumption that $p_n(g)$, computed from imputation, does not depend on $\theta$, and hence does not affect the derivative).

The second derivative is

$$
\begin{aligned}
D_k D_j h &= D_k \left( \frac{\sum_g D_j f_{Y_n,g} \cdot p_n(g)}{\sum_g f_{Y_n,g} \cdot p_n(g)} \right) \\
&= \frac{\sum_g D_k D_j Y_n \cdot p_n(g)}{\sum_g Y_n \cdot p_n(g)} - \frac{\left(\sum_g D_j f_{Y_n,g} \cdot p_n(g)\right) \otimes \left(\sum_g D_k f_{Y_n,g} \cdot p_n(g)\right)}{\left(\sum_g f_{Y_n,g} \cdot p_n(g)\right)^2} \\
&= \left( \frac{\sum_g D_k D_j f_{Y_n,g} \cdot p_n(g)}{\sum_g f_{Y_n,g} \cdot p_n(g)} \right) - (D_j h \otimes D_k h) \qquad (15)
\end{aligned}
$$

**Implementation**

In our implementation, we rely on a linear algebra library (Eigen) which deals with vectors and matrices. To simplify this we first collect the parameters in a single column vector, as

$$
\theta = \begin{bmatrix} \theta_1^t & \theta_2^t & \ldots & \theta_M^t \end{bmatrix}^t
$$

With this convention, $D\ell$ becomes a $1 \times dM$ row vector, and the second derivative $D^2\ell$ is represented as a $dM \times dM$ matrix.

The expressions for the loglikelihood and its derivatives share common terms. We take advantage of these to reduce computation. Define coefficients $A, B, C$ as

$$
A_{n,g} = \frac{f_{Y_n,g} \cdot p_n(g)}{\sum_h f_{Y_n,h} \cdot p_n(h)}
$$

and

$$
B_{n,g,j} = A_{n,g} \cdot (\psi_{Y_n,j} - f_{j,g})
$$

and

$$
C_{n,g,j,k} = A_{n,g} \cdot ((\psi_{Y_n,k} - f_{k,g})(\psi_{Y_n,j} - f_{j,g}) - f_{j,g}(\psi_{j,k} - f_{k,g}))
$$

Then by the above:

$$
D_j \ell = \sum_g \sum_n \left( z_n(g)^t \cdot B_{n,g,j} \right) = \sum_g \mathrm{diag}(B_{\cdot,g,j}) \times Z(g) \qquad (16)
$$

where $\mathrm{diag}(B_{\cdot,g,j})$ denotes the diagonal matrix with $n$th diagonal entry equal to $B_{n,g,j}$. This can be implemented by matrix multiplication. Similarly

$$
D_k D_j \ell = \sum_g \sum_n \left( z_n(g) \otimes z_n(g)^t \cdot C_{n,g,j,k} \right) - \sum_n \left( (D_j \ell)^t \otimes (D_k \ell) \right) \qquad (17)
$$

We follow this implementation strategy, first computing $f_{i,g}$ for each sample and outcome level, using this to compute the loglikelihood and $A_{n,g}$. We then compute $B_{n,g,j}$ and $C_{n,g,j,k}$ and use these to compute the first and second derivative.

### Supplementary Text 7 - Multidimensional inverse variance-weighted meta-analysis

Let $\hat{\beta}_i$ be a parameter estimate from logistic regression in population $i$, and let $V_i$ denote the estimated variance-covariance matrix of $\hat{\beta}_i$. We compute the meta-analysis estimate $\hat{\beta}_{\text{meta}}$ as

$$V_{\text{meta}} = \left(\sum_i V_i^{-1}\right)^{-1} \quad \text{and} \quad \hat{\beta}_{\text{meta}} = V_{\text{meta}} \cdot \left(\sum_i V_i^{-1}\hat{\beta}_i\right) \tag{1}$$

We note that $\hat{\beta}_{\text{meta}}$ is also equal to the combined maximum likelihood estimate, under the assumption that the likelihood function in each population is gaussian with the given mean $\hat{\beta}_i$ and covariance $V_i$ and all studies are independent. In the one-dimensional case, writing lower-case letters for the scalar quantities instead of matrices, (1) reduces to

$$v_{\text{meta}} = \frac{1}{\sum_i \frac{1}{v_i}} \quad \text{and} \quad \hat{\beta}_{\text{meta}} = v_{\text{meta}} \cdot \sum_i \frac{1}{v_i}\hat{\beta}_i$$

A two-tailed p-value for $\beta$ can thus be computed by performing a Wald test, comparing $\hat{\beta}_{\text{meta}}$ to the normal distribution with mean 0 and variance $v_{\text{meta}}$.

In the general case, suppose $\hat{\beta}$ is $d$-dimensional (e.g. representing effects on $d$ subphenotypes). A p-value for each component of $\hat{\beta}_{\text{meta}}$ can be obtained by a Wald test using the corresponding diagonal entry of $V_{\text{meta}}$. To obtain a global p-value against the null that all parameters are zero, let $V_{\text{meta}} = LL^t$ be the Cholesky decomposition of $V_{\text{meta}}$ and $a = L^{-1}\hat{\beta}_{\text{meta}}$. Then

$$\text{var}(a) = \text{Id}$$

We can therefore compute a p-value by computing the sum of squared entries,

$$\zeta = \sum_{k=1}^{d} a_k^2 \quad \text{such that} \quad \zeta \sim \chi_d^2$$

and computing a p-value from quantiles of the $\chi^2$ distribution.

In the interpretation of p-values as the probability of obtaining an estimate "as extreme, or more extreme" than the observed estimate, under the null model, we note that this treats points as "more extreme" when they have lower probability in the full likelihood $\mathcal{N}(0; V_{\text{meta}})$.

In practice the Cholesky decomposition does not need to be computed directly, because

$$\begin{aligned}
\zeta = a^t a &= \hat{\beta}_{\text{meta}}^t L^{t-1} L^{-1} \hat{\beta}_{\text{meta}} \\
&= \left(\sum_i V_i^{-1}\hat{\beta}_i\right)^t V_{\text{meta}} \cdot V_{\text{meta}}^{-1} \cdot V_{\text{meta}} \left(\sum_i V_i^{-1}\hat{\beta}_i\right)^t \\
&= \left(\sum_i V_i^{-1}\hat{\beta}_i\right) \cdot \hat{\beta}_{\text{meta}}
\end{aligned}$$

Thus, the terms of the sum can be computed from $\hat{\beta}_{\text{meta}}$ and the left hand term, which is already computed as part of the computation (1).

#### Text S8 - Analysis of population differentiation

Main text presents an analysis of allele frequency differentiation between continents and between African populations for associated variants (**Fig 7a**). We note here further details of this analysis.

Our between-continent analysis highlights an essential difficulty in making between-continent comparisons of this type for individual variants, namely that it is difficult for protective alleles below around 20-30% frequency to achieve extreme ranks. This is true even for strong resistance alleles that are essentially only present in Africa (e.g. HbS, $rank_{EUR|AFR} = 0.18$; DUP4, $rank_{EUR|AFR} = 0.44$), and presumably reflects the fact that a large number of low-frequency alleles were lost during ancestral bottlenecks in the history of non-African populations. This suggests that, while the observation of elevated frequencies in Africa for particular mutations might imply the action of selection due to *P.falciparum* malaria, as is frequently suggested [16,41,42], it might be equally consistent with neutral evolution under the high levels of genetic drift experienced by historical populations.

In main text we noted enrichment for high levels of within-Africa differentiation among variants with the highest evidence for association (Main text and figure **7b-c**). We note here specific alleles with evidence for differentiation. Most prominent among these is the glycophorin variant DUP4 ($P_{XtX} = 4 \times 10^{-5}$), which as reported previously[7] is only present at high frequency in east African populations (maximum observed frequency < 0.2% in populations west of Cameroon; frequency = 3.7%, 4.9% and 8.9% in Malawi, Tanzania and Kenya respectively). Another associated variant, rs12150480 in *AP2B1*, is of interest since it appears differentiated both within Africa and between continents ($BF_{avg} = 3{,}521$; $P_{XXt} = 5 \times 10^{-3}$ $rank_{EUR|AFR} = 0.036$). This reflects the fact that the protective allele, which also appears to be ancestral, is at much lower frequency in European and Asian populations (84-88%) than in Africa (~98%). Notably, however, rs12150480 is one of a number of variants that show considerable heterogeneity both in frequency and in estimated effect size across populations (e.g. identified as those with maximum Bayes factor ($BF_{max}$) > 25,000 and 100 times greater than the fixed-effect Bayes factor; **Table S4; Fig S10**). The possibility of geographically localized effects, or for locus-specific confounding driven by gradients in selection pressure, cannot be discounted. Fully unraveling such signals is likely to require amalgating data at finer geographic scales[43,44].

References

1. Malaria Genomic Epidemiology, N., Band, G., Rockett, K.A., Spencer, C.C. & Kwiatkowski, D.P. A novel locus of resistance to severe malaria in a region of ancient balancing selection. *Nature* **526**, 253-7 (2015).
2. MalariaGEN. Reappraisal of known malaria resistance loci in a large multi-centre study. *Nature Genetics* (2014).
3. Shah, S.S. *et al.* Heterogeneous alleles comprising G6PD deficiency trait in West Africa exert contrasting effects on two major clinical presentations of severe malaria. *Malar J* **15**, 13 (2016).
4. Clarke, G.M. *et al.* Characterisation of the opposing effects of G6PD deficiency on cerebral malaria and severe malarial anaemia. *Elife* **6**(2017).
5. Jallow, M. *et al.* Genome-wide and fine-resolution association analysis of malaria in West Africa. *Nat Genet* **41**, 657-65 (2009).
6. Band, G. *et al.* Imputation-based meta-analysis of severe malaria in three African populations. *PLoS Genet* **9**, e1003509 (2013).
7. Leffler, E.M. *et al.* Resistance to malaria through structural variation of red blood cell invasion receptors. *Science* **356**(2017).
8. Toure, O. *et al.* Candidate polymorphisms and severe malaria in a Malian population. *PLoS One* **7**, e43987 (2012).
9. Olaniyan, S.A. *et al.* Tumour necrosis factor alpha promoter polymorphism, TNF-238 is associated with severe clinical outcome of falciparum malaria in Ibadan southwest Nigeria. *Acta Trop* **161**, 62-7 (2016).
10. Apinjoh, T.O. *et al.* Association of cytokine and Toll-like receptor gene polymorphisms with severe malaria in three regions of Cameroon. *PLoS One* **8**, e81071 (2013).
11. Apinjoh, T.O. *et al.* Association of candidate gene polymorphisms and TGF-beta/IL-10 levels with malaria in three regions of Cameroon: a case-control study. *Malar J* **13**, 236 (2014).
12. Manjurano, A. *et al.* Candidate human genetic polymorphisms and severe malaria in a Tanzanian population. *PLoS One* **7**, e47463 (2012).
13. Manjurano, A. *et al.* USP38, FREM3, SDC1, DDC, and LOC727982 Gene Polymorphisms and Differential Susceptibility to Severe Malaria in Tanzania. *J Infect Dis* (2015).
14. Manjurano, A. *et al.* African glucose-6-phosphate dehydrogenase alleles associated with protection from severe malaria in heterozygous females in Tanzania. *PLoS Genet* **11**, e1004960 (2015).
15. Ravenhall, M. *et al.* Novel genetic polymorphisms associated with severe malaria and under selective pressure in North-eastern Tanzania. *PLoS Genet* **14**, e1007172 (2018).
16. Opi, D.H. *et al.* Two complement receptor one alleles have opposing associations with cerebral malaria and interact with alpha(+)thalassaemia. *Elife* **7**(2018).
17. Ndila, C.M. *et al.* Human candidate gene polymorphisms and risk of severe malaria in children in Kilifi, Kenya: a case-control association study. *Lancet Haematol* **5**, e333-e345 (2018).
18. Shah, S.S. *et al.* Genetic determinants of glucose-6-phosphate dehydrogenase activity in Kenya. *BMC Med Genet* **15**, 93 (2014).
19. Uyoga, S. *et al.* Glucose-6-phosphate dehydrogenase deficiency and the risk of malaria and other diseases in children in Kenya: a case-control and a cohort study. *Lancet Haematol* **2**, e437-44 (2015).

20. Dunstan, S.J. *et al.* Variation in human genes encoding adhesion and proinflammatory molecules are associated with severe malaria in the Vietnamese. *Genes Immun* **13**, 503-8 (2012).
21. Manning, L. *et al.* A Toll-like receptor-1 variant and its characteristic cellular phenotype is associated with severe malaria in Papua New Guinean children. *Genes Immun* **17**, 52-9 (2016).
22. Flint, J., Harding, R.M., Boyce, A.J. & Clegg, J.B. The population genetics of the haemoglobinopathies. *Baillieres Clin Haematol* **11**, 1-51 (1998).
23. Higgs, D.R. The molecular basis of alpha-thalassemia. *Cold Spring Harb Perspect Med* **3**, a011718 (2013).
24. Williams, T.N. & Weatherall, D.J. World distribution, population genetics, and health burden of the hemoglobinopathies. *Cold Spring Harb Perspect Med* **2**, a011692 (2012).
25. Lam, K.W. & Jeffreys, A.J. Processes of copy-number change in human DNA: the dynamics of {alpha}-globin gene deletion. *Proc Natl Acad Sci U S A* **103**, 8921-7 (2006).
26. Crosnier, C. *et al.* Basigin is a receptor essential for erythrocyte invasion by Plasmodium falciparum. *Nature* **480**, 534-7 (2011).
27. McLaren, W. *et al.* The Ensembl Variant Effect Predictor. *Genome Biol* **17**, 122 (2016).
28. Consortium, E.P. An integrated encyclopedia of DNA elements in the human genome. *Nature* **489**, 57-74 (2012).
29. Roadmap Epigenomics, C. *et al.* Integrative analysis of 111 reference human epigenomes. *Nature* **518**, 317-30 (2015).
30. Xu, J. *et al.* Combinatorial assembly of developmental stage-specific enhancers controls gene expression programs during human erythropoiesis. *Dev Cell* **23**, 796-811 (2012).
31. GTEx Consortium. Human genomics. The Genotype-Tissue Expression (GTEx) pilot analysis: multitissue gene regulation in humans. *Science* **348**, 648-60 (2015).
32. Lessard, S. *et al.* An erythroid-specific ATP2B4 enhancer mediates red blood cell hydration and malaria susceptibility. *J Clin Invest* **127**, 3065-3074 (2017).
33. Westra, H.J. *et al.* Systematic identification of trans eQTLs as putative drivers of known disease associations. *Nat Genet* **45**, 1238-1243 (2013).
34. MacArthur, J. *et al.* The new NHGRI-EBI Catalog of published genome-wide association studies (GWAS Catalog). *Nucleic Acids Res* **45**, D896-D901 (2017).
35. Wellcome Trust Case Control, C. *et al.* Bayesian refinement of association signals for 14 loci in 3 common diseases. *Nat Genet* **44**, 1294-301 (2012).
36. Alvarez, M.I. *et al.* Human genetic variation in VAC14 regulates Salmonella invasion and typhoid fever through modulation of cholesterol. *Proc Natl Acad Sci U S A* **114**, E7746-E7755 (2017).
37. Dunstan, S.J. *et al.* Variation at HLA-DRB1 is associated with resistance to enteric fever. *Nat Genet* **46**, 1333-6 (2014).
38. Gilchrist, J.J. *et al.* Genetic variation in VAC14 is associated with bacteremia secondary to diverse pathogens in African children. *Proc Natl Acad Sci U S A* **115**, E3601-E3603 (2018).
39. Astle, W.J. *et al.* The Allelic Landscape of Human Blood Cell Trait Variation and Links to Common Complex Disease. *Cell* **167**, 1415-1429 e19 (2016).
40. Lopez-Mejias, R. *et al.* A genome-wide association study suggests the HLA Class II region as the major susceptibility locus for IgA vasculitis. *Sci Rep* **7**, 5088 (2017).
41. Ma, S. *et al.* Common PIEZO1 Allele in African Populations Causes RBC Dehydration and Attenuates Plasmodium Infection. *Cell* **173**, 443-455 e12 (2018).
42. Zhang, D.L. *et al.* Erythrocytic ferroportin reduces intracellular iron accumulation, hemolysis, and malaria risk. *Science* **359**, 1520-1523 (2018).

43. Piel, F.B. *et al.* Global distribution of the sickle cell gene and geographical confirmation of the malaria hypothesis. *Nat Commun* **1**, 104 (2010).
44. Mackinnon, M.J. *et al.* Environmental Correlation Analysis for Genes Associated with Protection against Malaria. *Mol Biol Evol* **33**, 1188-204 (2016).
